# Locus specific human endogenous retroviruses reveal new lymphoma subtypes

**DOI:** 10.1101/2023.06.08.544208

**Authors:** Bhavya Singh, Nicholas Dopkins, Tongyi Fei, Jez L. Marston, Stephanie Michael, Helena Reyes-Gopar, Gislaine Curty, Jonas J. Heymann, Amy Chadburn, Peter Martin, Fabio E. Leal, Ethel Cesarman, Douglas F. Nixon, Matthew L. Bendall

## Abstract

The heterogeneity of cancers are driven by diverse mechanisms underlying oncogenesis such as differential ‘cell-of-origin’ (COO) progenitors, mutagenesis, and viral infections. Classification of B-cell lymphomas have been defined by considering these characteristics. However, the expression and contribution of transposable elements (TEs) to B cell lymphoma oncogenesis or classification have been overlooked. We hypothesized that incorporating TE signatures would increase the resolution of B-cell identity during healthy and malignant conditions. Here, we present the first comprehensive, locus-specific characterization of TE expression in benign germinal center (GC) B-cells, diffuse large B-cell lymphoma (DLBCL), Epstein-Barr virus (EBV)-positive and EBV-negative Burkitt lymphoma (BL), and follicular lymphoma (FL). Our findings demonstrate unique human endogenous retrovirus (HERV) signatures in the GC and lymphoma subtypes whose activity can be used in combination with gene expression to define B-cell lineage in lymphoid malignancies, highlighting the potential of retrotranscriptomic analyses as a tool in lymphoma classification, diagnosis, and the identification of novel treatment groups.

## Introduction

Transposable elements (TEs) account for roughly 45% of the human genome^1, 2^. They include retrotransposons^3–5^, which can be further broken down into short interspersed nuclear elements (SINEs), long interspersed nuclear elements (LINEs), and human endogenous retroviruses (HERVs). HERVs are the remains of ancient retroviral infections that integrated within the germline^6, 7^. Since their integration, HERVs have accumulated mutations and deletions, but some of them have been co-opted by the host and can mediate key physiological processes^8–14^. Under some conditions, the derepression of HERVs can be associated with viral infectivity, pathogenic inflammation, and oncogenesis^15–19^. Regulation of their expression is thought to be a driving factor in the initiation and sustainment of some human diseases^20–22^.

While factors underlying the deregulation of HERV expression remain poorly defined^23^, there is a strong causal relationship with viral infections co-opting HERV expression or derailing their regulatory networks^15, 24^. Transactivation of TEs by cancer-associated viruses such as with Epstein-Barr Virus (EBV) could help drive the heterogenous development of non-Hodgkin B-cell lymphomas^24–30^. This heterogeneity in aggressive B-cell lymphomas may also be driven by other confounding factors, such as translocations events occurring at immunoglobulin, proto-oncogene, and tumor suppressor gene loci, somatic mutations, and often, differential ‘cells-of-origin’ (COO) derived from the germinal center (GC)^31–35^.

Characterizing TE activity has posed unique challenges due to their repetitive nature, poor delineation, non-canonical activity, and low expression^36^. Recent advancements in computational biology now permit more accurate depiction of TE activity by next generation sequencing (NGS) technologies^37–42^. When HERVs are transcribed their products can be collected in RNAseq libraries, and the collective noun for these transcripts, in contrast to the gene derived transcriptome, is called the “retrotranscriptome”. Oncogenic TE-gene chimeric transcripts have been identified in a subset of diffuse large B-cell lymphoma (DLBCL) cases^27^, and HERV dysregulation has been observed in response to EBV^30, 43^ and human immunodeficiency virus-1 (HIV-1)^44–48^ infections, both of which are associated with Burkitt lymphoma (BL) and DLBCL. B-cell lymphomas have been subcategorized by classifiers such as LymphGen^35^ and EcoTyper^49^ to aid in treatment selections, these classifications have not included TE expression. Here, we present a comprehensive, locus-specific analysis of TE expression in germinal Center (GC) B-cells, DLBCL, EBV-positive and negative BL, and follicular lymphoma (FL) to create the first retrotranscriptomic atlas of GC derived non-Hodgkin’s lymphomas. Our results classify lymphomas by locus-specific TE expression and identify additional prognostic categories, with the potential for new approaches to treatments.

## Results

### The retrotranscriptomic landscape of B-cell lymphomas and germinal center B-cells: HERVs distinguish specific B-cell subsets

We obtained RNA-seq data from FACS-sorted B-cell populations from two publicly available datasets^31, 50^. The Agirre et al.^50^ (B-AG) B-cell dataset was comprised of dark zone (DZ), light zone (LZ), naïve B (NB), memory B (MB), plasmablasts (PB), and bone marrow plasma cells (BMPC) from 35 samples, while the Holmes et al.^31^ (B-HM) B-cell dataset was comprised of DZ, LZ, NB, MB, and the whole germinal center (GCB) from 17 samples. RNA-seq reads were aligned to the human genome (hg38) using a splice-aware aligner, STAR. Quantification of gene features in the GENCODE (v38) annotation was performed by STAR, while TE expression of 14,896 HERV and 13,545 LINE elements was quantified with Telescope^51^. As a filtering criterion, we included elements with >5 reads in at least 10% of the samples, leaving 1,464 HERVs and 1,939 LINEs in the B-HM dataset, and 1,118 HERVs and 1,520 LINEs in the B-AG dataset (Supplementary Table 1).

The retrotranscriptome of healthy B cells, including GC cells, were used for comparison with B-cell lymphoma retrotranscripts. In both B-HM and B-AG, NB-cells had the highest percentage of reads assigned to TEs (0.61%, 0.74%), followed by MB-cells in B-HM (0.6%), and by PB (0.88%) and BMPC (0.86%) in B-AG (Fig. 1A-B). In both datasets, DZ had the lowest TE expression (0.41% in B-HM and 0.71% in B-AG). Plasmablasts (PBs) and bone marrow derived plasma cells (BMPCs) had the lowest HERV expression despite having the highest TE expression, indicating that a larger proportion of their TE fragments came from LINE elements (Fig. 1C-D).

**Figure 1:**
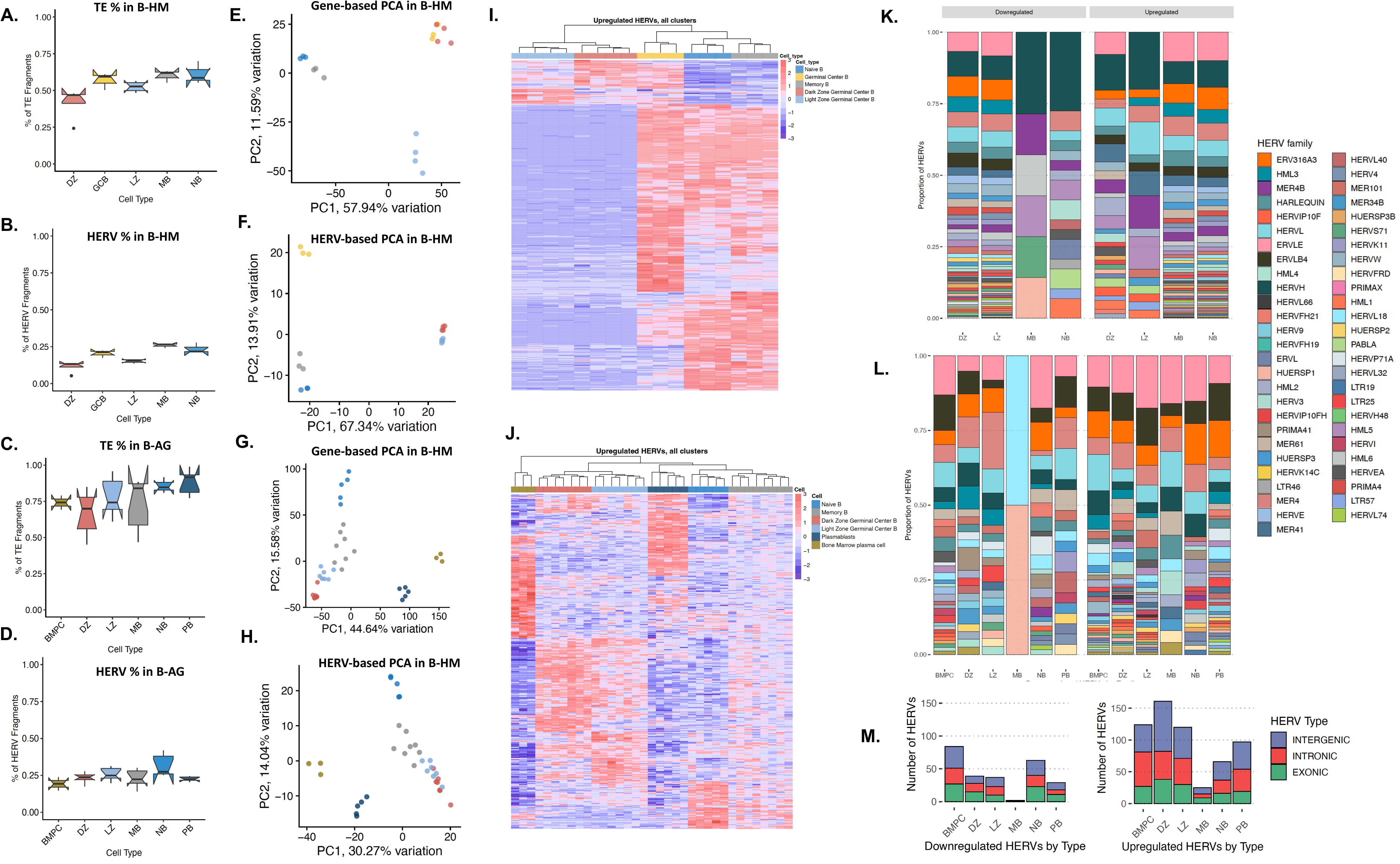
HERVs distinguish specific B cell subsets. **A.** TE reads, and **B.** HERV reads as a percent of all filtered sequencing reads per cell-type in the B-HM dataset. **C.** TE reads, and **D.** HERV reads as a percent of all filtered sequencing reads per cell-type in the B-AG dataset **E.** PCA plot of germinal center B cells from the Holmes dataset (NB, MB, DZ, LZ, and whole GCB), clustered by genes from the hg38 human genome annotation. **F.** PCA plot of germinal center B cells from the Holmes dataset, clustered by HERV expression using the Telescope annotation. HERV expression uniquely distinguishes B cell subsets compared to genes, with HERVs in the light zone and dark zone following similar patterns of expression. **G.** PCA plot of germinal center B cells from the Agirre dataset (NB, MB, DZ, LZ, PB, and BMPB), clustered by genes from the hg38 human genome annotation. **H.** PCA plot of germinal center B cells from the Agirre dataset, clustered by HERV expression using the Telescope annotation**. I.** Heatmap of top upregulated HERVs by cell-type in the Holmes dataset (p-value < 0.001, log2fold change > 1.5). Light zone and dark zone display downregulation of HERVs that are most highly expressed in other cell - types. **J.** Heatmap of top upregulated HERVs by cell-type in the Agirre dataset (p-value < 0.001, log2fold change > 1.5). Light zone and dark zone display downregulation of HERVs that are most highly expressed in other cell-types, with plasmablasts and bone marrow plasma cells displaying the highest number of differentially expressed HERVs. **K-L.** Relative abundance of HERV families upregulated and downregulated per cell-type in the Holmes and Agirre datasets, displaying a high number of loci assigned to ERVLE, HERVH, ERV316A3, HARLEQUIN, ERVLB4,and HERVFH21**. M.** Number of upregulated HERVs in cell-types in the B-AG dataset, colored by the location of HERVs in relation to nearby genes (exonic, intergenic, intronic).

We performed an unsupervised principal component analysis (PCA) to visualize sample placement-based gene or HERV expression in B-cell subpopulations in the B-HM (Fig. 1E, F) and B-AG datasets (Fig. 1G, H). Similar to the gene-driven PCA, the first principal component of a HERV-driven PCA in the B-HM dataset separated the NB and MB-cells from LZ and DZ cells (Fig. 1F). While the second principal component in the gene driven PCA separated the LZ and DZ, the HERV expression in LZ and DZ was comparatively similar, leading to closer clustering. In the B-AG dataset, the first principal component segregated the PB and BMPC from LZ, DZ, MB, and NB, while the second principal component separated the NB, MB, LZ, and DZ (Fig. 1G). Analogous to the B-HM dataset, HERV expression was more similar between LZ and DZ than gene expression (Fig. 1H). The GC B retrotranscriptomic landscape changes throughout B-cell differentiation.

Next, we identified unique sets of significantly differentially expressed (DE) HERVs in each B-cell subtype (Supplementary Table 2). The cell subtypes with the highest number of upregulated HERVs were observed in the BMPC, PB, and DZ subsets in the B-AG dataset (Figure 1I, Supplementary Fig. 1A-F) and in NB, MB, and GCB in the B-HM dataset (Fig. 1J, Supplementary Fig. 2A-F). In both datasets, the most DE loci belonged to the ERVLE, HERVH, ERV316A3, ERVLB4, and MER4 families (Fig. 1K-L). Interestingly, HERVs along the 22q11 locus such as HUERSP3B_22q11.22 and ERVLE_22q11.22b were commonly upregulated in the DZ, suggesting changes in nucleosomal accessibility at this site. HARLEQUIN_1q32.1, which has previously been found to be differentially expressed in prostate, breast, and colon cancers, was downregulated in the DZ and upregulated in the PBs and BMPCs compared to other B-cell subtypes^52^. PBs, which have been hypothesized to be the COO of ABC-DLBCL, displayed upregulation in 3 HERVP71A loci among the top DE-HERVs (Supplementary Fig 3, Supplementary Fig. 4). Collectively, these data suggest significant changes in HERV loci expression can be correlated to B-cell fate within the GC.

### Lymphoma subtypes have distinct HERV expression landscapes

Since HERV expression profiles are unique to tissue sites^8, 53, 54^ and patterns of malignancy, we hypothesized that different B-cell lymphomas would display unique HERV signatures that could be used to further classify malignancy subtypes. BL had the highest percentage of reads assigned to TEs and HERVs (2.27% and 0.65%), followed by FL (0.61% and 0.24%), and DLBCL (0.49% and 0.2%) (Supplementary Fig. 5). By conducting unsupervised clustering via PCA-based metrics, we found that HERVs (Figure 2B) better segregate FL, ABC, EBV+ BL, EBV negative BL, GCB, and unclassified DLBCL cases than genes (Figure 2A). Further characterizing of lymphoma types showed that BL had 2910 uniquely upregulated HERV loci compared to DLBCL and FL, which had 184 and 31, respectively (Fig. 2C-F). Within the lymphoma subtypes, GCB-DLBCL had the highest number of uniquely upregulated HERVs at 511, followed by endemic EBV+ BL at 456 loci, and sporadic EBV negative BL at 409 loci (Supplementary Fig 6A). When accounting for shared upregulated loci, BL exhibited broad upregulation of HERVs across all subtypes when compared to DLBCL and FL (Fig. 2F-H). Similar to benign B-cells, the highest number of differentially expressed loci belonged to the ERVLE, ERV316A3, HERVH, ERVLB4, HERVL, HERVFH21, HML3, and HARLEQUIN families, with the highest upregulation of a HERV family being that of HERVH in GCB-DLBCL (Fig. 2E, Supplementary Fig. 6C). We also observed HERV-based DZ markers being broadly upregulated in BL compared to other lymphoma subtypes, such as MER61_3q13.11, HERV3_14q32.33, and HARLEQUIN_19p12b (Supplementary Fig 4, Supplementary Fig 7). A key HERV marker of PB and BMPCs, HARLEQUIN_1q32.1, was significantly upregulated in a subset of ABC-DLBCLs (p<0.001, Supplementary Fig 8). Collectively, these data demonstrate that HERVs act as novel retrotranscriptomic markers that can be used to discriminate heterogeneity between B-cell malignancies.

**Figure 2:**
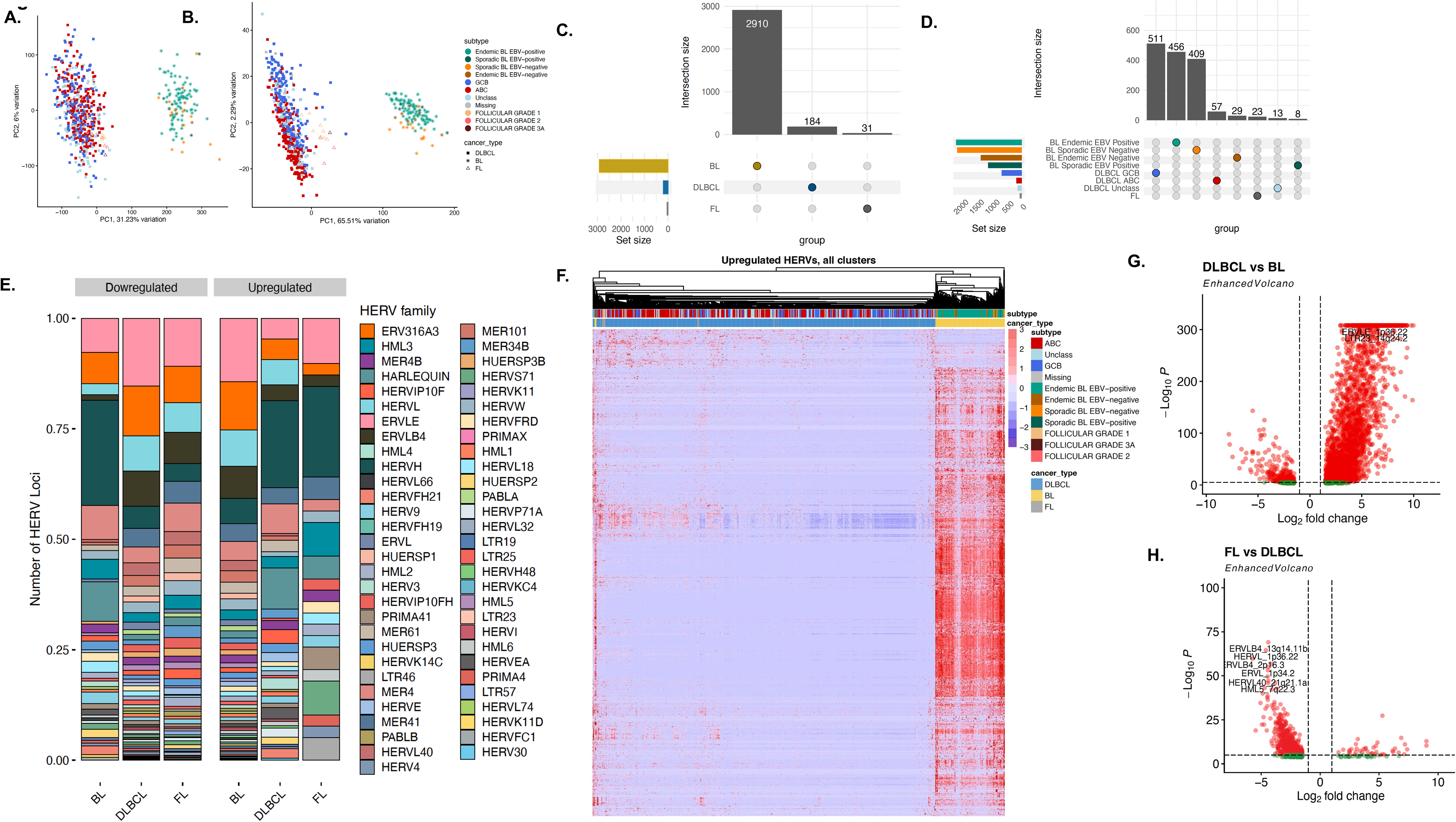
HERV expression is specific to lymphoma subtypes. **A**. PCA plot of 529 DLBCL samples from the TCGA and NCICCR datasets, 113 BL samples from CGCI, and 12 FL samples, clustered by genes from the hg38 human genome annotation. **B.** PCA plot of 529 DLBCL samples from the TCGA and NCICCR datasets, 113 BL samples from CGCI, and 12 FL samples, clustered by HERV expression from the Telescope annotation. **C.** Upset plot of the number of unique and shared HERVs upregulated in each cancer type (p < 0.001, log2fold change > 1.5). Within the three non-Hodgkin’s B cell lymphomas, Burkitt lymphoma displays the highest HERV upregulation. **D.** Upset plot of the number of unique and shared HERVs upregulated in each cancer sub-type, including ABC, GCB, and unclassified DLBCL, sporadic and endemic BL by EBV status, and follicular lymphoma. **E.** Relative abundance of HERV families per lymphoma type, displaying a high number of loci assigned to ERVLE, HERVH, ERV316A3, HERVL, ERVLB4,and HERVFH21 **F.** Heatmap of upregulated HERVs in each lymphoma subtype (p < 0.001, log2fold change > 1.5), showcasing a remarkable upregulation of HERVs in BL compared to DLBCL and FL. **G.** Volcano plot of differentially-expressed HERVs in DLBCL and BL (p-value < 0.001, log2fold change > 1.5. H. Volcano plot of differentially-expressed HERVs in FL and DLBCL (p-value < 0.001, log2fold change > 1.5.

### A subset of HERV features differentiate lymphoma subtypes and GC-B COO

We next asked whether the HERV-driven B-cell malignancy signatures could complement gene expression data to best define the GC COO. Our goal was to reduce the large number of DE HERV features to the lowest possible targets for reliable classification. Including only DE HERVS with an FDR <0.001 and log2fold >1.5, we used two unsupervised feature selection methods, 1) the random forest classification with the Boruta algorithm^55^, and 2) the randomized least absolute shrinkage and selection operator (LASSO) regression^56^, identifying just 5 HERVs to differentiate between DLBCL, BL, and FL (Fig 3A). Out of the 5 HERVs, ERVL_1p34.2 expression differentiated between BL and FL, while ERLB4_2p16.3 differentiated between DLBCL, and FL and BL (Fig 3B-3G). We next created feature sets for each B-cell subtype from the B-AG dataset, using the top 150 upregulated genes and top 25 upregulated HERVs for MB, NB, DZ, LZ, PB, and BMPC (Supplementary Table 3). To assign COO, we ran a fast HERV and gene-set enrichment analysis (F-HAGSEA) using an adaptive multilevel split Monte Carlo method^57^. Consistent with known literature^34^, we found that all BL subsets were enriched in DZ signatures, ABC-DLBCL enriched in PB and MB signatures, GCB-DLBCL in LZ, and, interestingly, FL in NB and LZ (Fig 3H). Overall, our findings indicate that HERVs are uniquely expressed in healthy B-cells and lymphoma subtypes, and that HERV expression profiles can be further used in combination with gene expression profiles to best define the COO for B-cell malignancies.

**Figure 3:**
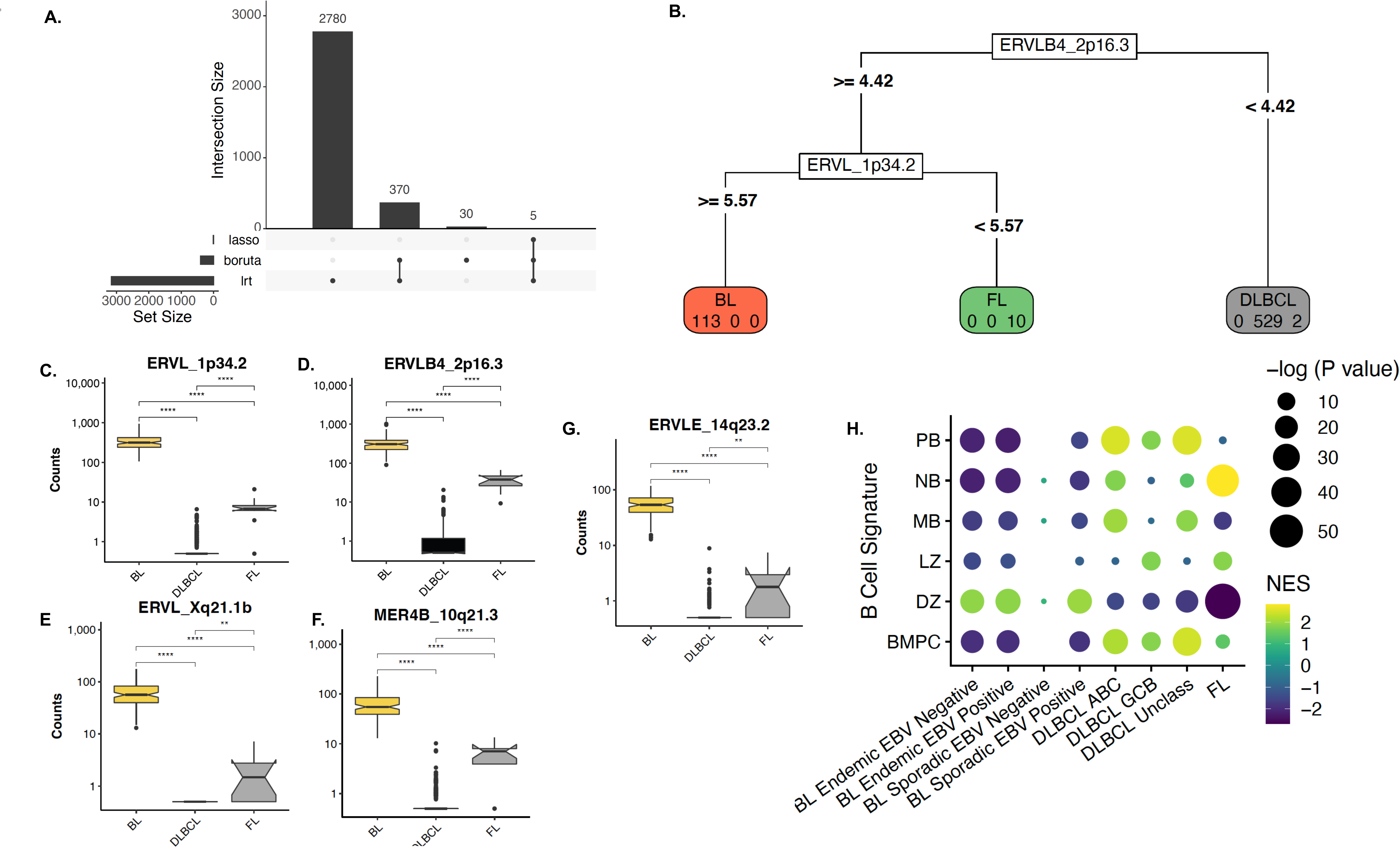
HERV expression aids in identifying lymphoma subtypes and potential GC B COO. **A.** UpsetR plot displaying the number of features selected by DESeq2 lowest likelihood ratio (LTR), the random forest classification with the Boruta algorithm, and the randomized least absolute shrinkage and selection operator (LASSO) regression, with 5 features being selected by all three methods. **B**. A subset of four HERVs can independently categorize lymphoma subtypes, with **C.** ERVL_1p34.2 expression differentiating between BL and FL, and **D.** ERLB4_2p16.3 differentiating between DLBCL, and FL and BL, in addition to **E.** ERVL_Xq21.1b, **F.** MER4B_10q21.3, and **G.** ERVLE_14q23.2. **H.** Correlation plot of lymphoma sub-types with gene and HERV-based B-cell-of-origin signatures. Signature gene sets were created using a subset of the top 150 and top 25 upregulated genes and HERVs per cell-type from the Agirre 2019 B cell dataset.

### Seven distinct HERV signatures categorize diffuse large B-cell lymphoma

Given that ABC-DLBCL and GCB-DLBCL display distinct patterns of HERV expression, we investigated whether subsets within the COO classes possessed unique HERV signatures that could further define their characterization. We performed unsupervised consensus clustering with ConsensusClusterPlus^58^ based on DE HERVs to identify the number of potential subsets, *k*, along with the strength of each sample’s membership in the identified class. While the most stable *k* yielded 3 clusters most consistent with current COO classes, we chose a *k* of 7 to potentially identify sub-classes of HERV signatures within the ABC, GCB, and unclassified DLBCLs (Fig. 4A-C). When comparing HERV clusters (HCs) to the COO subtypes, the ABC-DLBCL were split predominantly into HC1 and HC2, while HC4 and HC6 belonged predominantly to the GCB-DLBCL class. HC3 and HC5 were mixed clusters of all three classifications, while HC7 encompassed ABC-DLBCL and the highest number of unclassified samples (Fig 4D, Supplementary Fig. 9A). When compared to the LymphGen classes, HC2 consisted predominantly of MCD, HC3 contained the highest number of BN2, and HC4 and HC6 encompassed the highest number of EZB. The N1 subclass was split between HC5 and HC7 (Fig. 4E, Supplementary Fig. 9A). HC6 had the highest number of uniquely upregulated HERVs at 1,682 loci, while HC7 had the highest number of uniquely downregulated HERVs, at 202 loci (Fig. 4F-G). Compared to healthy B-cells, loci from the HERVH family represented a higher proportion of upregulated HERVs (Fig. 4H), with HC7 displaying the highest upregulation of HERVH transcripts. Four key HERVs that could differentiate the DLBCL clusters (Supplementary Fig. 10A) were HERVH_16p13.2e, HERVW_2q23.3, HML2_7p22.1, and HERVH_7q11.23a (Supplementary Fig. 10B-E). HERVH_16p13.2e differentiates HC7 from the remaining clusters, while HERVH_16p13.2e differentiates HC1 and HC2. HML2_7p22.1 separates HC4 and HC6 from HC3, HC4, and HC7, and then further differentiates within the clusters.

**Figure 4:**
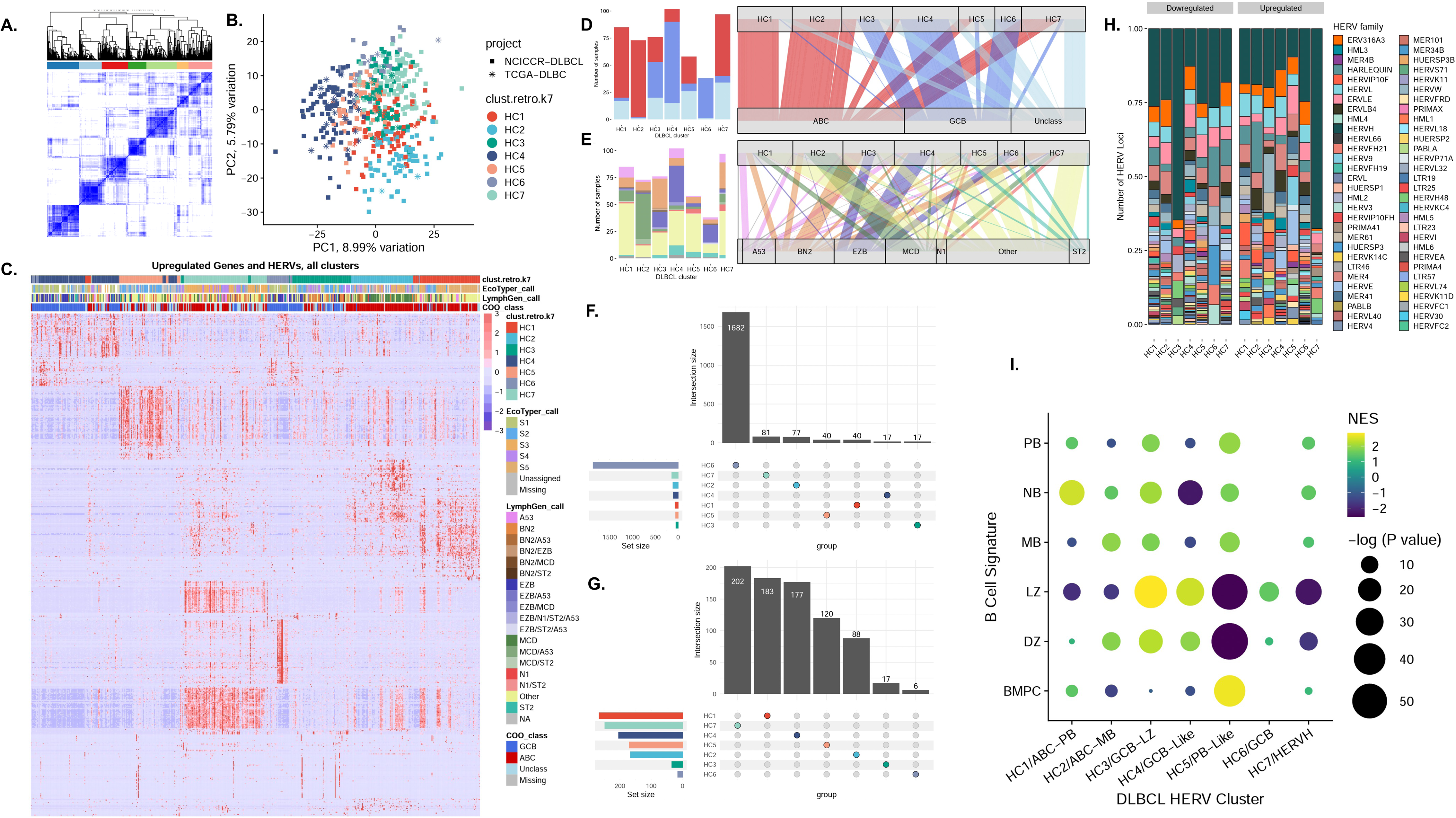
Seven distinct HERV signatures in diffuse large B-cell lymphoma. **A.** Consensus clustering of TCGA and NCICCR DLBCL samples find seven distinct sample clusters, based on expression values of the top 10% of most variable HERVs. **B.** PCA of DLBCL samples, colored by HERV clusters. **C.** Alluvial diagram showcasing HERV cluster assignment in comparison with recent DLBCL classification paradigms, including the LymphGen, EcoTyper, DBL Hit presence, and classic cell-of-origin classifications. When comparing HERV clusters to the COO subtypes, HC1 and HC2 belong predominantly to the ABC-DLBCL class, while HC4 and HC6 belong predominantly to the GCB-DLBCL class. HC3 and HC5 are mixed clusters of all three classifications, while HC7 encompasses ABC-DLBCL, with the highest number of unclassified samples. **D.** When comparing HERV clusters to the LymphGen classes, HC2 consists predominantly of MCD, HC3 consists of the highest number of BN2, and HC4 and HC6 encompass the highest number of EZB. The N1 subclass is split between HC5 and HC7. **E.** Heatmap of the top 50 upregulated genes and HERVs per DLBCL cluster (p < 0.001, log2fold change > 1.5). **F**. Upset plot of the uniquely upregulated HERVS per cluster, and **G.** Upset plot of the uniquely downregulated HERVs per cluster, finding the highest number of unique genes in C6. **H.** Relative abundance of HERV families per DLBCL type**. I.** Gene and HERV-driven B-cell-of-origin classification of each HERV-driven DLBCL cluster. Signature gene sets were created using a subset of the top 150 and top 25 upregulated genes and HERVs per cell-type from the Agirre 2019 B cell dataset. HC1 and HC2, which belong predominantly to the ABC-DLBCL subclass, are enriched in NB and PB, and MB and DZ gene-sets respectively. HC3, which is a mixed subtype, is most enriched in LZ signatures. HC4 and HC6, which are both predominantly GCB-DLBCLs, are enriched in LZ signatures. HC5 and HC7, which are mixed subtypes containing ABC-DLBCL and unclassified samples, are most enriched for BMPC signatures, with negative enrichment scores for both LZ and DZ.

To determine potential GC-B COO for the seven DLBCL subsets, we conducted an F-HAGSEA analysis against the B-cell signatures, using feature ranks derived from DESEq2 differential testing^59^ (Fig 4I). HC1 and HC2 were most enriched in NB and PB, and MB and DZ gene-sets respectively. HC3, which is a mixed subtype, was most enriched in LZ signatures. HC4 and HC6, which are both predominantly GCB-DLBCLs, were also enriched in LZ signatures. HC5 and HC7 were most enriched for BMPC signatures, with negative enrichment scores for both LZ and DZ. We thus designated HC1 and HC2 with the names “ABC-PB” and “ABC-MB” (Supplementary Fig. 11), HC3, HC4 and HC6 with the names “GCB-LZ”, “GCB-Like”, and “GCB” (Supplementary Fig. 12), HC5 with the name “PB-Like”, and HC7 with “HERVH” (Supplementary Fig. 13). Overall, our results identified 7 distinct HERV signatures in DLBCL samples which identify novel subclasses of the currently implemented DLBCL COO classifications.

### Two distinct HERV signatures are found in Burkitt lymphoma that are indicative of EBV status

Since HERVs are transactivated by EBV^30^, we hypothesized that heterogenous HERV expression profiles in BL are driven by infection with EBV. We performed unsupervised PCA clustering of pediatric BL samples based on gene expression (Fig. 5A) and HERV expression (Fig. 5B) alone. Surprisingly, we found that while the gene-based PCA did not segregate samples by EBV status, HERV expression separated BL status into EBV+ and EBV-clusters. To confirm the results of the PCA, we performed consensus clustering of samples based on HERV expression, finding the most stable clusters with a *k* of 2 (Fig. 5C-D). The BL cluster 1 (BL-C1) was composed entirely of EBV- samples (13 EBV-endemic BL samples and 3 EBV-sporadic BL samples) while BL cluster 2 (BL-C2) was composed of primarily EBV positive samples (4 EBV- endemic BL samples, 4 EBV+ sporadic BL, and 89 EBV+ endemic BL). Collectively, these separations were driven by an overall upregulation of TEs in BL-C2 (Fig. 5E-G), with 253 uniquely upregulated HERVs in BL-C2, compared to 66 in BL-C1 (Supplementary Fig. 14A). We next sought to identify the HERV signatures driving separation of BL-C1 and BL-C2 with the Boruta algorithm, LASSO regression, and the likelihood ratio test (LRT) provided by DESEQ2. In doing such, we identified a subset of four HERVs that further distinguished between the BL-C1 and BL-C2 (Fig 6A). Amongst all HERVs, we identified ERVLE_2p25.3c (Fig 6B), MER61_4p16.3 (Fig 6C), ERV316A3_2q21.2b (Fig 6D), and ERVLE_5p13.2c (Fig. 6E) as definitive markers that distinguished between the entirely EBV-BL-C1, and the largely EBV+ BL-C2 (Supplementary Fig 15). We further identified BL-C1 to have a more distinct DZ signature compared to BL-C2, and additionally found a higher relative upregulation of Hallmark pathways identified by the Molecular Signatures Database (MsigDB)^60, 61^ when compared to BL-C2 (Fig 6F-G). Collectively, these results demonstrate that EBV status is a major determinant of HERV expression in BL subtypes, and that the expression of HERVs can be applied to better define the heterogeneity of pediatric BL.

**Figure 5:**
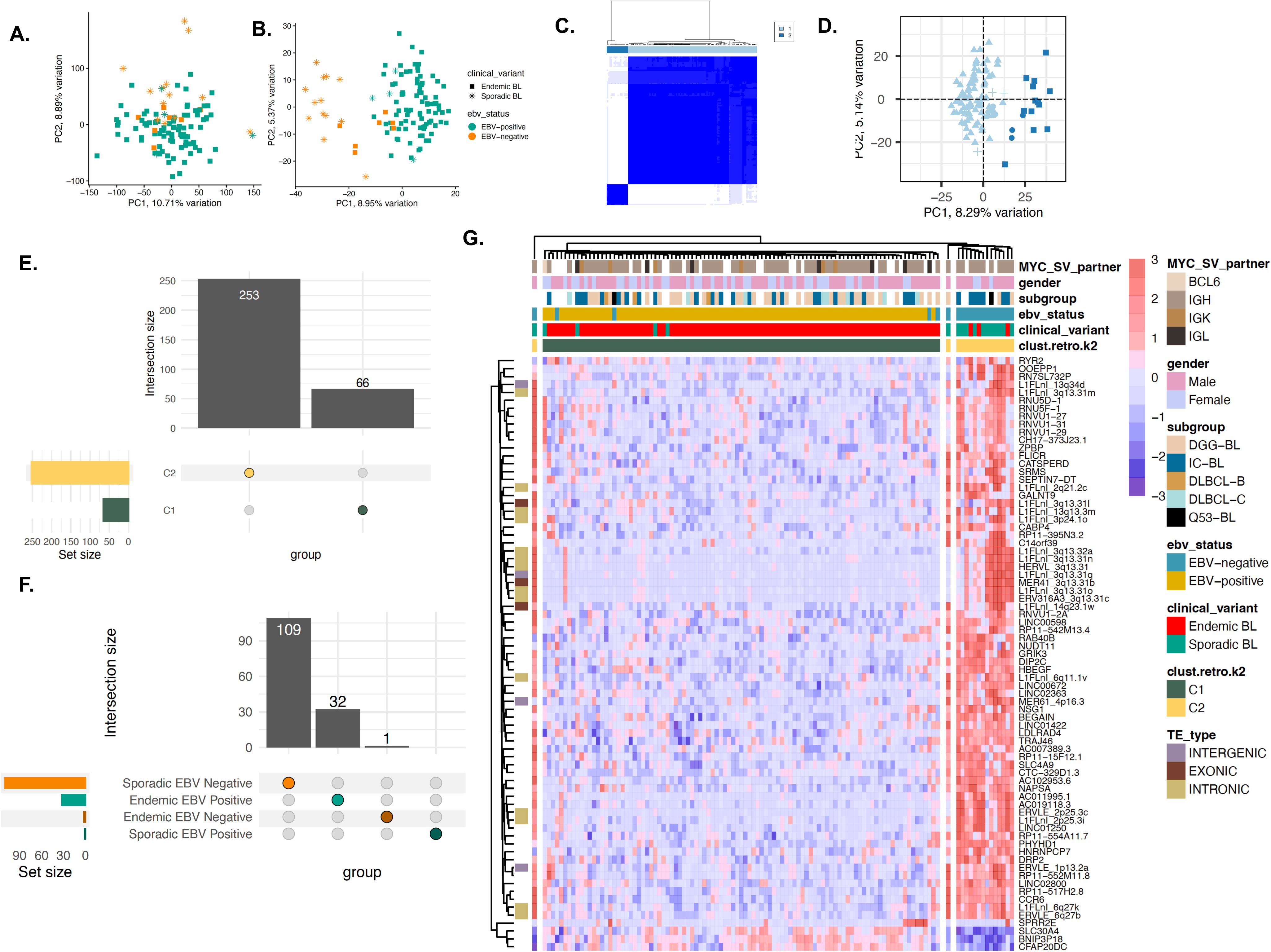
Two distinct HERV signatures are found in Burkitt lymphoma independent of EBV status. **A.** PCA plot of 113 BL samples from CGCI datasets, 113 BL samples from CGCI, clustered by genes from the hg38 human genome annotation. **B.** PCA plot of BL samples, clustered by HERV expression from the Telescope annotation. HERV-only clustering reliably separates the EBV-positive and EBV-negative samples, showcasing distinct expression patterns in the HERVs that are not captured with gene-only clustering. **C-D**. Consensus clustering of BL samples find two distinct sample clusters, with BL-C1 containing all EBV-positive endemic and sporadic BL samples, along with three EBV-negative endemic BL samples. BL-C2 consists of all EBV negative sporadic BL samples, along with three EBV-negative endemic BL samples **E.** BL-C2, which predominantly contains EBV negative sporadic BL samples, contains 253 uniquely upregulated HERVs, compared to 66 in BL-C1**. F.** When comparing within subtypes, EBV-sporadic BL has the most number of uniquely upregulated HERVs, followed by EBV+ endemic BL. **G.** Heatmap of the top 50 upregulated genes and HERVs per DLBCL cluster (p < 0.001, log2fold change > 1.5).

**Figure 6:**
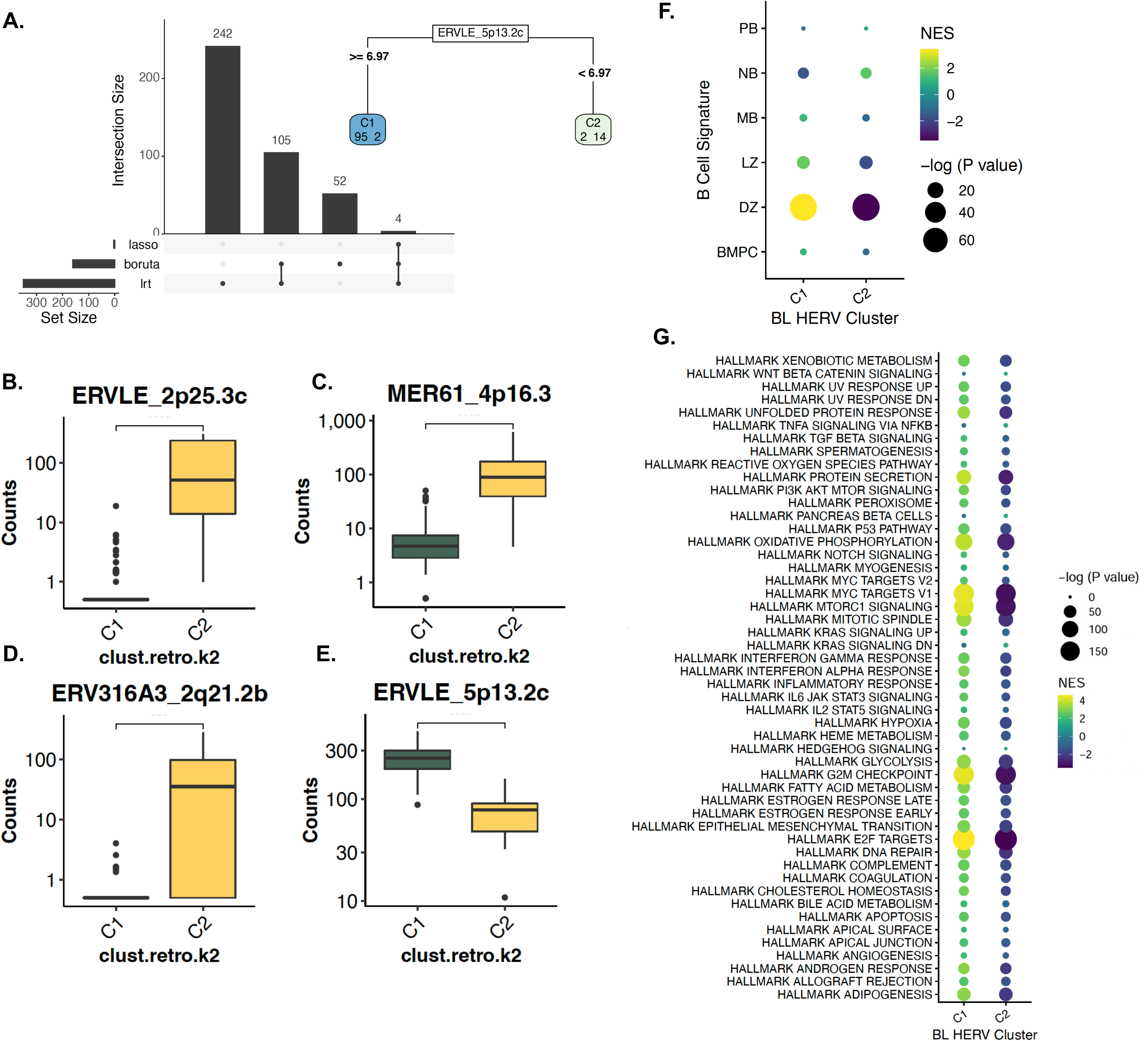
BL subtypes and EBV status have distinct biological and HERV signatures. **A.** Feature selection of differentially-expressed HERVs per cluster using DESeq2 LRT, Boruta, and Lasso find 4 HERVs sufficient to distinguish between BL-C1 and BL-C2, including **B.** ERVLE_2p25.3c, **C.** MER61_4p16.3**, D.** ERV316A3_2q21.2b, and **E.** ERVLE_5p13.2c. **F.** Gene and HERV-driven B-cell-of-origin classification of each HERV-driven BL cluster. Signature gene sets were created using a subset of the top 150 and top 25 upregulated genes and HERVs per cell-type from the Agirre 2019 B cell dataset. BL-C1 displays an enrichment of DZ gene-sets compared to BL-C2. **G.** Enrichment of hallmark pathways for the two HERV clusters, showcasing an overall upregulation in BL-C1 compared to BL-C2 for MYC targets, E2F targets, and epithelial mesenchymal transition.

### HERV expression is linked with survival outcomes in DLBCL

Finally, we hypothesized that the seven HERV-driven DLBCL subclasses with distinct predictive COO would display retrotranscriptomic differences that correlate with their prognostic outcome. We implemented an FGSEA analysis with the Hallmark pathways collected from MsigDB to calculate broad phenotypic alterations between our COO subtypes (Fig. 7A). HC1/ABC-PB displayed an overall downregulation of most Hallmark pathways, although HC2/ABC-MB, which was enriched for MB and DZ signatures, showed the highest enrichment for the “MYC targets V1”, “G2M checkpoint”, and “E2F targets” pathways. HC3/GCB-LZ displayed enrichment for “epithelial mesenchymal transition”, “mitotic spindle”, and a negative enrichment for the “DNA repair”, “interferon alpha and gamma response”, “MYC targets V1”, “MYC targets V2”, and “oxidative phosphorylation” pathways. HC4/GCB-like was enriched in “oxidative phosphorylation”, “MYC targets V1”, “epithelial mesenchymal transition”, and “adipogenesis” pathways, while HC6/GCB displayed a negative enrichment of “MYC targets V1” and “MYC targets V2” pathways. HC7/HERVH displayed an overall negative enrichment for most Hallmark pathways compared to the other clusters. The HC5/PB-Like showed a highly significant enrichment of the “interferon gamma and alpha response”, “inflammatory response”, “IL6 JAK STAT3 signaling”, “TNFA signaling via NFKB”, and “IL2 STAT5 signaling” pathways. Overall, samples from the HC5/PB-Like cluster had the highest enrichment for pathways indicating changes in local immunity (Supplementary Fig. 16), including “Cytotoxic T-lymphocyte-associated protein 4 (CTLA4)”, “TCR”, “IL17”, “IL10”, and “IL12”.

**Figure 7:**
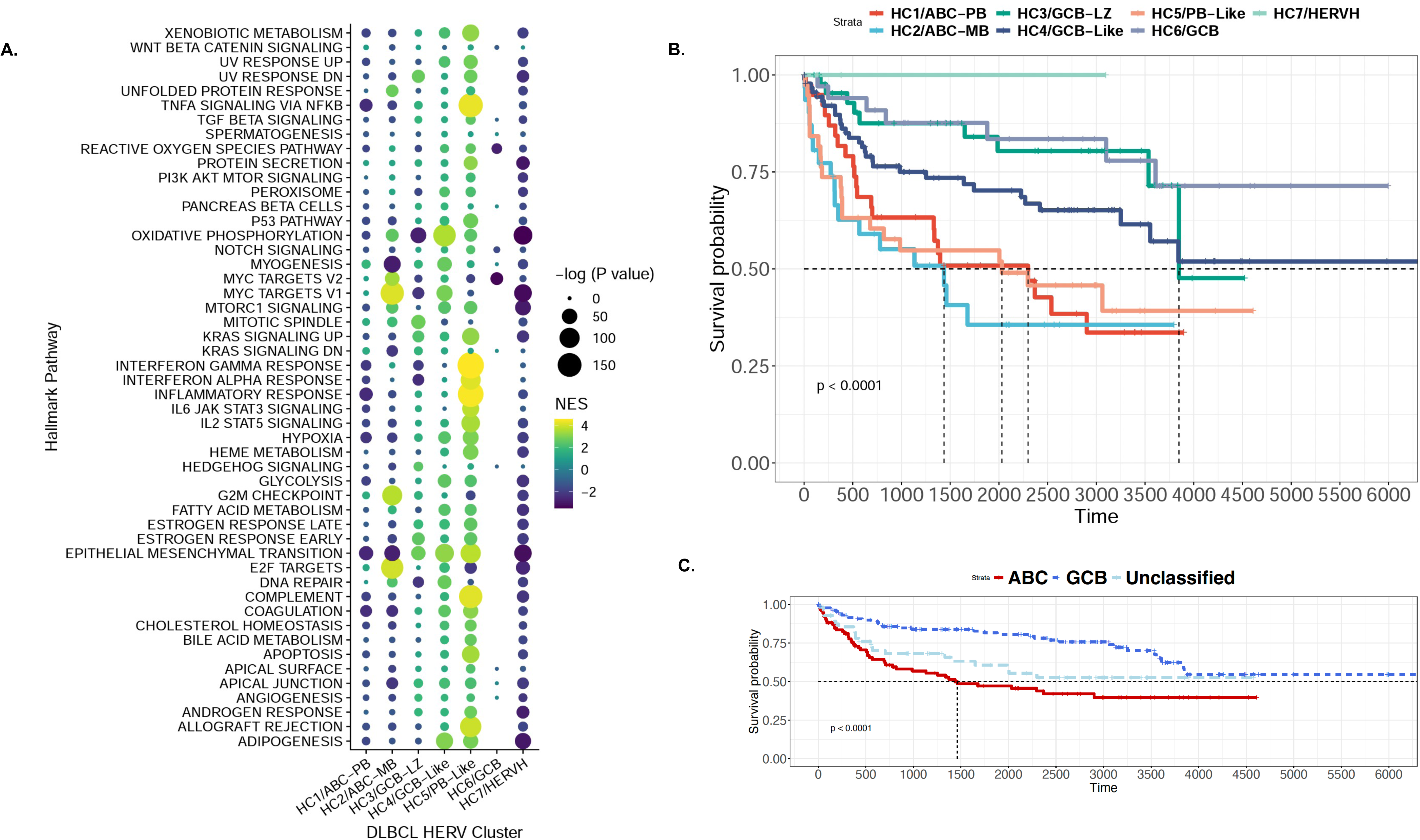
HERV-driven DLBCL subtypes have distinct biological properties and survival outcomes. **A.** Enrichment of hallmark pathways for the seven HERV clusters, showcasing distinct enrichment patterns for each cluster. HC1, which contains predominantly ABC-DLBCL and is enriched for NB, PB, and BMPC signatures, displays an overall downregulation of most hallmark pathways. HC2, which contains predominantly ABC-DLBCL and is enriched for MB and DZ signatures, shows the highest enrichment for MYC targets V1, G2M checkpoint, and E2F targets. HC3, which is a mixed cluster with LZ signatures, shows enrichment for epithelial mesenchymal transition, mitotic spindle, and a negative enrichment for DNA repair, interferon alpha and gamma response, MYC targets, and oxidative phosphorylation. HC4, which consists predominantly of GCB-DLBCL and displays LZ and DZ signatures, is enriched in oxidative phosphorylation, MYC targets V1, epithelial mesenchymal transition, and adipogenesis. HC5, which is another mixed cluster with BMPC signatures, shows a highly significant enrichment of interferon gamma and alpha response, inflammatory response, IL6 JAK STAT3 signaling, TNFA signaling via NFKB, and IL2 STAT5 signaling. HC6 shows a negative enrichment of MYC targets V2. HC7 displays an overall negative enrichment for most pathways compared to the other clusters. **B.** Survival plot of the seven DLBCL clusters, showcasing the worst prognosis for HC1/ABC-PB (n=39) and HC2/ABC-MB (n=31), followed by HC5/PB-Like (n=38), HC4/GCB-Like (n=92), HC3/GCB-LZ (n=45) and HC6/GCB (n=34), and HC7/HERVH (n=3). **C.** Survival plot of the original DLBCL cell-of-origin classifications, showing the worst prognosis for ABC-DLBCL, followed by Unclassified-DLBCL, and GCB-DLBCL.

We performed a Kaplan-Meier analysis to examine the relationship between HERV clusters and clinical outcomes, in comparison with previous COO classifications (Fig. 7B-C). Consistent with previous findings^62^, ABC-associated groups had the shortest long-term survival. Groups with the worst prognoses were HC1/ABC-PB (n=39) and HC2/ABC-MB (n=30), followed by HC5/PB-Like (n=37), HC4/GCB-Like (n=89), HC3/GCB-LZ (n=45), HC6/GCB (n=34), and HC7/HERVH (n=3) (Fig. 7B). Importantly, when implemented on the same cases denoted as ABC-DLBCL, GCB-DLBCL, or unclassified, the HERV-based classifications identified patient subsets that significantly correlated with prognostic outcomes. Patients in the HC5/PB-Like cluster (43% ABC, 45% Unclassified, 12% GCB) had a survival outcome much closer to the ABC-like clusters HC1 and HC2, despite having a large proportion of unclassified and GCB diagnoses. Similarly, prognostic values of previously unclassified DLBCLs had a significant range of favorable to unfavorable outcomes (Supplementary Fig. 17). Overall, novel DLBCL subclasses based on HERV signatures were able to be predictive of prognostic outcomes within the ABC-DLBCL, GCB-DLBCL, and Unclassified-DLBCL cases.

## Discussion

Prior to this study, there was limited data on HERV expression in both healthy and malignant proliferating B-cells, partly due to the challenges of TE quantification^36^. In this study, we developed the first comprehensive locus-specific atlas of TE expression in human GC B cells and in B-cell malignancies arising out of the GC. The GC reaction is a focal component of the adaptive immune response, where NBs travel to the follicles of secondary lymphoid organs to respond to T-cell dependent antigen challenges^63^. Through repeated cycling of proliferation and somatic hypermutation in the DZ and affinity selection in the LZ, B-cells terminally differentiate into either MBs or PBs^63^. Following development in the GC, PBs then migrate to the bone marrow to facilitate long-term humoral immunity by becoming BMPCs^64^. In malignant transformation events, this pathway of B-cell development is expropriated and gives rise to lymphomagenesis^65^. Disease-specific expression of HERVs have been previously noted as diagnostic markers^66, 67^ and further postulated as therapeutic targets for the treatment of B-cell lymphomas^27^. By characterizing the retrotranscriptome in the healthy GC and associated B-cell lymphomas, we identified HERVs specific to stages of the GC reaction and used them to further classify the COO in B-cell malignancies.

Our analyses of GC B-cells have enabled the construction of a reference of normal HERV expression during the various stages of B-cell maturation. As has been observed in other normal human tissue^8, 53, 54^, we found that HERV expression in B-cell subpopulations is highly specific to the cell types, including fully mature B-cells. The level of TE expression ranges throughout the GC reaction, with higher expression in NB cells, moderately high in the LZ, lower in the DZ, and higher again in MB, PB, and BMPC. Despite having relatively high TE transcription, PB and BMPC had the lowest percent of HERV fragments, and contradictorily, the highest number of uniquely upregulated HERV loci. Loci belonging to the HERVP71A family were highly expressed in the PB, potentially indicating the importance of this HERV family’s expression in PB cell fate. Importantly, despite HERV expression representing under 1% of the coding and non-coding transcriptome, our analysis demonstrates that HERV expression alone is able to independently distinguish GC cell types. We identified a signature based on 11 HERV markers to classify GC B-cells.

It is generally accepted that the COO for many of the non-Hodgkin B-cell lymphomas is a germinal center B cell, as indicated by the detection of somatically mutated immunoglobulin genes in their genomes. BLs are thought to be derived from the DZ, while FL and GCB-DLBCL resemble LZ cells, and ABC-DLBCLs are broadly derived from GC cells arrested during plasma cell differentiation^34, 65^. To further define these transformation events, we combined the HERV signature with gene expression data in these cell types to define their B-cell lineage. In doing this, we were able to confirm that HERV expression in these non-Hodgkin B-cell lymphomas corresponded with their previously identified GC COO. Like for the case of GC B-cells, HERV transcripts were again able to better distinguish lymphoma types than by analyzing gene expression alone, particularly between DLBCL and FL. In accordance with previous findings, BL samples most closely resembled the DZ, GCB-DLBCL resembled the LZ, ABC-DLBCL corresponded with MBs, PBs, and BMPCs, and FL corresponded with the LZ and NBs. We also found specific HERV markers of GC B-cell types upregulated in their associated B-cell lymphomas, including the DZ-associated element HARLEQUIN_19p12.b as a key marker of BL. Similarly, HARLEQUIN_1q32.1, which is a PB-associated HERV, is a key marker of ABC-DLBCL and has been previously implicated in prostate, breast, and colon cancers^52^.

We also used HERV signatures to expand the current COO classifications from three subsets into seven subsets, with each corresponding to single or mixed B-cell subtypes from the GC. Together, the two ABC-like clusters represented precursors to PBs and MBs. Recent findings have reported intermediate phases of the GC between the LZ and DZ compartmentalization^31^, in addition to MB precursors which are reflected in a fraction of DLBCLs^68^. These studies are consistent with our findings of “mixed” DLBCLs with competing gene and HERV signatures, particularly HC2/ABC-MB, which had an MB-like signature and encompassed a large number of MCD-DLBCL cases. The HC2/ABC-MB cases were most enriched in “MYC targets v1” and “MYC targets v2” pathways, both of which have been associated with tumor aggressiveness and proliferation^68^. The HC2/ABC-MB cases also had a significant upregulation of FABP7, a gene that is known to form TE chimeric transcripts and is upregulated in a subset of DLBCL cases^27^. HC1/ABC-PB and HC2/ABC-MB also showed increased expression of PRMD15, which is known to regulate multiple oncogenic pathways^69^.

One of the key features differentiating between our classification system for DLBCLs was HML2_7p22.1, a largely intact HERV provirus which contains an open reading frame (ORF) for a fusogenic retroviral envelope gene^53, 70, 71^. HML2_7p22.1 is one of two HERVs from the HML2 family that possess an intact envelope^53^. While HML2_7p22.1 is expressed in 15 different human tissue types^53^, it has also been implicated for its fusogenic activity in melanoma cell lines^72^ and may be immunosuppressive in nature^73^.

In the retrotranscriptome, BL had a threefold higher proportion of HERV transcription compared to DLBCL and FL, and significant upregulation in the number of DE HERVs. These data suggest aberrant overexpression of HERVs in BL, and further demonstrate the importance of their investigation in lymphomagenesis. The entirely EBV- BL-C2 cluster displayed a broad upregulation of HERVs in comparison to the largely EBV+ BL-C1 cluster, which was conversely associated with a stronger DZ signature. This suggests separate mechanisms of HERV-mediated malignancy in our two clusters of BL.

To test the clinical significance of our HERV-based clustering technique, we assessed the prognostic outcomes of the individually classified subgroups. Our HERV-based clustering identified additional sub-clusters within the GCB-like and unclassified DLBCL cases that demonstrate distinct survival outcomes. Briefly, the ABC-like clusters predictably had the least favorable prognostic outcomes. The GCB-like clusters displayed greater range than what would have been defined as a single class, with the HC3/GCB-Like cases having worse survival outcomes compared to HC4/GCB-LZ and HC6/GCB. Cases from the HC5/PB-Like cluster, which is likely to originate from intermediate phases of the GC reaction, had the least favorable outcomes, second only to the ABC-like clusters. This difference in clinical outcome may be attributed to the drastic changes observed in immune signatures within this cluster. The HC7/HERVH cluster lacked survival data to draw definitive conclusions, and therefore requires further investigation. However, this cluster demonstrated a clear downregulation of most Hallmark pathways expressed in the majority of our DLBCL clusters and is likely phenotypically distinct.

Taken together, our current analysis of healthy GC-B cells and B cell lymphomas suggests that malignant cells may retain both transcriptomic and retrotranscriptomic signatures from their COO. The observed increase in HERV transcripts in cancerous tissue, particularly BL, suggests a change in the epigenetic state of the B-cell derived COO in relation with infection status. This is relevant for other EBV and HIV-1 associated B cell lymphomas as well, where infection status may promote differential patterns of HERV expression. The identification of overexpressed HERV ORFs in cancer is of great interest for the pharmacological intervention of human malignancies due to the specificity of TE-derived tumor specific antigens^42^. Notably, these TE antigens are overexpressed under malignant conditions due to changes in the retrotranscriptome and have improved upon existing immunotherapies as novel targets^39, 40, 74–76, 78–82^. Overall, the predictive capabilities of the HERV-driven lymphoma clustering suggest a further need to understand the regulatory, transcriptional, and post-transcriptional activity of these endogenous retroelements in both healthy tissues and in malignant states. The characterization of HERV expression in the healthy GC and B cell lymphomas should therefore serve as a resource for the diagnostic and therapeutic potential of these elements in malignancies.

## Methods

### Data Availability

All samples were obtained from previously published studies^31, 32, 50, 76, 77^. Samples belonging to the B-AG (n=35) and B-HM (n=17) datasets were downloaded as FASTQ files using fasterq-dump from the SRA toolkit. RNA-seq data from the HIV-DLBCL samples (n=529) belonging to the TCGA and NCICCR research programs were obtained via the Genome Data Commons (dbGaP). Samples datasets were downloaded as FASTQ files using fasterq-dump from the SRA toolkit. RNA-seq data from the HIV-DLBCL samples (n=529) belonging to the TCGA and NCICCR research programs were obtained via the dbGaP accession “phs001444.v2.p1”. The BL samples (n=113) were obtained as part of CGCI’s Burkitt Lymphoma Genome Sequencing Project (BLGSP), and accessed via dbGaP accession “phs000235.v16.p4”. The FL samples (n=12) were obtained as part of CGCI’s Non-Hodgkin Lymphoma - Follicular Lymphoma (NHL - FL) initiative and accessed through SRA toolkit via the dbGaP accession “phs000235.v7.p2”. Clinical, demographic, and survival metadata was obtained via the TCGABiolinks R package (v2.18.0). LymphGen^35^, EcoTyper^49^, *Chapuy et al.*^78^, and *Holmes et al.*^31^ DLBCL classification calls were obtained from the respective publications.

### Data processing pipelines and code availability

Custom and reproducible Snakemake (v7.14.0) pipelines were created for the DLBCL, (https://github.com/nixonlab/DLBCL_HERV_atlas_GDC), BL (https://github.com/nixonlab/burkitt_lymphoma_TE_atlas), FL (https://github.com/nixonlab/follicular_lymphoma_TE_atlas), and healthy B-cell datasets (https://github.com/nixonlab/HERV_GCB_Bulk), separated by the source of data access. Input samples were supplied through the config.yaml file for each pipeline, which also contained consistent parameters for data processing. The same package versions were used for gene and TE quantification in each Snakemake pipeline^79^. All downstream analysis was conducted in R (v4.0.2), and can be accessed on GitHub (https://github.com/singhbhavya/hematological_malignancies_te_analysis).

### Transcriptomic profiling and locus-specific HERV prediction

For DLBCL and BL, downloaded BAM files were converted to FASTQ using picard-slim (v2.25). FASTQ files for all samples were then aligned to Hg38 using STAR (v2.7.9a), with parameters “--outSAMattributes NH HI NM MD AS XS --outSAMtype BAM Unsorted --quantMode GeneCounts --outSAMstrandField intronMotif --outFilterMultimapNmax 200 --winAnchorMultimapNmax 200 -- outSAMunmapped Within KeepPairs”. We used Telescope (v1.0.3) for retrotranscriptomic profiling, which allows for the locus-specific identification of TEs using expectation maximization algorithm. The Telescope assign module was used with the parameters “--theta_prior 200000 -- max_iter 200”, along with a custom transposable element annotation (retro.hg38.v1), accessible at https://github.com/mlbendall/telescope_annotation_db. Meta annotations for TEs with the nearest genes, gene overlaps, and the TE status of intronic, exonic, or intergenic, were obtained from https://github.com/liniguez/Telescope_MetaAnnotations.

### Unsupervised clustering

Gene and TE counts were first filtered, such that only features with more than 5 observations within a minimum sample threshold (5 samples for the 529 DLBCLs, 5 samples for 113 BLs, 2 samples for 12 FLs, 2 samples for 17 B-HM, and 4 samples for 35 B-AG) were retained. Normalized counts were calculated using the estimated size factors within DESeq2 (v1.30.1), and subsequently transformed using variance-stabilizing transformation^61^. PCA was carried out on the transformed counts, and then visualized using PCATools (v2.2.0). Clustering on DLBCL and BL samples was performed using ConsensusClusterPlus (v1.54.0), with 1000 repetitions. Clusters were calculated for *k*=2 through *k*=9, and assessed through the calculated consensus matrices, silhouette statistics, molecular and clinical indicators, and agreement with previously-described classifications. Final clusters of *k*=7 for DLBCL and *k*=2 for BL were chosen based on the aforementioned statistical and clinical indicators. Fisher’s exact test was used to test each cluster against categorical variables and previous classifications. Alluvial plots comparing HERV clusters to previous DLBCL and Bl classifications were created using ggalluvial (v0.12.3).

### Differential expression analysis

DE testing was performed between and within lymphoma subtypes, and separately within B-cell subtypes for the B-AG and B-HM datasets. A negative binomial model was used for DE testing, with a significance cutoff of p=0.001, and a log2fold change cutoff of >1.5. B-cell subtypes were compared individually within the B-AG and B-Hm datasets, with a design of ∼cell_type + 0. To compare between lymphoma types, two DE models were created, with the broad lymphoma type (∼ cancer_type + 0, where cancer_type refers to DLBCL, BL, or FL), and a narrower lymphoma subtype (∼ subtype + 0, where the subtypes included ABC-DLBCL, GCB-DLBCL, Unclassified, EBV+/- Sporadic and Endemic BL, and FL). Differential expression testing was also performed within DLBCL (∼COO + 0), BL (∼ebv_status + 0), and the unsupervised HERV clusters for DLBCL (∼ clust.retro.k7 + 0) and BL (clust.retro.k2 + 0) respectively. Results were extracted as DESeqResults objects, with a numbered contrast of each group compared against all others. HERVs that were uniquely upregulated and downregulated per group were visualized with UpsetR (1.4.0) and ComplexUpset (1.3.3). The top n differentially expressed genes and HERVs were visualized with pheatmap (1.0.12). The significance and effect size of DE genes and HERVs were calculated and visualized with EnhancedVolcano (1.8.0).

### HERV-based feature selection and model

Supervised learning and HERV-based feature selection was implemented as previously described^80^. Briefly, pre-filtered HERV matrices were used for DESeq2’s likelihood ratio test (LRT), which was used to create a model of the HERV clusters for DLBCL (∼clust.retro.k7 + 1). BL (∼clust.retro.k2 + 1), and healthy B-cells from the B-AG dataset (∼cell_type +1), with a significance cutoff of FDR < 0.001. Variance transformed counts from DESeq2 were extracted for feature selection with the Boruta random forest algorithm and the randomized LASSO regression. LASSO regression with stability selection was used to find the minimum optimal numbers of features defining each group, using the glmnet (v4.1-6) and c060 (v0.2-9) packages. LASSO was implemented with multinomial logistic regression with a grouped penalty, ensuring that each selected feature had multinomial coefficients of either all non-0 or all 0. Stability selection was performed with 200 subsamples, and a proportion threshold of 0.6. For a less stringent feature selection of all relevant features, we used the Boruta (v8.0.0) algorithm and the randomForest package (v4.6-12) for random classification, with ntree = 1000 and maxRuns = 1000. Final features were selected using an intersection of the three methods and visualized with UpsetR. The LASSO signature was used to create a final classification tree, with recursive partitioning implemented in rpart (v4.1.19) and rpart.plot (v3.1.1).

### HERV- and gene-set enrichment analyses

Preranked gene-set enrichment analysis (GSEA) was performed using the fgsea package (v1.16.0), which uses an adaptive multilevel split Monte Carlo method. Fold change statistics and p-values from DESEq2 differential testing were used to estimate gene and HERV ranks. Overall biological signatures in BL and DLBCL HERV clusters were calculated using the Hallmark and Kyoto Encyclopedia of Genes and Genomes^81^gene sets from MSigDB^61^. We created custom B-cell signature gene sets, using the top 150 genes and top 25 HERVs upregulated in each B-cell subtype in the B-AG dataset, and performed a combined HAGSEA to determine potential COO of our DLBCL and BL HERV clusters. Effect size and p-values were visualized for the GSEA and HAGSEA using corrplot (v0.92) in R.

### Survival analysis

Survival analysis was conducted using the survival R package (v3.1-12), using the log-rank test for group-level comparisons (rho=0). Kaplan-Meier survival plots were drawn using survminer (v0.4.9) and ggplot2 (v3.3.6).

### Statistical analyses

All analyses were performed in Bash, R (v4.0.2), and the BioConductor package manager (v1.30.19). Significance values for all DE analyses were calculated with the Wald test, with the Benjamini and Hochberg method for multiple testing correction. Comparisons between mean HERV and gene expression were conducted with the t-test, on normalized counts from DESeq2. Feature selection was performed using the multiple likelihood ratio test in DESeq2, the Boruta random forest algorithm, and the randomized LASSO regression.

## Acknowledgments

The work was supported in part by US National Institutes of Health (NIH) grant CA260691 (DFN). MLB is supported in part by the Department of Medicine Fund for the Future program at Weill Cornell Medicine sponsored by the Elsa Miller Foundation. JLM was supported in part by a Medical Scientist Training Program grant to the Weill Cornell–Rockefeller–Sloan Kettering Tri-Institutional MD-PhD Program (T32GM007739), and by a grant from the Melanoma Research Foundation CK0041482163, generously supported by the Silverstein family.

The results shown here are in whole or part based upon data generated by the TCGA Research Network: https://www.cancer.gov/tcga, including data generated by the Cancer Genome Characterization Initiative (phs000235), developed by the National Cancer Institute. Information about CGCI projects can be found at https://ocg.cancer.gov/programs/cgci. The Genomic Variation in Diffuse Large B-cell Lymphomas study was supported by the Intramural Research Program of the National Cancer Institute, National Institutes of Health, Department of Health and Human Services. The datasets have been accessed through the NIH database for Genotypes and Phenotypes (dbGaP).

We would like to acknowledge helpful discussions with members of the Cesarman, Feschotte, and Leal labs, in addition to the overall HERV Lymphoma team.

## Author Contributions

Study design and conception: B.S. M.L.B., D.F.N. Performed analyses: B.S. Wrote the paper: B.S. Provided support with data analysis and interpretation: T.F., N.D., J.L.M. Created conceptual figures: S.M. Contributed knowledge, revised the manuscript: all authors.

## Declaration of Interests

Peter Martin: ADCT: Consultancy. All other authors declare no competing interests.

**Supplementary Figure 1: Unique and differentially expressed HERV loci in the B-HM dataset. A.** Upset plot of the number of unique and shared HERVs upregulated in each B cell type (p < 0.001, log2fold change > 1.5**). B.** Upset plot of the number of unique and shared HERVs downregulated in each B cell type (p < 0.001, log2fold change > 1.5). **C.** Volcano plot of differentially expressed HERVs in all cell types versus DZ, **D.** all versus LZ, **E.** all versus MB, and **F.** all versus NB.

**Supplementary Figure 2: Unique and differentially expressed HERV loci in the B-AG dataset. A.** Upset plot of the number of unique and shared HERVs upregulated in each B cell type (p < 0.001, log2fold change > 1.5**). B.** Upset plot of the number of unique and shared HERVs downregulated in each B cell type (p < 0.001, log2fold change > 1.5). **C.** Volcano plot of differentially expressed HERVs in all cell types versus DZ, **D.** all versus LZ, **E.** all versus MB, and **F.** all versus NB.

**Supplementary Figure 3: Plasmablasts and bone marrow plasma cells express distinct HERV profiles compared to GC B cells in the B-AG dataset. A.** Volcano plot of differentially expressed HERVs in all cell types versus BMPC**, B.** all versus PB**. C.** Heatmap of the top 75 upregulated genes and HERVs in PB (p < 0.001, log2fold change > 1.5), and **D.** BMPC.

**Supplementary Figure 4: Key features differentiating B-AG B cell subsets based on feature selection with DESeq2 LRT, Boruta, and Lasso. A.** UpsetR plot displaying the number of features selected by DESeq2 lowest likelihood ratio (LTR), the random forest classification with the Boruta algorithm, and the randomized least absolute shrinkage and selection operator (LASSO) regression, with 11 features being selected by all three methods. **B.** Rpart decision tree, displaying that HERVP71A_8q24.13 differentiates plasma cells (PB and BMPC) from the rest of the B cells. HERVL_2p12a differentiates DZ from the remaining cell types, while HUERSP2_6p22.3 differentiates LZ from MB and NB**. C.** Normalized counts plotted for the 11 HERV features differentiating the B cell subtypes: ERVLB4_14q23.3, HERVL_2p12a, HERVP71A_8q24.13, MER61_19p12c, HARLEQUIN_19p12b, HERVFRD_2p12a, PABLB_7q11.21, HERVL_1q23.3a, HERVP71A_15q24.2, HUERSP2_6p22.3, ERVLE_6p25.1b.

**Supplementary Figure 5: Total % of reads assigned to TEs and HERVs by lymphoma type and sub-type. A.** Mean of the percentage (%) of reads assigned to TEs in BL, DLBCL, and FL, and **B.** their respective subtypes. **C.** Mean of the percentage (%) of reads assigned to HERVs in BL, DLBCL, and FL, and **D**. their respective subtypes.

**Supplementary Figure 6: HERV upregulation and downregulation in lymphoma subtypes.**

**A.** Upset plot of the number of unique and shared HERVs upregulated in each cancer sub-type, including ABC, GCB, and unclassified DLBCL, sporadic and endemic BL by EBV status, and follicular lymphoma. **B.** Upset plot of the number of unique and shared HERVs downregulated in each cancer sub-type. **C.** Relative abundance of HERV families per lymphoma sub-type. GCB-DLBCL contains the highest number of upregulated HERV loci.

**Supplementary Figure 7: Upregulation of DZ-associated HERVs in BL compared to DLBCL and FL.** Four DZ-associated HERVs are significantly upregulated in BL compared to DLBCL and FL, as determined by a t-test to compare the means (p < 0.05). **A.** MER61_3q13.11, **B.** HML5_1q22**, C.** HERV3_14q32.33, **D.** HARLEQUIN_19p12b.

**Supplementary Figure 8: Upregulation of PB-associated HARLEQUIN_1q32.1 in ABC-DLBCL compared to other lymphoma subtypes.** HARLEQUIN_1q32.1, which is **A**. associated with BMPC and PB, is significantly upregulated in **B.** ABC-DLBCL compared to GCB-DLBCL and unclassified-DLBCL and BL (t-test, p < 0.005).

**Supplementary Figure 9: Unsupervised HERV-based classification of DLBCL samples compared to previous classifications. A.** Alluvial plot of 529 DLBCL samples, and their respective class calls for the COO classifications, DBL Hit status, scCOO group, Chapuy group, EcoTyper class, and Lymphgen class, compared to the HERV-based clusters. Transcriptomic and retrotranscriptome signatures do not clearly segregate the samples based on previous classification, as observed in **B.** Gene-based PCA plot of 529 DLBCL samples, colored by COO classification, **C.** HERV-based PCA plot of 529 DLBCL samples, colored by COO classification, **D.** Gene-based PCA plot of 529 DLBCL samples, colored by EcoTyper classes, **E.** HERV-based PCA plot of 529 DLBCL samples, colored by EcoTyper classes, **F.** Gene-based PCA plot of 529 DLBCL samples, colored by LymphGen classifications, and **G.** HERV-based PCA plot of 529 DLBCL samples, colored by LymphGen classifications.

**Supplementary Figure 10: Key features differentiating B-AG B cell subsets based on feature selection with DESeq2 LRT, Boruta, and Lasso**. **A.** UpsetR plot displaying the number of features selected by DESeq2 lowest likelihood ratio (LTR), the random forest classification with the Boruta algorithm, and the randomized least absolute shrinkage and selection operator (LASSO) regression, with 3 features being selected by all three methods, and 4 by both LASSO and Boruta. **B** Normalized counts plotted for the 4 HERV features differentiating the B cell subtypes: **B.** HML2_7p22.1, **C.** HERVH_16p13.2e, **D.** HERVW_2q23.3, and **E.** HERVH_7q11.23a. **F.** Rpart decision tree, displaying that HERVH_16p13.2e differentiates HC7 from the remaining clusters. HERVW_2q23.3 differentiates HC1 and HC2 from the remaining clusters, and then further differentiates HC2 from HC1, where its expression is the highest. HML2_7p22.1 separates HC4 and HC6 from HC3, HC4, and HC7, and then further differentiates within the clusters. HERVH_7q11.23a differentiates HC2 from HC3, HC4 from HC6, and HC7 from HC3 and HC5.

**Supplementary Figure 11: ABC-like DLBCL clusters with unique HERV signatures**. HC1 and HC2 clusters contained the highest number of ABC-DLBCL samples. Top 75 differentially expressed genes and HERVs (p < 0.001, log2fold change > 1.5) in **A.** HC1, and **B.** HC2.

**Supplementary Figure 12: GCB-like DLBCL clusters with unique HERV signatures.** HC3 and HC4 clusters contained the highest number of GCB-DLBCL samples. Top 75 differentially expressed genes and HERVs (p < 0.001, log2fold change > 1.5) in **A.** HC3, and **B.** HC4.

**Supplementary Figure 13: PB-like and Post-GCB DLBCL clusters with unique HERV signatures.** Top 75 differentially expressed genes and HERVs (p < 0.001, log2fold change > 1.5) in **A.** The HC5 cluster, which was most associated with the PB cell-of-origin, and **B.** HC7 cluster, which was enriched in PB, BMPC, and MB.

**Supplementary Figure 14: HERV upregulation and downregulation in BL HERV clusters and clinical subtypes. A.** Volcano plot of differentially expressed HERVs in BL-C1 vs BL-C2 (p < 0.001, log2fold change > 1.5), **B.** EBV- versus EBV+. **C.** Relative abundance of loci assigned to HERV families the HERV-driven BL-C1 and BL-C2 clusters, and **D.** Comparing between all EBV negative, EBV positive, Endemic, Endemic EBV negative, Endemic EBV positive, Sporadic, Sporadic EBV negative.

**Supplementary Figure 15: Expression of selected BL features in other lymphoma subtypes.** Feature selection of differentially expressed HERVs in the two BL clusters found 4 HERVs sufficient to distinguish between BL-C1 and BL-C2. The same HERVs are also expressed in DLBCL and FL, but at different levels**. A.** ERVLE_2p25.3c is expressed most in Sporadic BL EBV negative**, B.** MER61_4p16.3 is expressed across lymphoma types, **C.** ERV316A3_2q21.2b has the highest expression in sporadic BL EBV negative, and **D.** ERVLE_5p13.2c is expressed in all lymphoma types, but with highest expression in BL.

**Supplementary Figure 16: Top enriched MSigDB gene sets and pathways in DLBCL HERV clusters. A.** Enrichment of Gene Ontology Biological Processes pathways for the seven HERV clusters, showcasing distinct enrichment patterns for each cluster. The most enriched pathways for HC1 were chromosome organization, chromatin remodeling, positive regulation of RNA metabolic process, ncRNA processes, mRNA metabolic process, and cellular response to DNA damage stimulus. The pathways most enriched in HC2 were rRNA processing, RNA processing, ribosome biogenesis, ribonucleoprotein complex biogenesis, ncRNA processing, ncRNA metabolic process, along with DNA metabolic process and chromosome organization. The pathways most enriched in HC3 were cell motility, cell adhesion, locomotion, epithelium development, response to endogenous stimulus. The pathways most enriched in HC4 were small molecule metabolic process, peptide and organonitrogen compound biosynthetic process, generation of precursor metabolites and energy, cytoplasmic translation, and amide metabolic processes. HC5 had an overall enrichment of immune response signatures. HC6 and HC7 did not have any positive enrichment. **B.** Enrichment of BioCarta pathways for the seven HERV clusters. Similar to the GO BP pathways, HC5 had the most striking enrichment of immune and inflammatory pathways.

**Supplementary Figure 17: HERV-driven DLBCL subtypes have distinct biological properties and survival outcomes for unclassified DLBCL samples.** Survival plot of five DLBCL clusters containing unclassified cases. HC3 and HC6 contained only one unclassified case each, and were thus omitted. HERV classes with the worst prognosis are HC2 and HC5, followed by HC1, HC4, and HC7.

