## Supplemental Information for "Locus specific human endogenous retroviruses reveal new lymphoma subtypes"

Supp Fig. 1

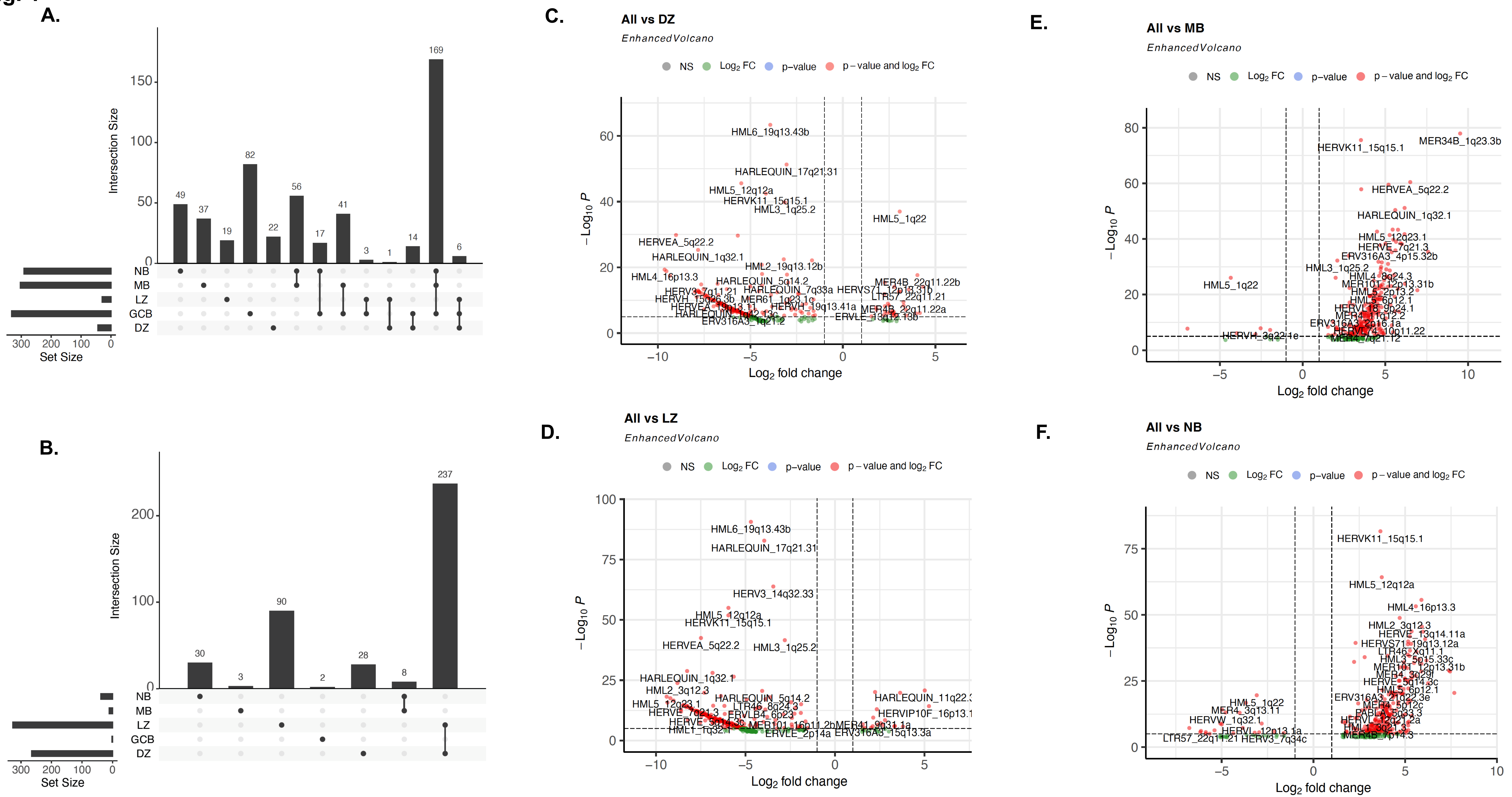

A.
Intersection Size
Set Size
NB
MB
LZ
GCB
DZ
B.
C.
All vs DZ
EnhancedVolcano
NS
Log2 FC
p–value
p – value and log2 FC
HML6_19q13.43b
HARLEQUIN_17q21.31
HML5_12q12a
HERVK11_15q15.1
HML3_1q25.2
HML5_1q22
HERVEA_5q22.2
HARLEQUIN_1q32.1
HML2_19q13.12b
HML4_16p13.3
HARLEQUIN_5q14.2
MER4B_22q11.22b
Log2 fold change
D.
All vs LZ
HERV3_14q32.33
HML2_3q12.3
HARLEQUIN_11q22.3
HERVIP10F_16p13.1
E.
All vs MB
MER34B_1q23.3b
HML5_12q23.1
HERVE_7q21.3
ERV316A3_4p15.32b
HERVH_3q22.1e
F.
All vs NB
HML2_3q12.3
HERVE_13q14.11a
HERVS71_19q13.12a
LTR46_Xq11.1
MER4_3q13.11
HERVW_1q32.1
LTR57_22q11.21
HERV3_7q34c

Supp Fig. 2

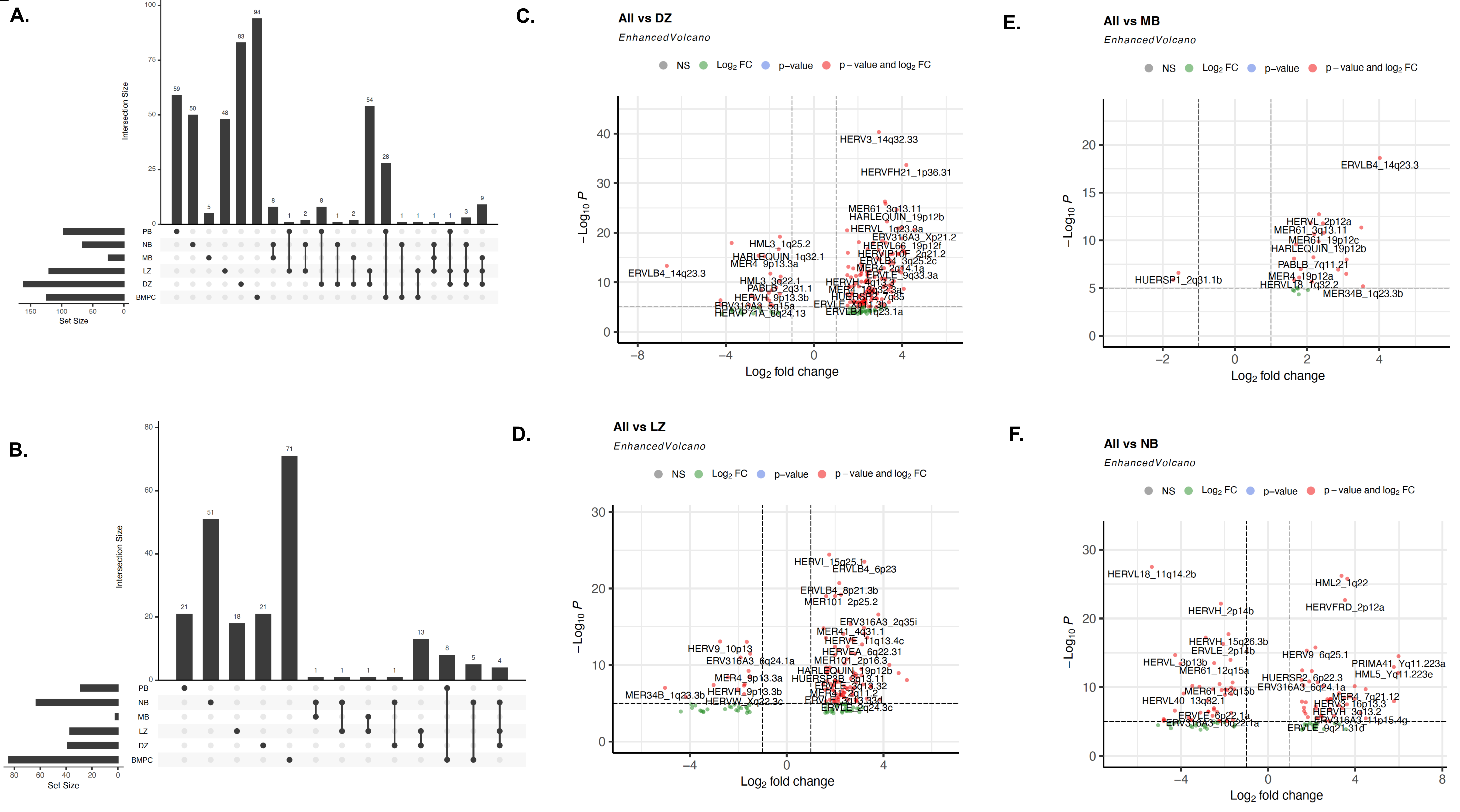
A.
B.
C.
D.
E.
F.
Intersection Size
Set Size
PB
NB
MB
LZ
DZ
BMPC
All vs DZ
EnhancedVolcano
All vs MB
All vs LZ
All vs NB
NS
Log2 FC
p–value
p – value and log2 FC
Log2 fold change
–Log10 P
ERVLB4_14q23.3
HERV3_14q32.33
HERVFH21_1p36.31
HERVL18_11q14.2b
HML2_1q22
HERVFRD_2p12a
HERVH_2p14b
HERVI_15q25.1
ERVLB4_6p23
MER34B_1q23.3b
HUERSP1_2q31.1b

**Supp Fig. 3**

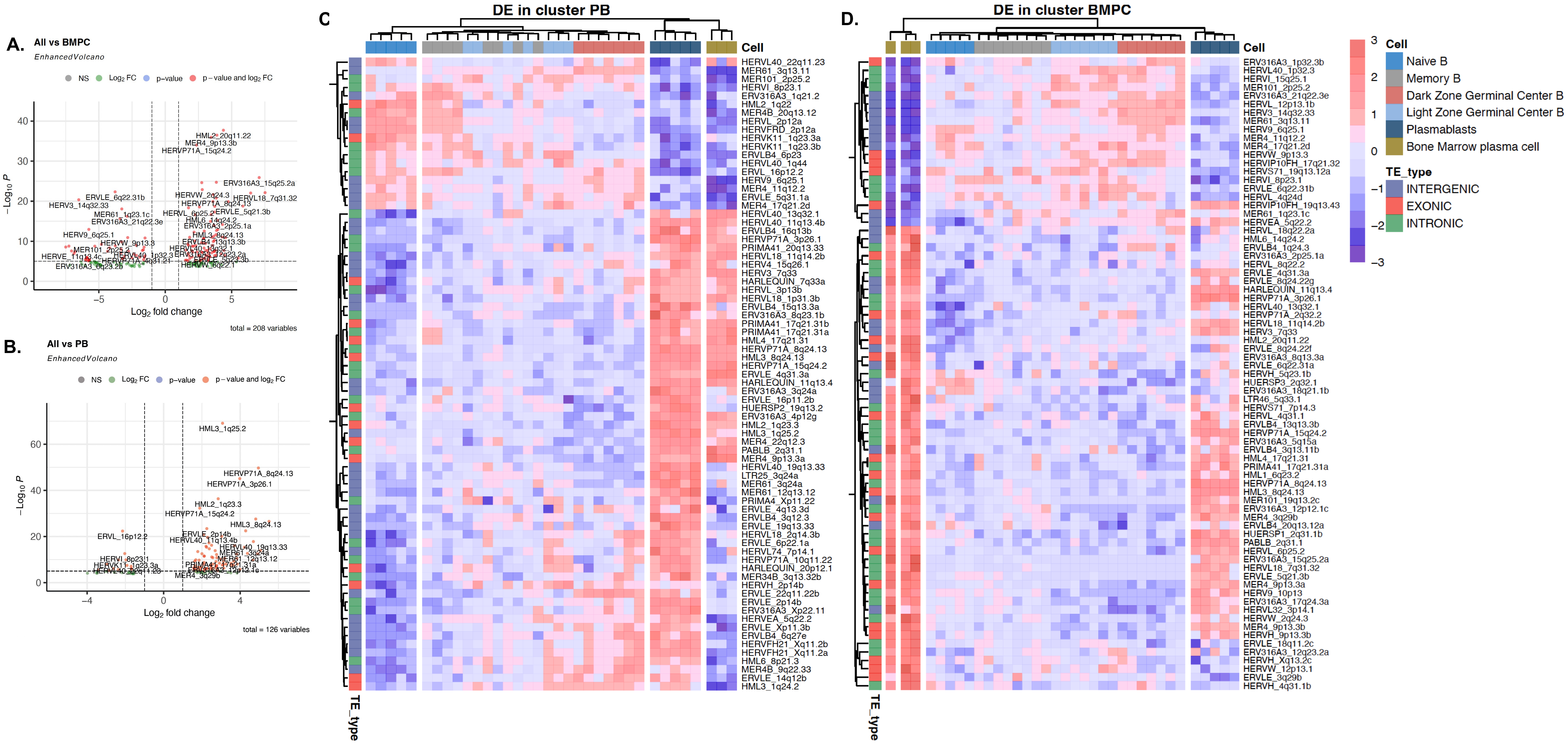
A.
All vs BMPC
EnhancedVolcano
NS
Log2 FC
p–value
p – value and log2 FC
−Log10 P
Log2 fold change
total = 208 variables
HML2_20q11.22
MER4_9p13.3b
HERVP71A_15q24.2
ERV316A3_15q25.2a
HERVL18_7q31.32
ERVLE_6q22.31b
HERV3_14q32.33
MER61_1q23.1c
ERV316A3_21q22.3e
HERV9_6q25.1
MER101_2p25.2
HERVE_11q13.4c
ERV316A3_6q23.2b
HERVW_2q24.3
HERVP71A_8q24.13
HERVL_6p25.2
ERVLE_5q21.3b
HML6_14q24.2
ERV316A3_2p25.1a
HML3_8q24.13
ERVLB4_13q13.3b
HERVL40_13q32.1
HERVW_9p13.3
HERVL40_1p32.3
HERVP71A_4q31.21
HERVW_6q22.1
B.
All vs PB
EnhancedVolcano
NS
Log2 FC
p–value
p – value and log2 FC
−Log10 P
Log2 fold change
total = 126 variables
HML3_1q25.2
HERVP71A_8q24.13
HERVP71A_3p26.1
HML2_1q23.3
HERVP71A_15q24.2
HML3_8q24.13
ERVLE_2p14b
HERVL40_11q13.4b
HERVL40_19q13.33
ERVL_16p12.2
HERVI_8p23.1
HERVK11_1q23.3a
HERVL40_22q11.23
MER61_3q24a
MER61_12q13.12
PRIMA41_17q21.31a
MER4_3q29b
C.
DE in cluster PB
Cell
TE_type
HERVL40_22q11.23
MER61_3q13.11
MER101_2p25.2
HERVI_8p23.1
ERV316A3_1q21.2
HML2_1q22
MER4B_20q13.12
HERVL_2p12a
HERVFRD_2p12a
HERVK11_1q23.3a
HERVK11_1q23.3b
ERVLB4_6p23
HERVL40_1q44
ERVL_16p12.2
HERV9_6q25.1
MER4_11q12.2
ERVLE_5q31.1a
MER4_17q21.2d
HERVL40_13q32.1
HERVL40_11q13.4b
ERVLB4_16q13b
HERVP71A_3p26.1
PRIMA41_20q13.33
HERVL18_11q14.2b
HERV4_15q26.1
HERV3_7q33
HARLEQUIN_7q33a
HERVL_3p13b
HERVL18_1p31.3b
ERVLB4_15q13.3a
ERV316A3_8q23.1b
PRIMA41_17q21.31b
PRIMA41_17q21.31a
HML4_17q21.31
HERVP71A_8q24.13
HML3_8q24.13
HERVP71A_15q24.2
ERVLE_4q31.3a
HARLEQUIN_11q13.4
ERV316A3_3q24a
ERVLE_16p11.2b
HUERSP2_19q13.2
ERV316A3_4p12g
HML2_1q23.3
HML3_1q25.2
MER4_22q12.3
PABLB_2q31.1
MER4_9p13.3a
HERVL40_19q13.33
LTR25_3q24a
MER61_3q24a
MER61_12q13.12
PRIMA4_Xp11.22
ERVLE_4q13.3d
ERVLB4_3q12.3
ERVLE_19q13.33
HERVL18_2q14.3b
ERVLE_6p22.1a
HERVL74_7p14.1
HERVP71A_10q11.22
HARLEQUIN_20p12.1
MER34B_3q13.32b
HERVH_2p14b
ERVLE_22q11.22b
ERVLE_2p14b
ERV316A3_Xp22.11
HERVEA_5q22.2
ERVLE_Xp11.3b
ERVLB4_6q27e
HERVFH21_Xq11.2b
HERVFH21_Xq11.2a
HML6_8p21.3
MER4B_9q22.33
ERVLE_14q12b
HML3_1q24.2
D.
DE in cluster BMPC
Cell
TE_type
ERV316A3_1p32.3b
HERVL40_1p32.3
HERVI_15q25.1
MER101_2p25.2
ERV316A3_21q22.3e
HERVL_12p13.1b
HERV3_14q32.33
MER61_3q13.11
HERV9_6q25.1
MER4_11q12.2
MER4_17q21.2d
HERVW_9p13.3
HERVIP10FH_17q21.32
HERVS71_19q13.12a
HERVI_8p23.1
ERVLE_6q22.31b
HERVL_4q24d
HERVIP10FH_19q13.43
MER61_1q23.1c
HERVEA_5q22.2
HERVL_18q22.2a
HML6_14q24.2
ERVLB4_1q24.3
ERV316A3_2p25.1a
HERVL_8q22.2
ERVLE_4q31.3a
ERVLE_8q24.22g
HARLEQUIN_11q13.4
HERVP71A_3p26.1
HERVL40_13q32.1
HERVP71A_2q32.2
HERVL18_11q14.2b
HERV3_7q33
HML2_20q11.22
ERVLE_8q24.22f
ERV316A3_8q13.3a
ERVLE_6q22.31a
HERVH_5q23.1b
HUERSP3_2q32.1
ERV316A3_18q21.1b
LTR46_5q33.1
HERVS71_7p14.3
HERVL_4q31.1
ERVLB4_13q13.3b
HERVP71A_15q24.2
ERV316A3_5q15a
ERVLB4_3q13.11b
HML4_17q21.31
PRIMA41_17q21.31a
HML1_6q23.2
HERVP71A_8q24.13
HML3_8q24.13
MER101_19q13.2c
ERV316A3_12p12.1c
MER4_3q29b
ERVLB4_20q13.12a
HUERSP1_2q31.1b
PABLB_2q31.1
HERVL_6p25.2
ERV316A3_15q25.2a
HERVL18_7q31.32
ERVLE_5q21.3b
MER4_9p13.3a
HERV9_10p13
ERV316A3_17q24.3a
HERVL32_3p14.1
HERVW_2q24.3
MER4_9p13.3b
HERVH_9p13.3b
ERVLE_18q11.2c
ERV316A3_12q23.2a
HERVH_Xq13.2c
HERVW_12p13.1
ERVLE_3q29b
HERVH_4q31.1b
Cell
Naive B
Memory B
Dark Zone Germinal Center B
Light Zone Germinal Center B
Plasmablasts
Bone Marrow plasma cell
TE_type
INTERGENIC
EXONIC
INTRONIC

**Supp Fig. 4**

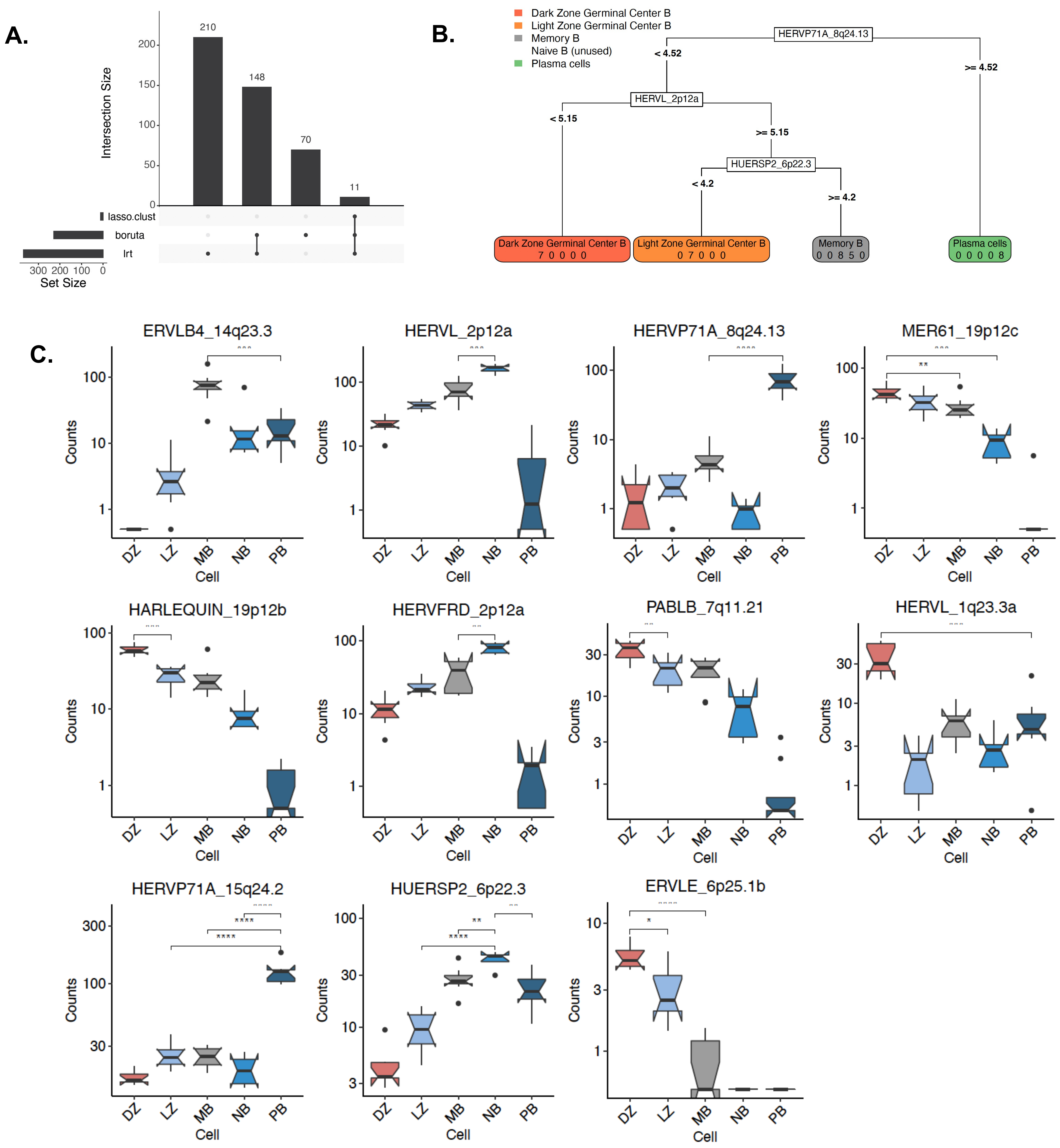

A.
Intersection Size
210
148
70
11
lasso.clust
boruta
lrt
Set Size
B.
Dark Zone Germinal Center B
Light Zone Germinal Center B
Memory B
Naive B (unused)
Plasma cells
HERVP71A_8q24.13
< 4.52
>= 4.52
HERVL_2p12a
< 5.15
>= 5.15
HUERSP2_6p22.3
< 4.2
>= 4.2
Dark Zone Germinal Center B 7 0 0 0 0
Light Zone Germinal Center B 0 7 0 0 0
Memory B 0 0 8 5 0
Plasma cells 0 0 0 0 8
C.
ERVLB4_14q23.3
HERVL_2p12a
HERVP71A_8q24.13
MER61_19p12c
HARLEQUIN_19p12b
HERVFRD_2p12a
PABLB_7q11.21
HERVL_1q23.3a
HERVP71A_15q24.2
HUERSP2_6p22.3
ERVLE_6p25.1b
Counts
Cell
DZ
LZ
MB
NB
PB

**Supp Fig. 5**

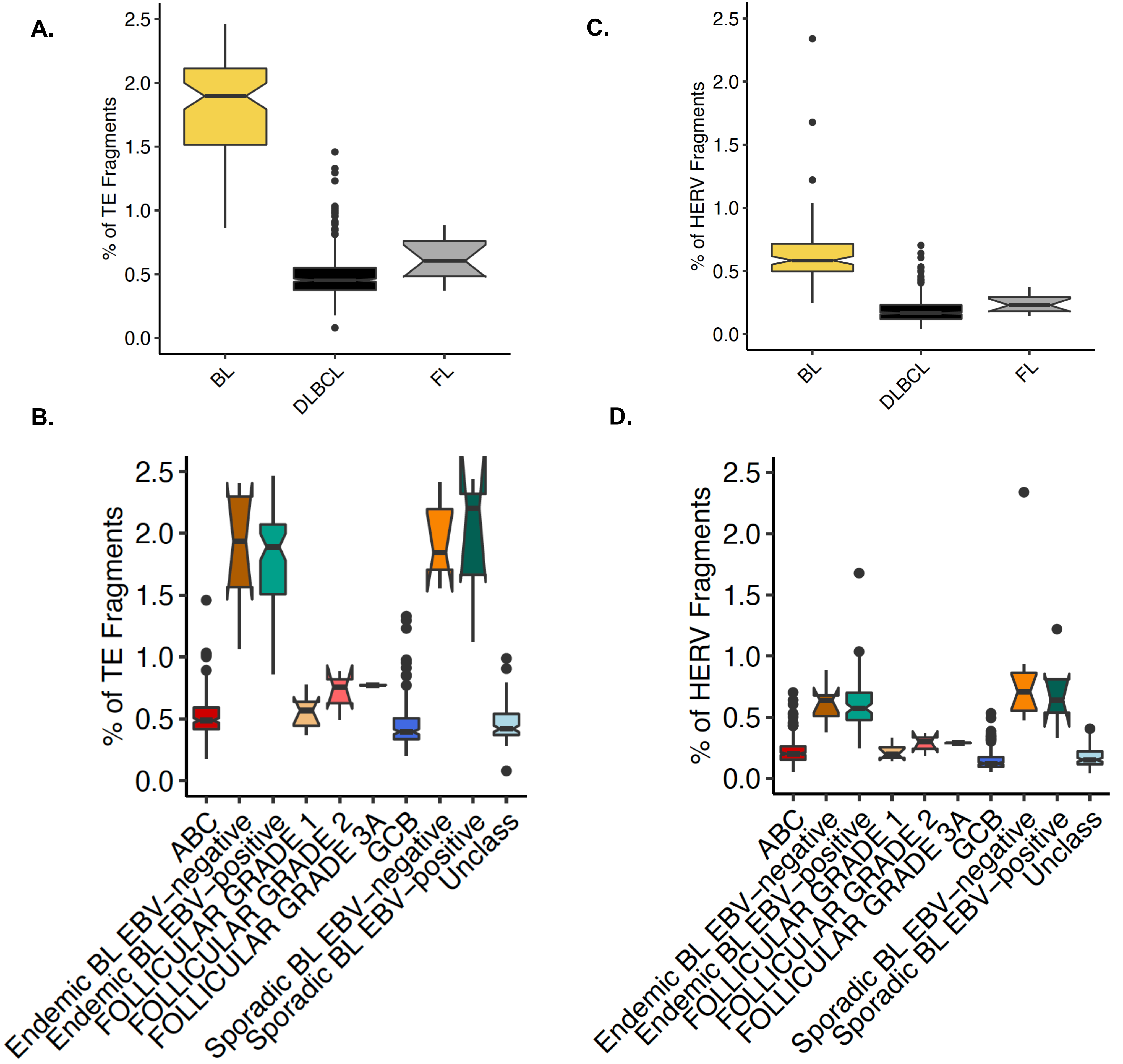
A.
% of TE Fragments
0.0
0.5
1.0
1.5
2.0
2.5
BL
DLBCL
FL
C.
% of HERV Fragments
0.0
0.5
1.0
1.5
2.0
2.5
BL
DLBCL
FL
B.
% of TE Fragments
0.0
0.5
1.0
1.5
2.0
2.5
ABC
Endemic BL EBV−negative
Endemic BL EBV−positive
FOLLICULAR GRADE 1
FOLLICULAR GRADE 2
FOLLICULAR GRADE 3A
GCB
Sporadic BL EBV−negative
Sporadic BL EBV−positive
Unclass
D.
% of HERV Fragments
0.0
0.5
1.0
1.5
2.0
2.5
ABC
Endemic BL EBV−negative
Endemic BL EBV−positive
FOLLICULAR GRADE 1
FOLLICULAR GRADE 2
FOLLICULAR GRADE 3A
GCB
Sporadic BL EBV−negative
Sporadic BL EBV−positive
Unclass

Supp Fig. 6

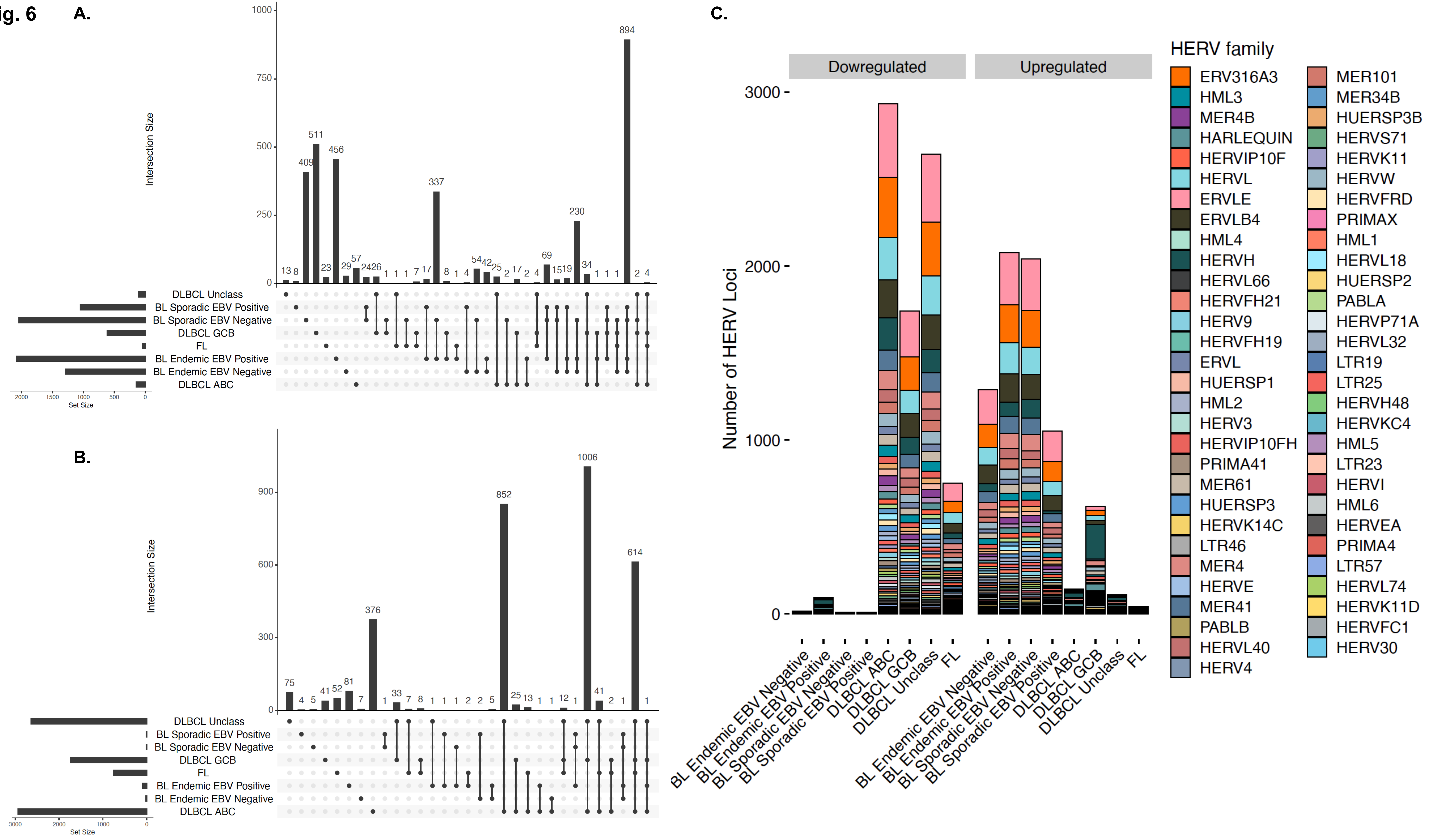
A.
Intersection Size
DLBCL Unclass
BL Sporadic EBV Positive
BL Sporadic EBV Negative
DLBCL GCB
FL
BL Endemic EBV Positive
BL Endemic EBV Negative
DLBCL ABC
Set Size
B.
C.
Dowregulated
Upregulated
Number of HERV Loci
HERV family
ERV316A3
HML3
MER4B
HARLEQUIN
HERVIP10F
HERVL
ERVLE
ERVLB4
HML4
HERVH
HERVL66
HERVFH21
HERV9
HERVFH19
ERVL
HUERSP1
HML2
HERV3
HERVIP10FH
PRIMA41
MER61
HUERSP3
HERVK14C
LTR46
MER4
HERVE
MER41
PABLB
HERVL40
HERV4
MER101
MER34B
HUERSP3B
HERVS71
HERVK11
HERVW
HERVFRD
PRIMAX
HML1
HERVL18
HUERSP2
PABLA
HERVP71A
HERVL32
LTR19
LTR25
HERVH48
HERVKC4
HML5
LTR23
HERVI
HML6
HERVEA
PRIMA4
LTR57
HERVL74
HERVK11D
HERVFC1
HERV30

Supp Fig. 7

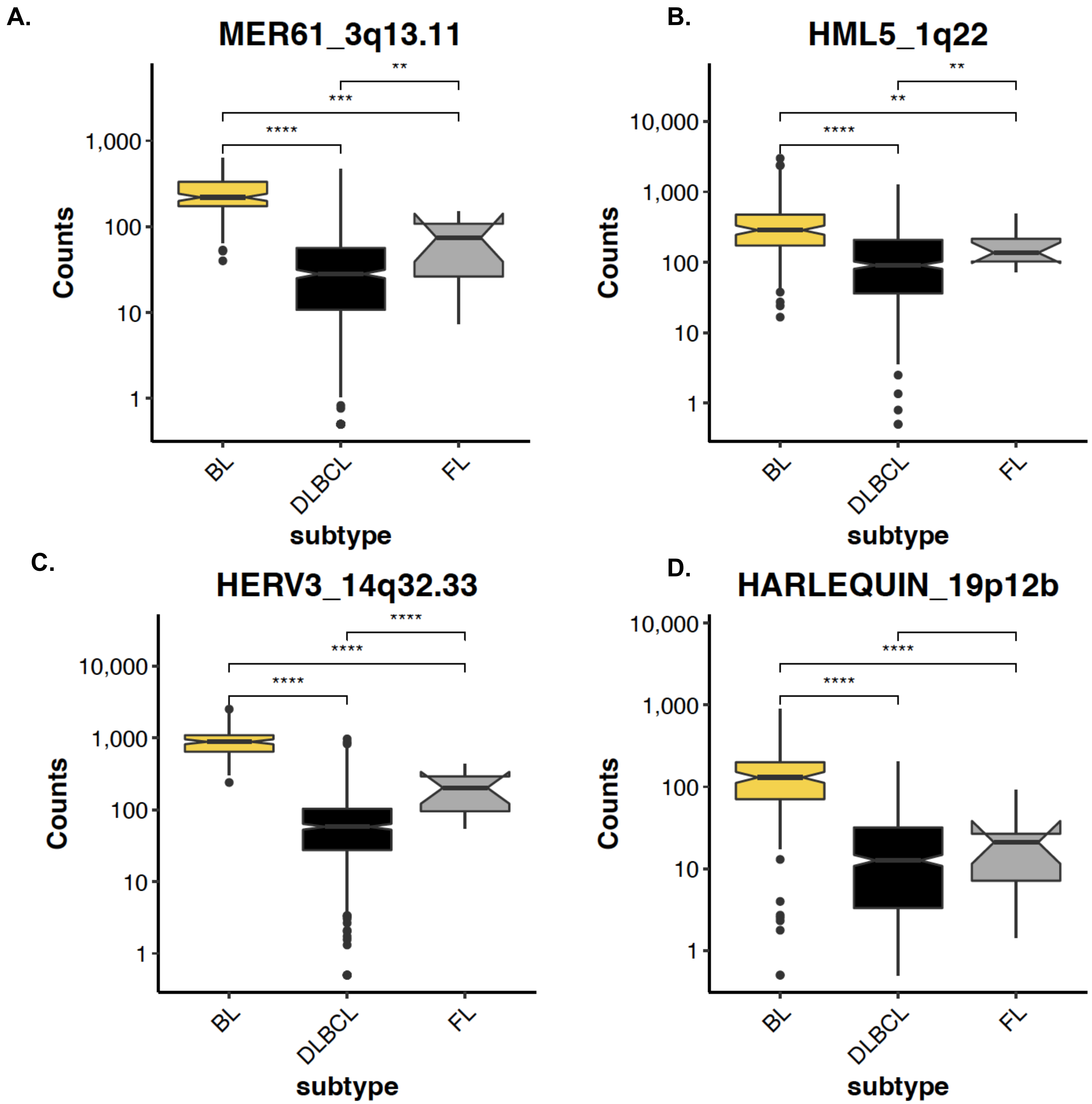
A.
MER61_3q13.11
**
***
****
1,000
100
10
1
Counts
BL
DLBCL
FL
subtype
B.
HML5_1q22
**
**
****
10,000
1,000
100
10
1
Counts
BL
DLBCL
FL
subtype
C.
HERV3_14q32.33
****
****
****
10,000
1,000
100
10
1
Counts
BL
DLBCL
FL
subtype
D.
HARLEQUIN_19p12b
****
****
10,000
1,000
100
10
1
Counts
BL
DLBCL
FL
subtype

Supp Fig. 8

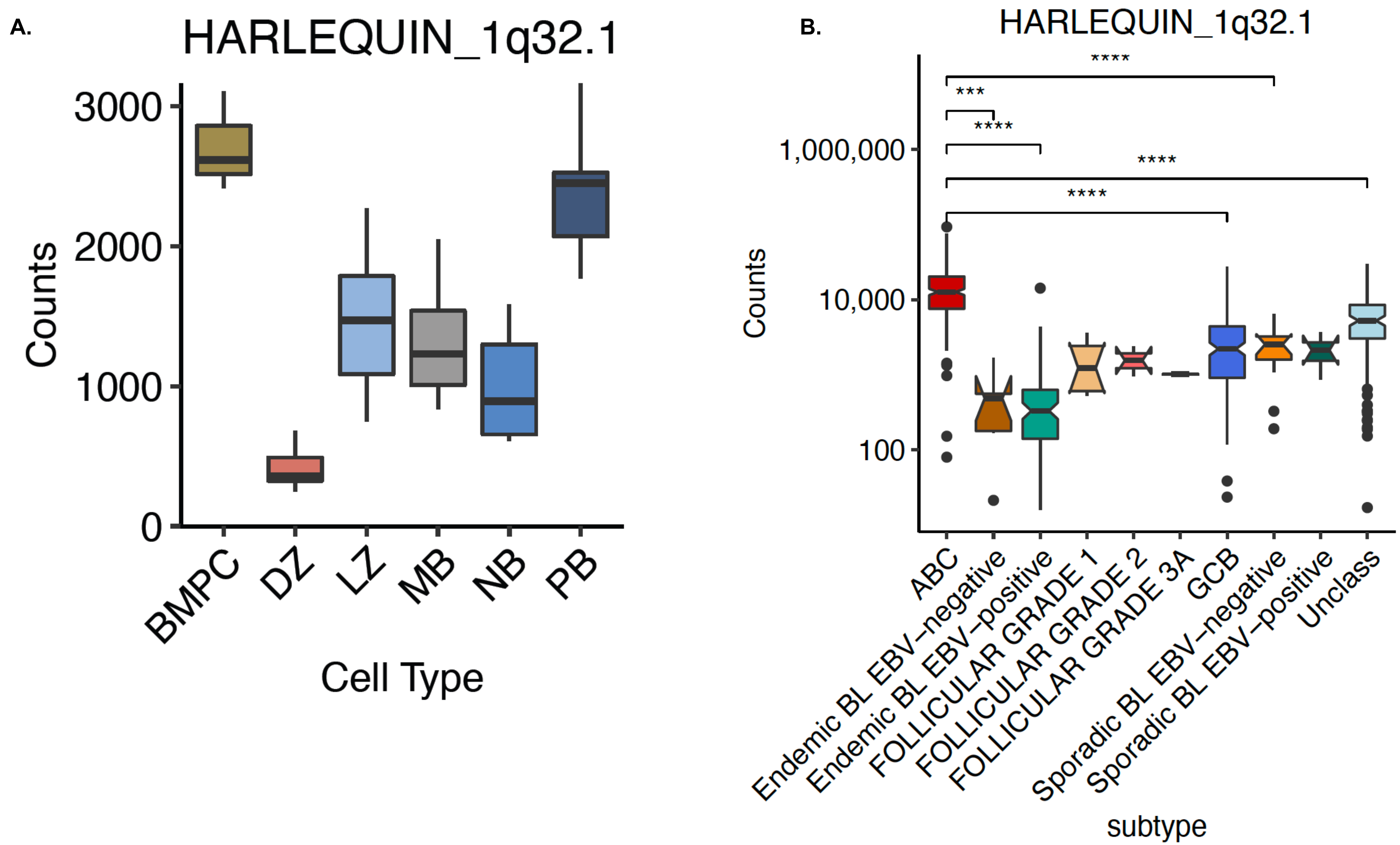
A.
HARLEQUIN_1q32.1
Counts
3000
2000
1000
0
BMPC
DZ
LZ
MB
NB
PB
Cell Type
B.
HARLEQUIN_1q32.1
Counts
1,000,000
10,000
100
****
***
****
****
****
ABC
Endemic BL EBV−negative
Endemic BL EBV−positive
FOLLICULAR GRADE 1
FOLLICULAR GRADE 2
FOLLICULAR GRADE 3A
GCB
Sporadic BL EBV−negative
Sporadic BL EBV−positive
Unclass
subtype

### Supp Fig. 9

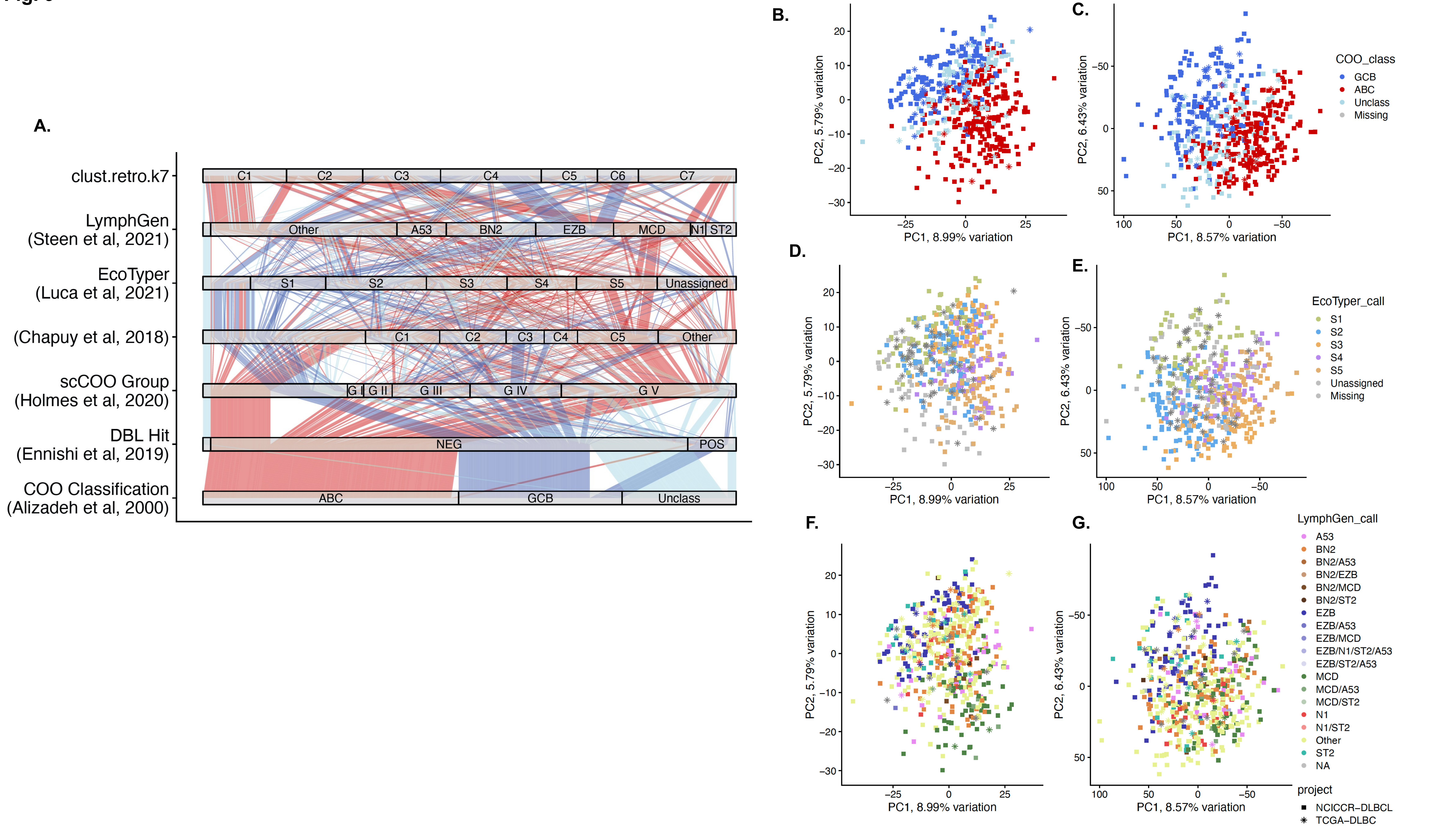

A.
clust.retro.k7
LymphGen (Steen et al, 2021)
EcoTyper (Luca et al, 2021)
(Chapuy et al, 2018)
scCOO Group (Holmes et al, 2020)
DBL Hit (Ennishi et al, 2019)
COO Classification (Alizadeh et al, 2000)
C1 C2 C3 C4 C5 C6 C7
Other A53 BN2 EZB MCD N1 ST2
S1 S2 S3 S4 S5 Unassigned
C1 C2 C3 C4 C5 Other
G I G II G III G IV G V
NEG POS
ABC GCB Unclass
B.
C.
D.
E.
F.
G.
PC1, 8.99% variation
PC2, 5.79% variation
PC1, 8.57% variation
PC2, 6.43% variation
COO_class
GCB
ABC
Unclass
Missing
EcoTyper_call
S1
S2
S3
S4
S5
Unassigned
Missing
LymphGen_call
A53
BN2
BN2/A53
BN2/EZB
BN2/MCD
BN2/ST2
EZB
EZB/A53
EZB/MCD
EZB/N1/ST2/A53
EZB/ST2/A53
MCD
MCD/A53
MCD/ST2
N1
N1/ST2
Other
ST2
NA
project
NCICCR–DLBCL
TCGA–DLBC

Supp Fig. 10

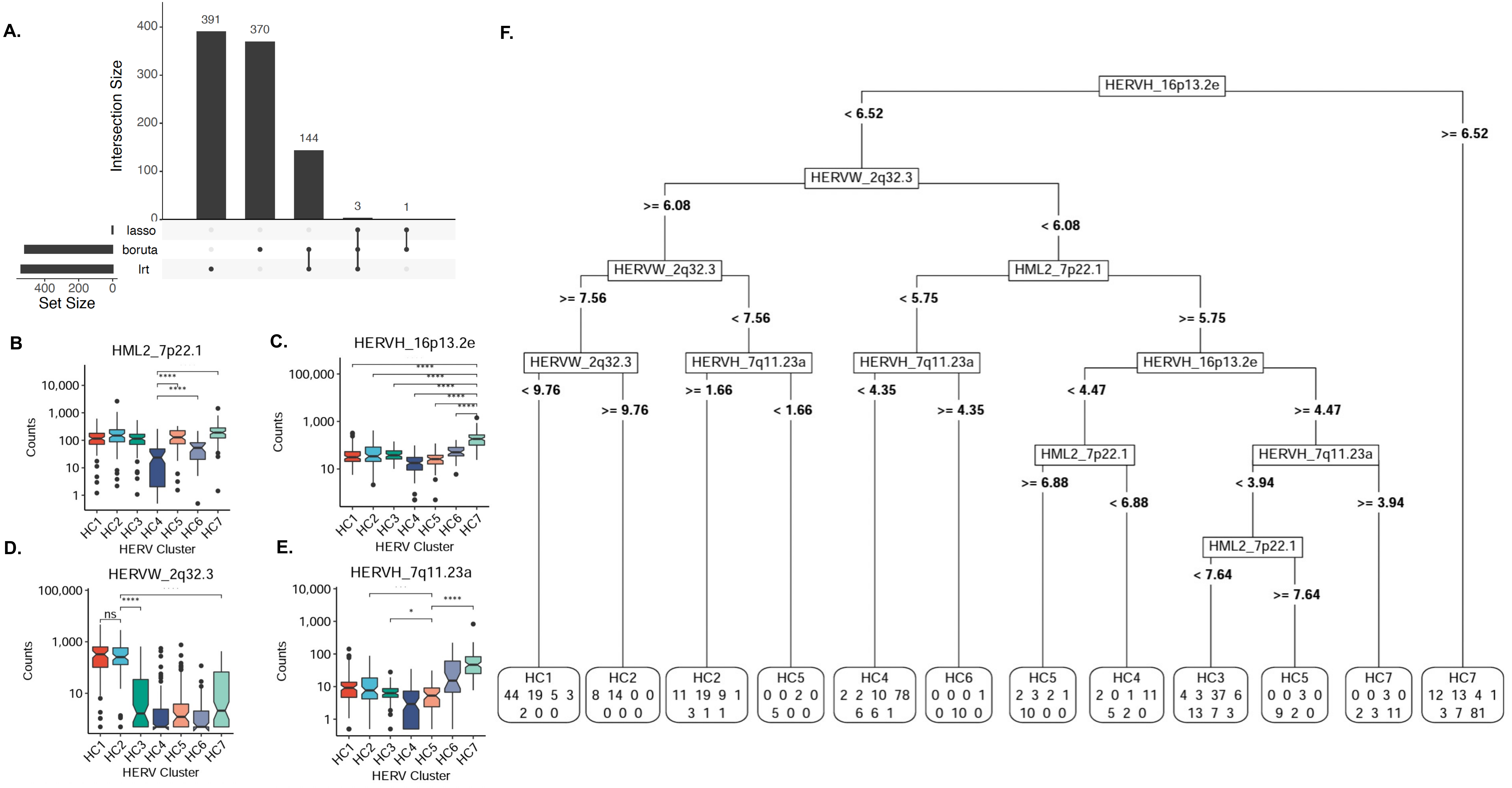

A.
Intersection Size
391
370
144
3
1
lasso
boruta
lrt
Set Size
B
HML2_7p22.1
Counts
HERV Cluster
C.
HERVH_16p13.2e
D.
HERVW_2q32.3
E.
HERVH_7q11.23a
F.
HERVH_16p13.2e
< 6.52
>= 6.52
HERVW_2q32.3
>= 6.08
< 6.08
HERVW_2q32.3
HML2_7p22.1
>= 7.56
< 7.56
< 5.75
>= 5.75
HERVW_2q32.3
HERVH_7q11.23a
HERVH_7q11.23a
HERVH_16p13.2e
< 9.76
>= 9.76
>= 1.66
< 1.66
< 4.35
>= 4.35
< 4.47
>= 4.47
HML2_7p22.1
HERVH_7q11.23a
>= 6.88
< 6.88
< 3.94
>= 3.94
HML2_7p22.1
< 7.64
>= 7.64
HC1 44 19 5 3 2 0 0
HC2 8 14 0 0 0 0 0
HC2 11 19 9 1 3 1 1
HC5 0 0 2 0 5 0 0
HC4 2 2 10 78 6 6 1
HC6 0 0 0 1 0 10 0
HC5 2 3 2 1 10 0 0
HC4 2 0 1 11 5 2 0
HC3 4 3 37 6 13 7 3
HC5 0 0 3 0 9 2 0
HC7 0 0 3 0 2 3 11
HC7 12 13 4 1 3 7 81

### Supp Fig. 11

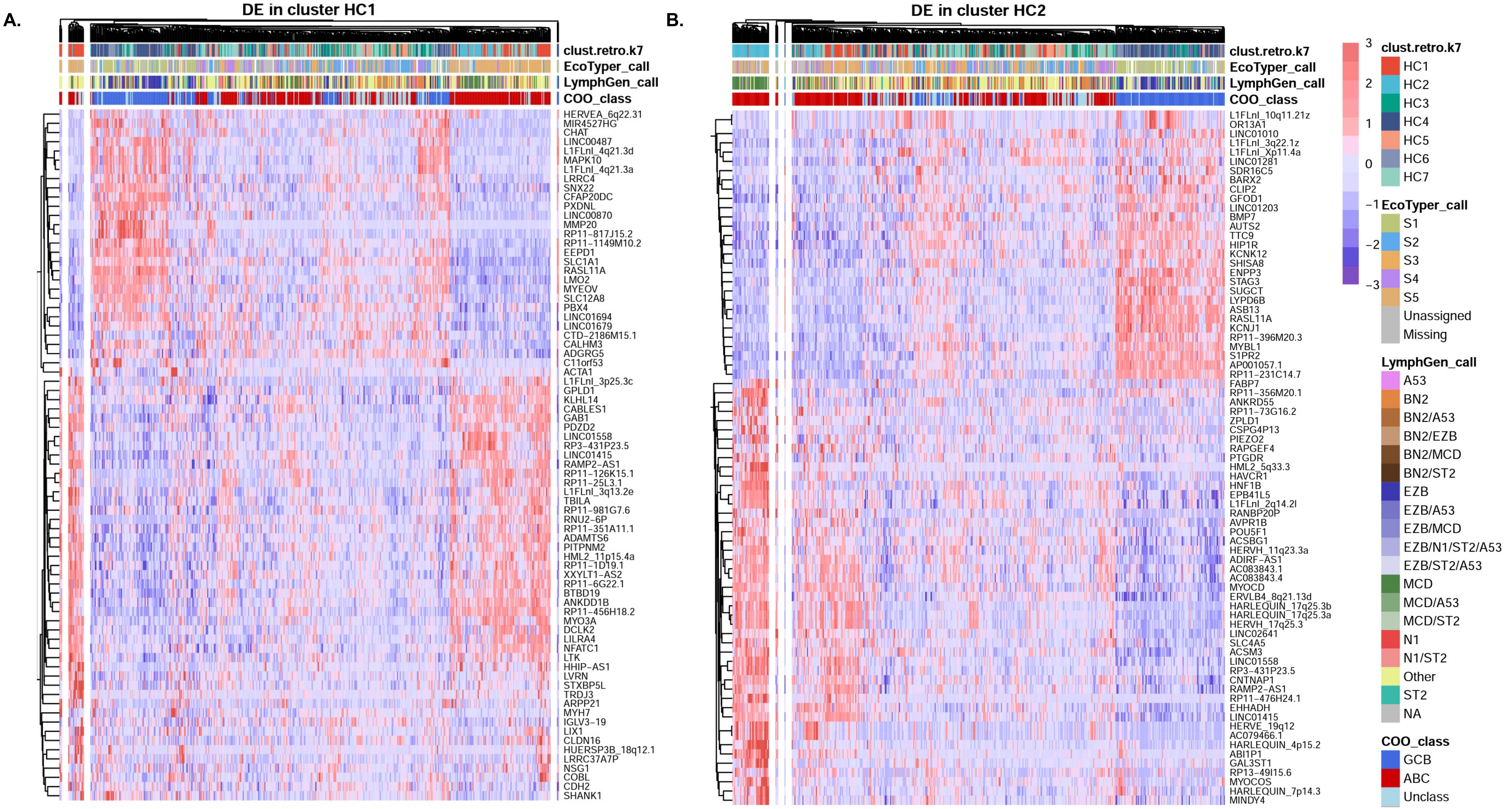
A.
DE in cluster HC1
clust.retro.k7
EcoTyper_call
LymphGen_call
COO_class
HERVEA_6q22.31
MIR4527HG
CHAT
LINC00487
L1FLnI_4q21.3d
MAPK10
L1FLnI_4q21.3a
LRRC4
SNX22
CFAP20DC
PXDNL
LINC00870
MMP20
RP11−817J15.2
RP11−1149M10.2
EEPD1
SLC1A1
RASL11A
LMO2
MYEOV
SLC12A8
PBX4
LINC01694
LINC01679
CTD−2186M15.1
CALHM3
ADGRG5
C11orf53
ACTA1
L1FLnI_3p25.3c
GPLD1
KLHL14
CABLES1
GAB1
PDZD2
LINC01558
RP3−431P23.5
LINC01415
RAMP2−AS1
RP11−126K15.1
RP11−25L3.1
L1FLnI_3q13.2e
TBILA
RP11−981G7.6
RNU2−6P
RP11−351A11.1
ADAMTS6
PITPNM2
HML2_11p15.4a
RP11−1D19.1
XXYLT1−AS2
RP11−6G22.1
BTBD19
ANKDD1B
RP11−456H18.2
MYO3A
DCLK2
LILRA4
NFATC1
LTK
HHIP−AS1
LVRN
STXBP5L
TRDJ3
ARPP21
MYH7
IGLV3−19
LIX1
CLDN16
HUERSP3B_18q12.1
LRRC37A7P
NSG1
COBL
CDH2
SHANK1
B.
DE in cluster HC2
clust.retro.k7
EcoTyper_call
LymphGen_call
COO_class
L1FLnI_10q11.21z
OR13A1
LINC01010
L1FLnI_3q22.1z
L1FLnI_Xp11.4a
LINC01281
SDR16C5
BARX2
CLIP2
GFOD1
LINC01203
BMP7
AUTS2
TTC9
HIP1R
KCNK12
SHISA8
ENPP3
STAG3
SUGCT
LYPD6B
ASB13
RASL11A
KCNJ1
RP11−396M20.3
MYBL1
S1PR2
AP001057.1
RP11−231C14.7
FABP7
RP11−356M20.1
ANKRD55
RP11−73G16.2
ZPLD1
CSPG4P13
PIEZO2
RAPGEF4
PTGDR
HML2_5q33.3
HAVCR1
HNF1B
EPB41L5
L1FLnI_2q14.2l
RANBP20P
AVPR1B
POU5F1
ACSBG1
HERVH_11q23.3a
ADIRF−AS1
AC083843.1
AC083843.4
MYOCD
ERVLB4_8q21.13d
HARLEQUIN_17q25.3b
HARLEQUIN_17q25.3a
HERVH_17q25.3
LINC02641
SLC4A5
ACSM3
LINC01558
RP3−431P23.5
CNTNAP1
RAMP2−AS1
RP11−476H24.1
EHHADH
LINC01415
HERVE_19q12
AC079466.1
HARLEQUIN_4p15.2
ABI1P1
GAL3ST1
RP13−49I15.6
MYOCOS
HARLEQUIN_7p14.3
MINDY4
3
2
1
0
−1
−2
−3
clust.retro.k7
HC1
HC2
HC3
HC4
HC5
HC6
HC7
EcoTyper_call
S1
S2
S3
S4
S5
Unassigned
Missing
LymphGen_call
A53
BN2
BN2/A53
BN2/EZB
BN2/MCD
BN2/ST2
EZB
EZB/A53
EZB/MCD
EZB/N1/ST2/A53
EZB/ST2/A53
MCD
MCD/A53
MCD/ST2
N1
N1/ST2
Other
ST2
NA
COO_class
GCB
ABC
Unclass

Supp Fig. 12

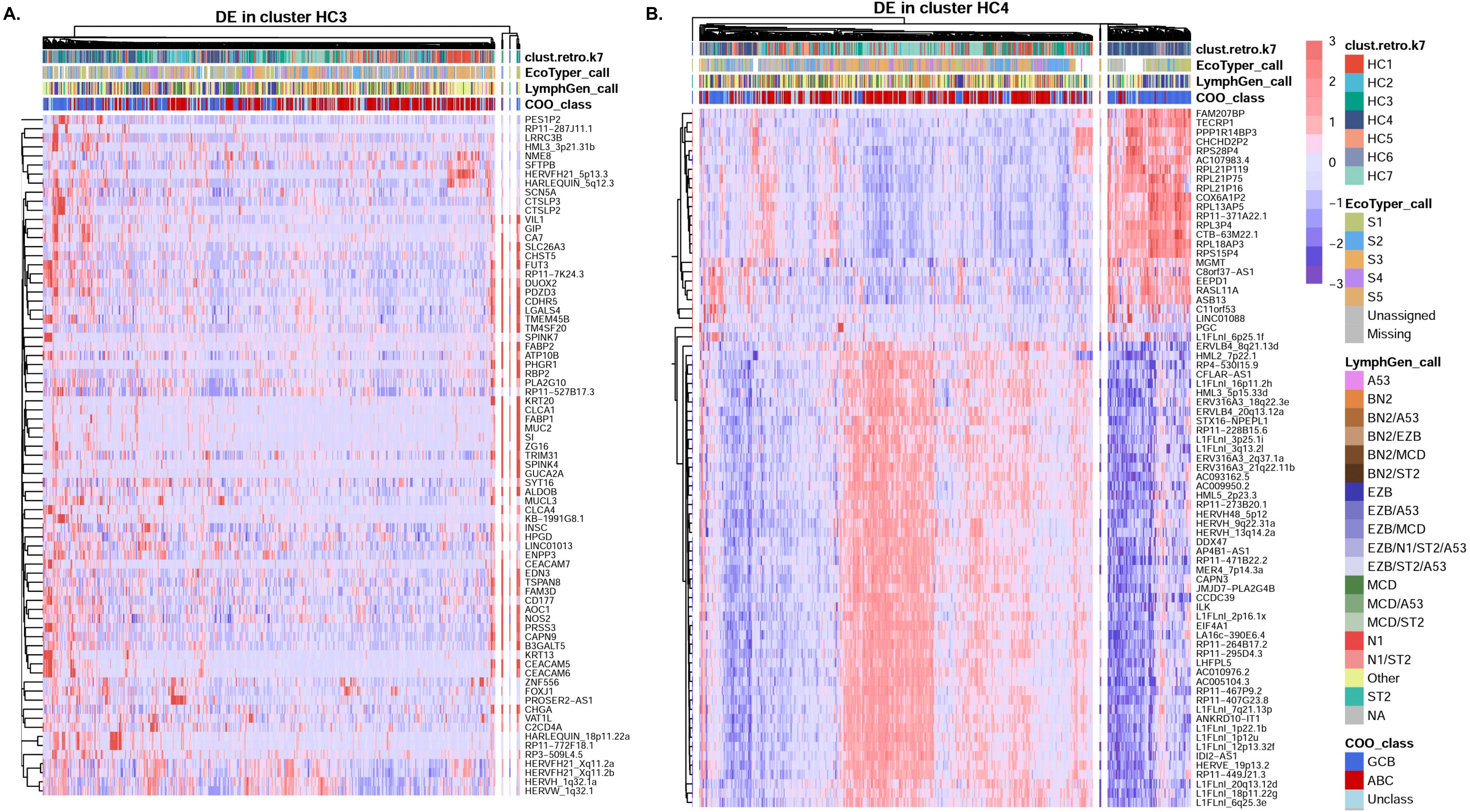

A.
DE in cluster HC3
clust.retro.k7
EcoTyper_call
LymphGen_call
COO_class
PES1P2
RP11-287J11.1
LRRC3B
HML3_3p21.31b
NME8
SFTPB
HERVFH21_5p13.3
HARLEQUIN_5q12.3
SCN5A
CTSLP3
CTSLP2
VIL1
GIP
CA7
SLC26A3
CHST5
FUT3
RP11-7K24.3
DUOX2
PDZD3
CDHR5
LGALS4
TMEM45B
TM4SF20
SPINK7
FABP2
ATP10B
PHGR1
RBP2
PLA2G10
RP11-527B17.3
KRT20
CLCA1
FABP1
MUC2
SI
ZG16
TRIM31
SPINK4
GUCA2A
SYT16
ALDOB
MUCL3
CLCA4
KB-1991G8.1
INSC
HPGD
LINC01013
ENPP3
CEACAM7
EDN3
TSPAN8
FAM3D
CD177
AOC1
NOS2
PRSS3
CAPN9
B3GALT5
KRT13
CEACAM5
CEACAM6
ZNF556
FOXJ1
PROSER2-AS1
CHGA
VAT1L
C2CD4A
HARLEQUIN_18p11.22a
RP11-772F18.1
RP3-509L4.5
HERVFH21_Xq11.2a
HERVFH21_Xq11.2b
HERVH_1q32.1a
HERVW_1q32.1
B.
DE in cluster HC4
clust.retro.k7
EcoTyper_call
LymphGen_call
COO_class
FAM207BP
TECRP1
PPP1R14BP3
CHCHD2P2
RPS28P4
AC107983.4
RPL21P119
RPL21P75
RPL21P16
COX6A1P2
RPL13AP5
RP11-371A22.1
RPL3P4
CTB-63M22.1
RPL18AP3
RPS15P4
MGMT
C8orf37-AS1
EEPD1
RASL11A
ASB13
C11orf53
LINC01088
PGC
L1FLnI_6p25.1f
ERVLB4_8q21.13d
HML2_7p22.1
RP4-530I15.9
CFLAR-AS1
L1FLnI_16p11.2h
HML3_5p15.33d
ERV316A3_18q22.3e
ERVLB4_20q13.12a
STX16-NPEPL1
RP11-228B15.6
L1FLnI_3p25.1i
L1FLnI_3q13.2l
ERV316A3_2q37.1a
ERV316A3_21q22.11b
AC093162.5
AC009950.2
HML5_2p23.3
RP11-273B20.1
HERVH48_5p12
HERVH_9q22.31a
HERVH_13q14.2a
DDX47
AP4B1-AS1
RP11-471B22.2
MER4_7p14.3a
CAPN3
JMJD7-PLA2G4B
CCDC39
ILK
L1FLnI_2p16.1x
EIF4A1
LA16c-390E6.4
RP11-264B17.2
RP11-295D4.3
LHFPL5
AC010976.2
AC005104.3
RP11-467P9.2
RP11-407G23.8
L1FLnI_7q21.13p
ANKRD10-IT1
L1FLnI_1p22.1b
L1FLnI_1p12u
L1FLnI_12p13.32f
IDI2-AS1
HERVE_19p13.2
RP11-449J21.3
L1FLnI_20q13.12d
L1FLnI_18p11.22g
L1FLnI_6q25.3e
3
2
1
0
-1
-2
-3
clust.retro.k7
HC1
HC2
HC3
HC4
HC5
HC6
HC7
EcoTyper_call
S1
S2
S3
S4
S5
Unassigned
Missing
LymphGen_call
A53
BN2
BN2/A53
BN2/EZB
BN2/MCD
BN2/ST2
EZB
EZB/A53
EZB/MCD
EZB/N1/ST2/A53
EZB/ST2/A53
MCD
MCD/A53
MCD/ST2
N1
N1/ST2
Other
ST2
NA
COO_class
GCB
ABC
Unclass

**Supp Fig. 13**

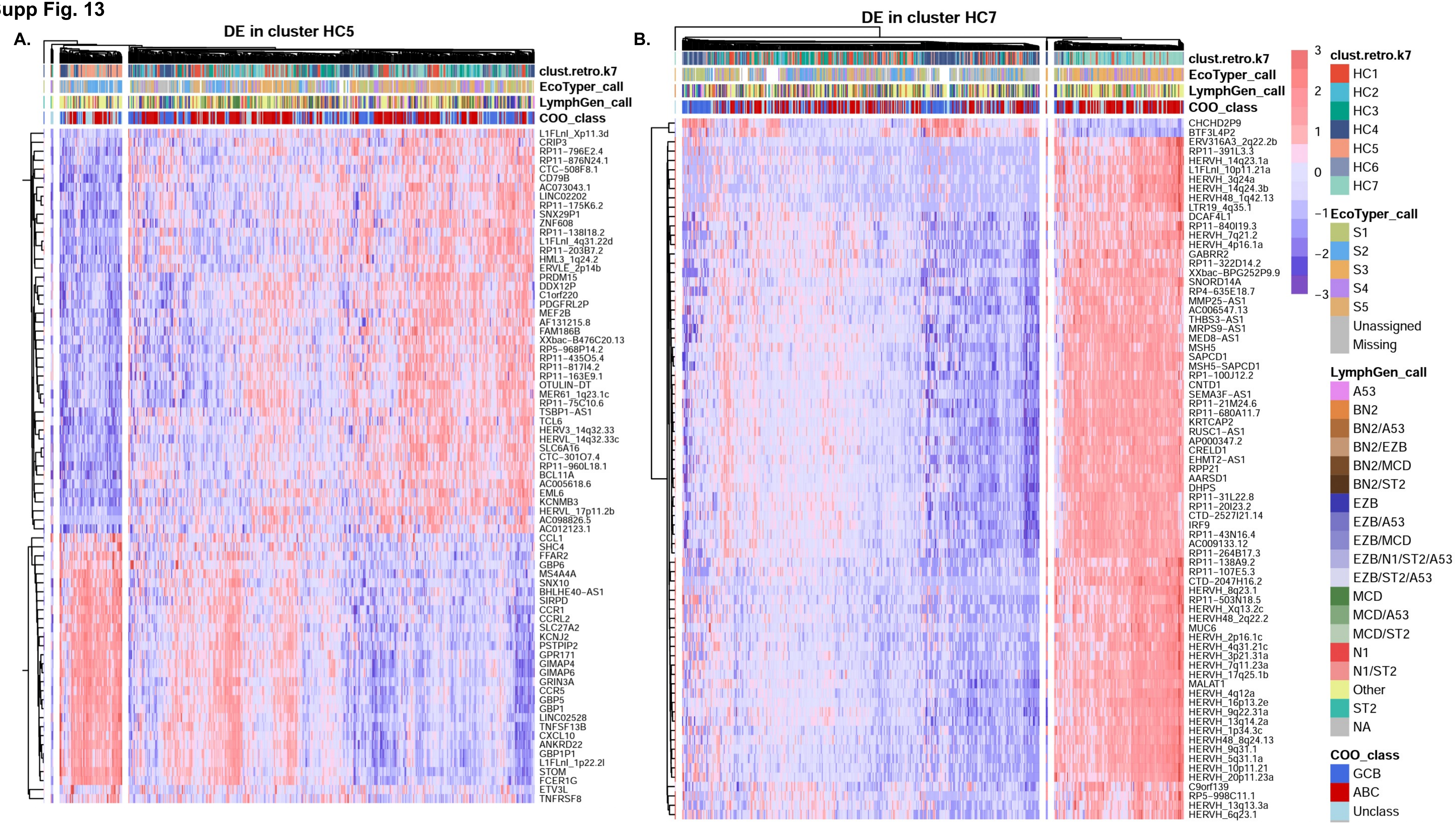
A.
DE in cluster HC5
clust.retro.k7
EcoTyper_call
LymphGen_call
COO_class
L1FLnI_Xp11.3d
CRIP3
RP11−796E2.4
RP11−876N24.1
CTC−508F8.1
CD79B
AC073043.1
LINC02202
RP11−175K6.2
SNX29P1
ZNF608
RP11−138I18.2
L1FLnI_4q31.22d
RP11−203B7.2
HML3_1q24.2
ERVLE_2p14b
PRDM15
DDX12P
C1orf220
PDGFRL2P
MEF2B
AF131215.8
FAM186B
XXbac−B476C20.13
RP5−968P14.2
RP11−435O5.4
RP11−817I4.2
RP11−163E9.1
OTULIN−DT
MER61_1q23.1c
RP11−75C10.6
TSBP1−AS1
TCL6
HERV3_14q32.33
HERVL_14q32.33c
SLC6A16
CTC−301O7.4
RP11−960L18.1
BCL11A
AC005618.6
EML6
KCNMB3
HERVL_17p11.2b
AC098826.5
AC012123.1
CCL1
SHC4
FFAR2
GBP6
MS4A4A
SNX10
BHLHE40−AS1
SIRPD
CCR1
CCRL2
SLC27A2
KCNJ2
PSTPIP2
GPR171
GIMAP4
GIMAP6
GRIN3A
CCR5
GBP5
GBP1
LINC02528
TNFSF13B
CXCL10
ANKRD22
GBP1P1
L1FLnI_1p22.2l
STOM
FCER1G
ETV3L
TNFRSF8
B.
DE in cluster HC7
clust.retro.k7
EcoTyper_call
LymphGen_call
COO_class
CHCHD2P9
BTF3L4P2
ERV316A3_2q22.2b
RP11−391L3.3
HERVH_14q23.1a
L1FLnI_10p11.21a
HERVH_3q24a
HERVH_14q24.3b
HERVH48_1q42.13
LTR19_4q35.1
DCAF4L1
RP11−840I19.3
HERVH_7q21.2
HERVH_4p16.1a
GABRR2
RP11−322D14.2
XXbac−BPG252P9.9
SNORD14A
RP4−635E18.7
MMP25−AS1
AC006547.13
THBS3−AS1
MRPS9−AS1
MED8−AS1
MSH5
SAPCD1
MSH5−SAPCD1
RP1−100J12.2
CNTD1
SEMA3F−AS1
RP11−21M24.6
RP11−680A11.7
KRTCAP2
RUSC1−AS1
AP000347.2
CRELD1
EHMT2−AS1
RPP21
AARSD1
DHPS
RP11−31L22.8
RP11−20I23.2
CTD−2527I21.14
IRF9
RP11−43N16.4
AC009133.12
RP11−264B17.3
RP11−138A9.2
RP11−107E5.3
CTD−2047H16.2
HERVH_8q23.1
RP11−503N18.5
HERVH_Xq13.2c
HERVH48_2q22.2
MUC6
HERVH_2p16.1c
HERVH_4q31.21c
HERVH_3p21.31a
HERVH_7q11.23a
HERVH_17q25.1b
MALAT1
HERVH_4q12a
HERVH_16p13.2e
HERVH_9q22.31a
HERVH_13q14.2a
HERVH_1p34.3c
HERVH48_8q24.13
HERVH_9q31.1
HERVH_5q31.1a
HERVH_10p11.21
HERVH_20p11.23a
C9orf139
RP5−998C11.1
HERVH_13q13.3a
HERVH_6q23.1
3
2
1
0
−1
−2
−3
clust.retro.k7
HC1
HC2
HC3
HC4
HC5
HC6
HC7
EcoTyper_call
S1
S2
S3
S4
S5
Unassigned
Missing
LymphGen_call
A53
BN2
BN2/A53
BN2/EZB
BN2/MCD
BN2/ST2
EZB
EZB/A53
EZB/MCD
EZB/N1/ST2/A53
EZB/ST2/A53
MCD
MCD/A53
MCD/ST2
N1
N1/ST2
Other
ST2
NA
COO_class
GCB
ABC
Unclass

Supp Fig. 14

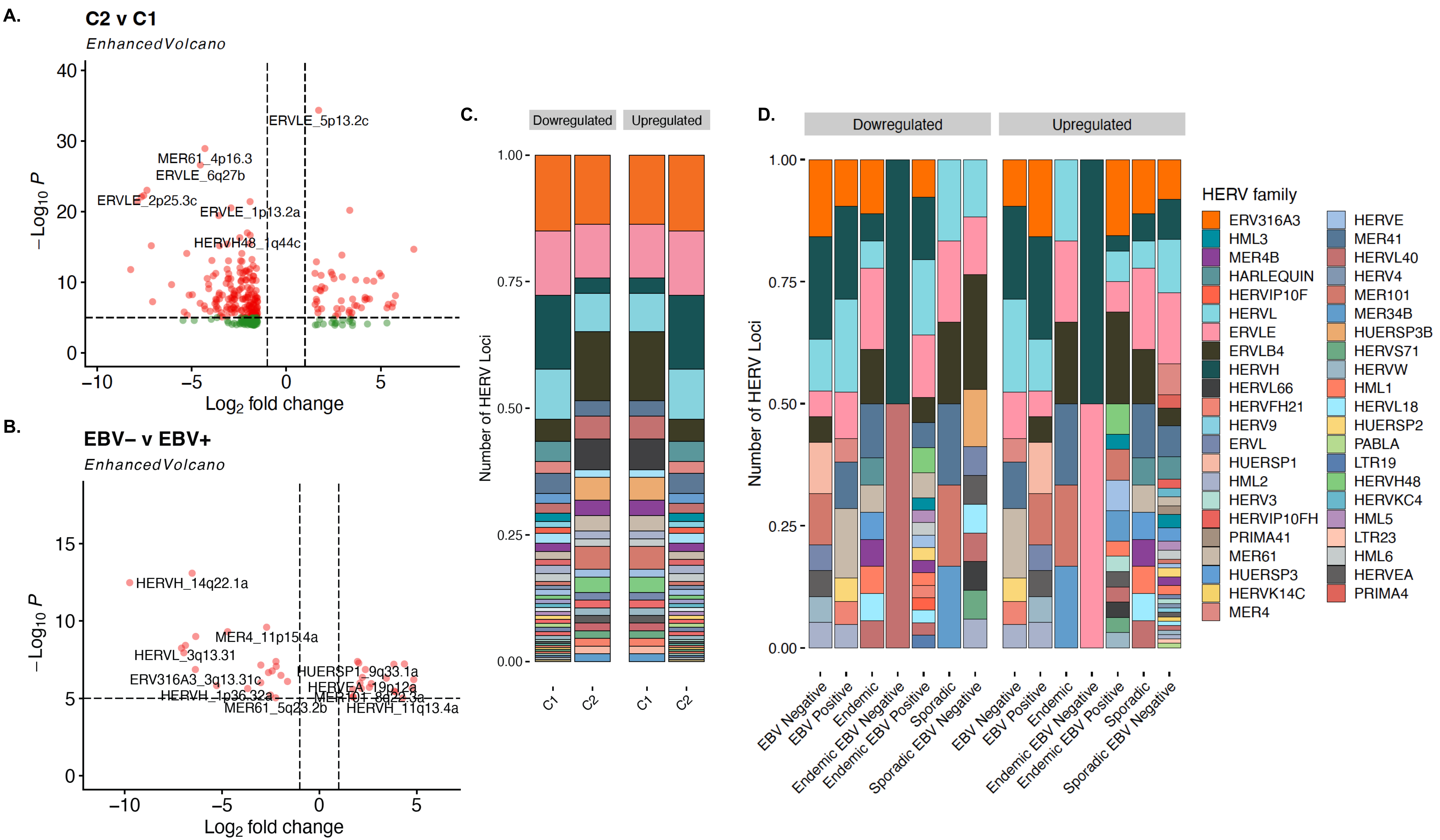

A.
C2 v C1
EnhancedVolcano
ERVLE_5p13.2c
MER61_4p16.3
ERVLE_6q27b
ERVLE_2p25.3c
ERVLE_1p13.2a
HERVH48_1q44c
$-Log_{10}$ P
$Log_2$ fold change
B.
EBV– v EBV+
EnhancedVolcano
HERVH_14q22.1a
MER4_11p15.4a
HERVL_3q13.31
ERV316A3_3q13.31c
HERVH_1p36.32a
MER61_5q23.2b
HUERSP1_9q33.1a
HERVEA_19p12a
MER101_8q22.3a
HERVH_11q13.4a
C.
Dowregulated
Upregulated
Number of HERV Loci
C1
C2
D.
Dowregulated
Upregulated
Number of HERV Loci
EBV Negative
EBV Positive
Endemic
Endemic EBV Negative
Endemic EBV Positive
Sporadic
Sporadic EBV Negative
HERV family
ERV316A3
HML3
MER4B
HARLEQUIN
HERVIP10F
HERVL
ERVLE
ERVLB4
HERVH
HERVL66
HERVFH21
HERV9
ERVL
HUERSP1
HML2
HERV3
HERVIP10FH
PRIMA41
MER61
HUERSP3
HERVK14C
MER4
HERVE
MER41
HERVL40
HERV4
MER101
MER34B
HUERSP3B
HERVS71
HERVW
HML1
HERVL18
HUERSP2
PABLA
LTR19
HERVH48
HERVKC4
HML5
LTR23
HML6
HERVEA
PRIMA4

Supp Fig. 15

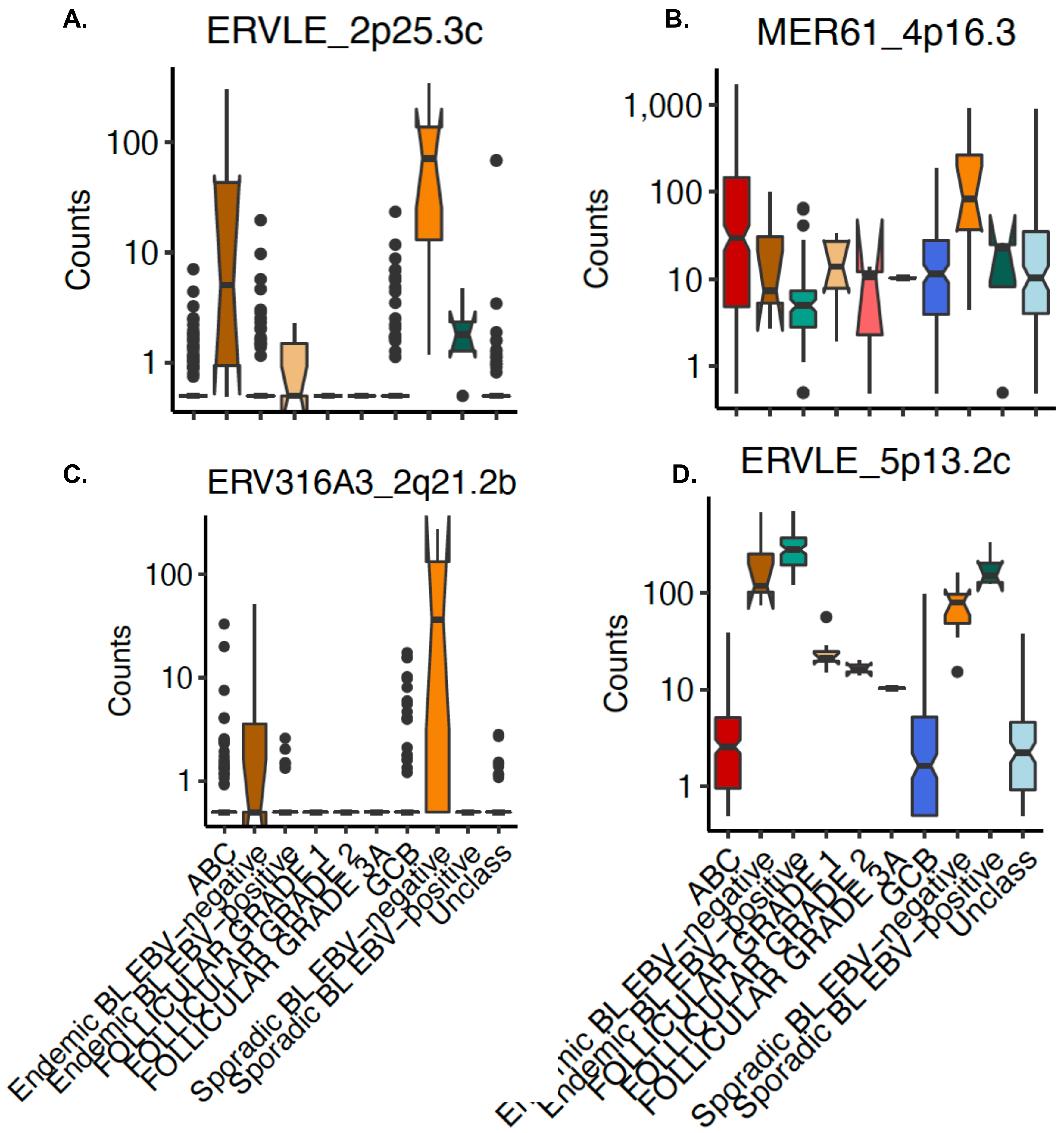

A.
ERVLE_2p25.3c
B.
MER61_4p16.3
C.
ERV316A3_2q21.2b
D.
ERVLE_5p13.2c
Counts
1
10
100
1,000
ABC
Endemic BL EBV-negative
Endemic BL EBV-positive
FOLLICULAR GRADE 1
FOLLICULAR GRADE 2
FOLLICULAR GRADE 3A
GCB
Sporadic BL EBV-negative
Sporadic BL EBV-positive
Unclass

**Supp Fig. 16**

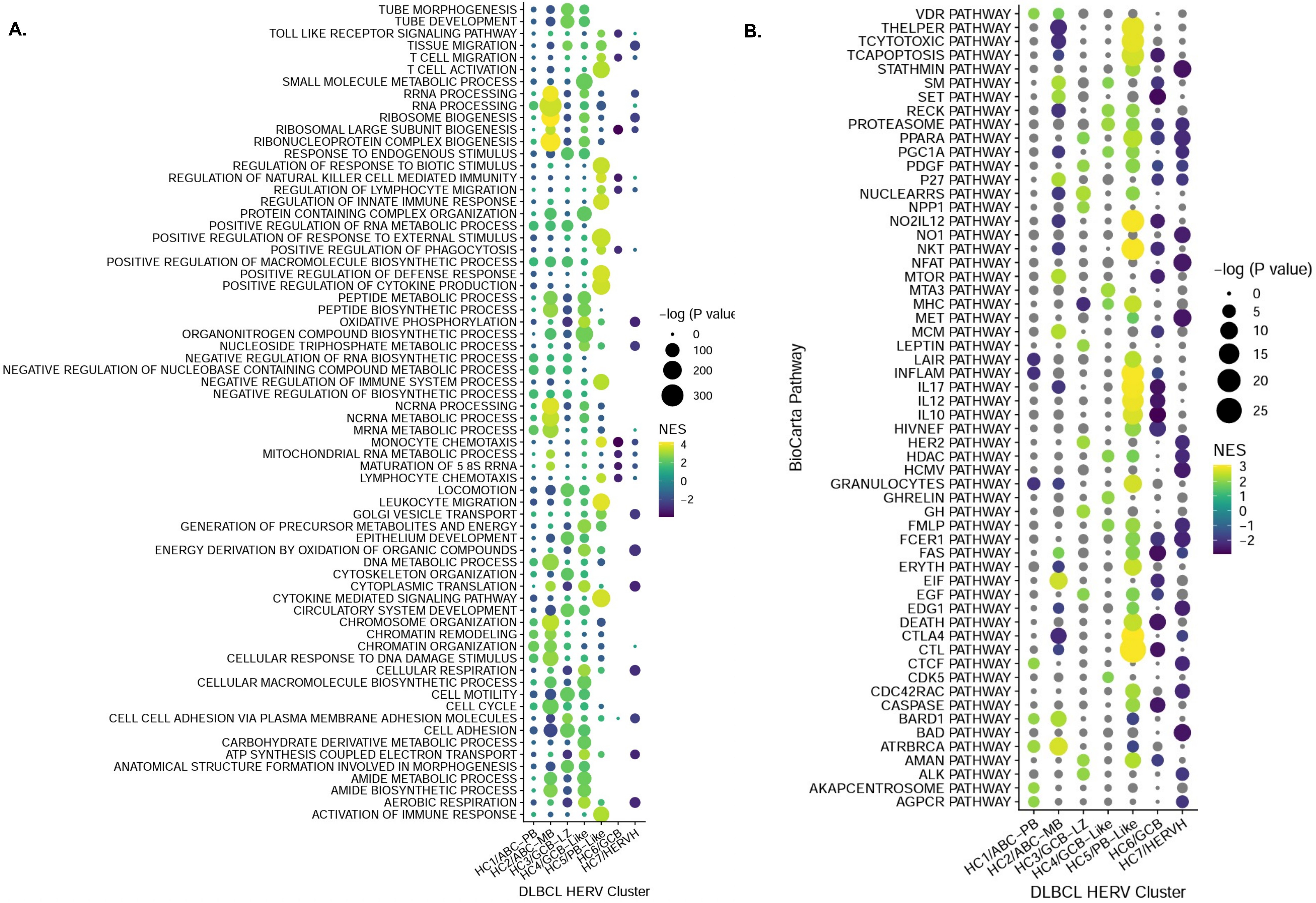
A.
GO BP Pathway
TUBE MORPHOGENESIS
TUBE DEVELOPMENT
TOLL LIKE RECEPTOR SIGNALING PATHWAY
TISSUE MIGRATION
T CELL MIGRATION
T CELL ACTIVATION
SMALL MOLECULE METABOLIC PROCESS
RRNA PROCESSING
RNA PROCESSING
RIBOSOME BIOGENESIS
RIBOSOMAL LARGE SUBUNIT BIOGENESIS
RIBONUCLEOPROTEIN COMPLEX BIOGENESIS
RESPONSE TO ENDOGENOUS STIMULUS
REGULATION OF RESPONSE TO BIOTIC STIMULUS
REGULATION OF NATURAL KILLER CELL MEDIATED IMMUNITY
REGULATION OF LYMPHOCYTE MIGRATION
REGULATION OF INNATE IMMUNE RESPONSE
PROTEIN CONTAINING COMPLEX ORGANIZATION
POSITIVE REGULATION OF RNA METABOLIC PROCESS
POSITIVE REGULATION OF RESPONSE TO EXTERNAL STIMULUS
POSITIVE REGULATION OF PHAGOCYTOSIS
POSITIVE REGULATION OF MACROMOLECULE BIOSYNTHETIC PROCESS
POSITIVE REGULATION OF DEFENSE RESPONSE
POSITIVE REGULATION OF CYTOKINE PRODUCTION
PEPTIDE METABOLIC PROCESS
PEPTIDE BIOSYNTHETIC PROCESS
OXIDATIVE PHOSPHORYLATION
ORGANONITROGEN COMPOUND BIOSYNTHETIC PROCESS
NUCLEOSIDE TRIPHOSPHATE METABOLIC PROCESS
NEGATIVE REGULATION OF RNA BIOSYNTHETIC PROCESS
NEGATIVE REGULATION OF NUCLEOBASE CONTAINING COMPOUND METABOLIC PROCESS
NEGATIVE REGULATION OF IMMUNE SYSTEM PROCESS
NEGATIVE REGULATION OF BIOSYNTHETIC PROCESS
NCRNA PROCESSING
NCRNA METABOLIC PROCESS
MRNA METABOLIC PROCESS
MONOCYTE CHEMOTAXIS
MITOCHONDRIAL RNA METABOLIC PROCESS
MATURATION OF 5 8S RRNA
LYMPHOCYTE CHEMOTAXIS
LOCOMOTION
LEUKOCYTE MIGRATION
GOLGI VESICLE TRANSPORT
GENERATION OF PRECURSOR METABOLITES AND ENERGY
EPITHELIUM DEVELOPMENT
ENERGY DERIVATION BY OXIDATION OF ORGANIC COMPOUNDS
DNA METABOLIC PROCESS
CYTOSKELETON ORGANIZATION
CYTOPLASMIC TRANSLATION
CYTOKINE MEDIATED SIGNALING PATHWAY
CIRCULATORY SYSTEM DEVELOPMENT
CHROMOSOME ORGANIZATION
CHROMATIN REMODELING
CHROMATIN ORGANIZATION
CELLULAR RESPONSE TO DNA DAMAGE STIMULUS
CELLULAR RESPIRATION
CELLULAR MACROMOLECULE BIOSYNTHETIC PROCESS
CELL MOTILITY
CELL CYCLE
CELL CELL ADHESION VIA PLASMA MEMBRANE ADHESION MOLECULES
CELL ADHESION
CARBOHYDRATE DERIVATIVE METABOLIC PROCESS
ATP SYNTHESIS COUPLED ELECTRON TRANSPORT
ANATOMICAL STRUCTURE FORMATION INVOLVED IN MORPHOGENESIS
AMIDE METABOLIC PROCESS
AMIDE BIOSYNTHETIC PROCESS
AEROBIC RESPIRATION
ACTIVATION OF IMMUNE RESPONSE
HC1/ABC−PB
HC2/ABC−MB
HC3/GCB−LZ
HC4/GCB−Like
HC5/PB−Like
HC6/GCB
HC7/HERVH
DLBCL HERV Cluster
−log (P value
0
100
200
300
NES
4
2
0
−2
B.
BioCarta Pathway
VDR PATHWAY
THELPER PATHWAY
TCYTOTOXIC PATHWAY
TCAPOPTOSIS PATHWAY
STATHMIN PATHWAY
SM PATHWAY
SET PATHWAY
RECK PATHWAY
PROTEASOME PATHWAY
PPARA PATHWAY
PGC1A PATHWAY
PDGF PATHWAY
P27 PATHWAY
NUCLEARRS PATHWAY
NPP1 PATHWAY
NO2IL12 PATHWAY
NO1 PATHWAY
NKT PATHWAY
NFAT PATHWAY
MTOR PATHWAY
MTA3 PATHWAY
MHC PATHWAY
MET PATHWAY
MCM PATHWAY
LEPTIN PATHWAY
LAIR PATHWAY
INFLAM PATHWAY
IL17 PATHWAY
IL12 PATHWAY
IL10 PATHWAY
HIVNEF PATHWAY
HER2 PATHWAY
HDAC PATHWAY
HCMV PATHWAY
GRANULOCYTES PATHWAY
GHRELIN PATHWAY
GH PATHWAY
FMLP PATHWAY
FCER1 PATHWAY
FAS PATHWAY
ERYTH PATHWAY
EIF PATHWAY
EGF PATHWAY
EDG1 PATHWAY
DEATH PATHWAY
CTLA4 PATHWAY
CTL PATHWAY
CTCF PATHWAY
CDK5 PATHWAY
CDC42RAC PATHWAY
CASPASE PATHWAY
BARD1 PATHWAY
BAD PATHWAY
ATRBRCA PATHWAY
AMAN PATHWAY
ALK PATHWAY
AKAPCENTROSOME PATHWAY
AGPCR PATHWAY
HC1/ABC−PB
HC2/ABC−MB
HC3/GCB−LZ
HC4/GCB−Like
HC5/PB−Like
HC6/GCB
HC7/HERVH
DLBCL HERV Cluster
−log (P value)
0
5
10
15
20
25
NES
3
2
1
0
−1
−2

**Supp Fig. 17**

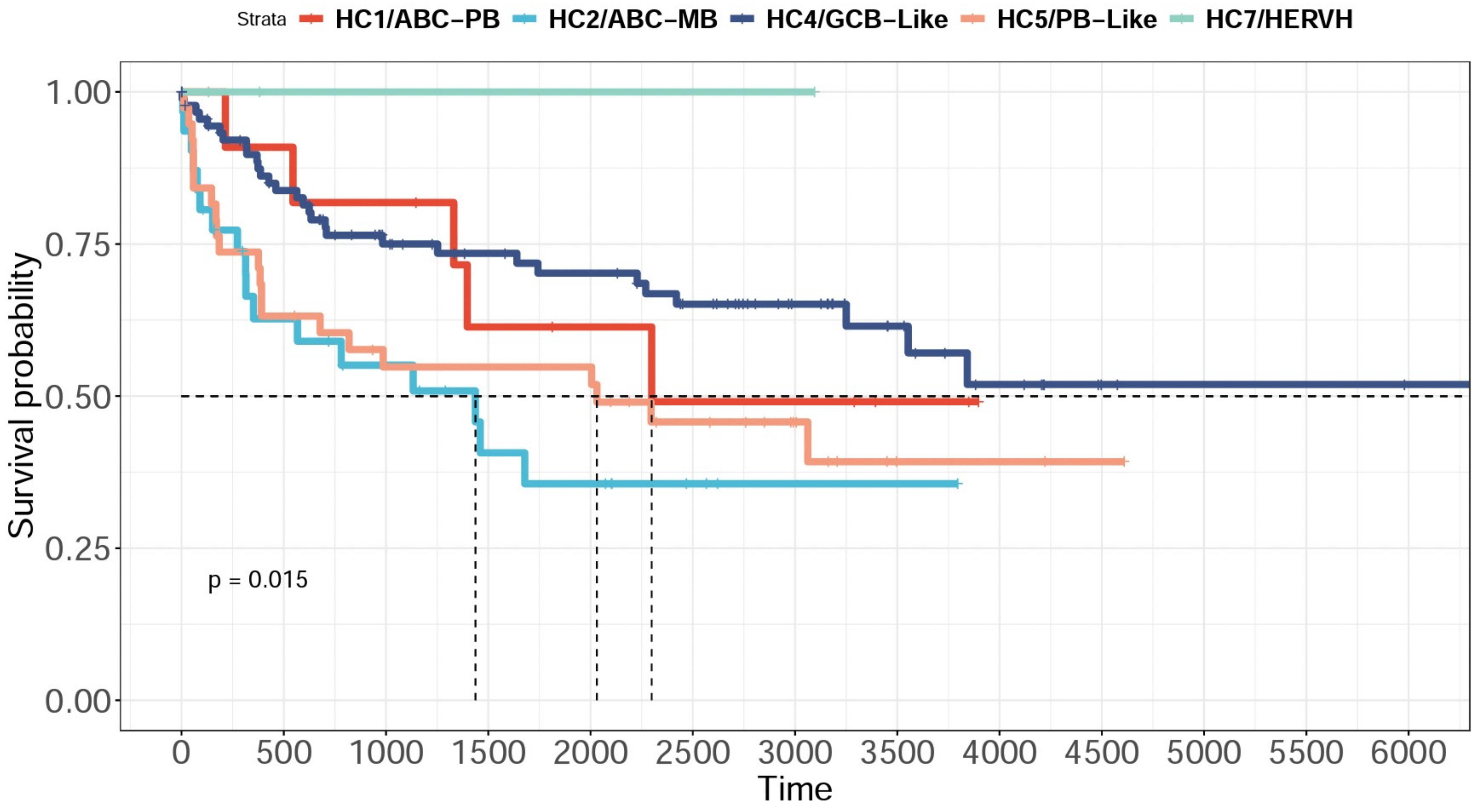

Strata
HC1/ABC−PB
HC2/ABC−MB
HC4/GCB−Like
HC5/PB−Like
HC7/HERVH
Survival probability
1.00
0.75
0.50
0.25
0.00
p = 0.015
0
500
1000
1500
2000
2500
3000
3500
4000
4500
5000
5500
6000
Time

### Supplementary Tables

| | Number of Genes | Number of LINEs | Number of HERVs |
|---|---|---|---|
| **DLBLC (n=529)** | 36,246 | 8,886 | 4,567 |
| **BL (n=113)** | 34,453 | 16,350 | 4,099 |
| **FL (n=12)** | 27,908 | 2,593 | 2,068 |
| **B-AG (n=35)** | 25,629 | 1,520 | 1,118 |
| **B-HM (n=17)** | 23,709 | 1,939 | 1,464 |

**Supplementary Table 1: Number of coding genes, LINEs, and HERV loci remaining in each dataset after filtering**

| Cell Type | baseMean | log2FoldChange | lfcSE | pvalue | padj | locus |
|---|---|---|---|---|---|---|
| BMPC | 10.8887346 | 4.37784863 | 0.33767889 | 1.94E-38 | 2.17E-35 | HML2_20q11.22 |
| BMPC | 46.2145428 | 3.88647483 | 0.30492399 | 3.29E-37 | 1.84E-34 | MER4_9p13.3a |
| BMPC | 20.9878326 | 3.44465987 | 0.27234532 | 1.15E-36 | 4.26E-34 | MER4_9p13.3b |
| BMPC | 44.8270989 | 2.35333027 | 0.19148483 | 1.03E-34 | 2.86E-32 | HERVP71A_15q24.2 |
| BMPC | 6.79069424 | 7.07315797 | 0.66226202 | 1.26E-26 | 2.81E-24 | ERV316A3_15q25.2a |
| BMPC | 9.19435008 | 3.84511792 | 0.3688537 | 1.92E-25 | 3.56E-23 | ERV316A3_5q15a |
| BMPC | 55.7281107 | 2.74673438 | 0.26396941 | 2.34E-25 | 3.73E-23 | ERVLE_4q31.3a |
| BMPC | 36.4028759 | 2.78409297 | 0.27798618 | 1.31E-23 | 1.82E-21 | HERVW_2q24.3 |
| BMPC | 166.558057 | -3.778879 | 0.38213487 | 4.65E-23 | 5.76E-21 | ERVLE_6q22.31b |
| BMPC | 11.2035607 | 7.51598707 | 0.76360265 | 7.36E-23 | 8.21E-21 | HERVL18_7q31.32 |
| DZ | 314.964823 | 2.94489674 | 0.2194209 | 4.54E-41 | 5.06E-38 | HERV3_14q32.33 |
| DZ | 11.0315708 | 4.19079876 | 0.34262119 | 2.11E-34 | 1.18E-31 | HERVFH21_1p36.31 |
| DZ | 69.0711909 | 3.22244667 | 0.29935487 | 5.06E-27 | 1.88E-24 | MER61_3q13.11 |
| DZ | 36.1160058 | 3.24496308 | 0.30403643 | 1.36E-26 | 3.80E-24 | HERVFH21_7q11.21 |
| DZ | 24.5969539 | 3.75589702 | 0.36076565 | 2.21E-25 | 4.93E-23 | HARLEQUIN_19p12b |
| DZ | 10.7288541 | 3.31204897 | 0.33535332 | 5.27E-23 | 9.80E-21 | HERVL_1q23.3a |
| DZ | 7.64190534 | 4.00058561 | 0.41853948 | 1.20E-21 | 1.90E-19 | MER4_2q14.1b |
| DZ | 99.9825746 | 3.81395034 | 0.3995716 | 1.36E-21 | 1.90E-19 | HERVL_12p13.1b |
| DZ | 6.50233386 | 4.54297017 | 0.47952142 | 2.69E-21 | 3.34E-19 | ERV316A3_Xp21.2 |
| DZ | 260.455902 | 1.50529524 | 0.15925132 | 3.31E-21 | 3.69E-19 | HML5_12q23.1 |
| LZ | 112.252844 | 1.75835201 | 0.16973012 | 3.78E-25 | 4.22E-22 | HERVI_15q25.1 |
| LZ | 26.4997086 | 3.21164587 | 0.31634322 | 3.23E-24 | 1.80E-21 | ERVLB4_6p23 |
| LZ | 51.5932719 | 2.17517652 | 0.2288356 | 1.99E-21 | 7.41E-19 | ERVLB4_8p21.3b |
| LZ | 74.7588178 | 2.24408657 | 0.24562613 | 6.47E-20 | 1.80E-17 | MER101_2p25.2 |
| LZ | 198.648314 | 1.63212081 | 0.17962949 | 1.03E-19 | 1.91E-17 | HERVH_2p14b |
| LZ | 314.964823 | 1.99906159 | 0.21988242 | 9.77E-20 | 1.91E-17 | HERV3_14q32.33 |
| LZ | 41.7091526 | 3.78914004 | 0.44762951 | 2.56E-17 | 2.86E-15 | ERV316A3_2q35i |
| LZ | 15.9321312 | 2.64051774 | 0.32523319 | 4.71E-16 | 4.37E-14 | MER41_4q31.1 |
| LZ | 99.9825746 | 3.1939485 | 0.39996917 | 1.40E-15 | 1.20E-13 | HERVL_12p13.1b |
| LZ | 76.5627064 | 1.52153044 | 0.19093247 | 1.60E-15 | 1.27E-13 | HML5_3q26.2 |

| | | | | | | |
|---|---|---|---|---|---|---|
| MB | 25.5376126 | 4.01253318 | 0.44604159 | 2.34E-19 | 2.41E-16 | ERVLB4_14q23.3 |
| MB | 54.6747308 | 2.32886096 | 0.31650359 | 1.87E-13 | 9.60E-11 | HERVL_2p12a |
| MB | 69.0711909 | 2.0987513 | 0.29658573 | 1.48E-12 | 4.28E-10 | MER61_3q13.11 |
| MB | 19.8407038 | 2.43669552 | 0.34513764 | 1.66E-12 | 4.28E-10 | HARLEQUIN_5q33.3 |
| MB | 4.11604419 | 3.50125529 | 0.50611128 | 4.58E-12 | 9.43E-10 | ERVLE_9q21.31d |
| MB | 23.2968619 | 2.45890083 | 0.36513474 | 1.65E-11 | 2.83E-09 | MER61_19p12c |
| MB | 26.8514584 | 2.19928295 | 0.32864405 | 2.20E-11 | 3.24E-09 | HERVFRD_2p12a |
| MB | 24.5969539 | 2.31764342 | 0.36077492 | 1.33E-10 | 1.52E-08 | HARLEQUIN_19p12b |
| MB | 68.1825026 | 1.69218373 | 0.2679894 | 2.71E-10 | 2.54E-08 | HML2_1q22 |
| MB | 16.0936966 | 2.17570587 | 0.37409464 | 6.03E-09 | 4.77E-07 | PABLB_7q11.21 |
| NB | 260.161745 | -5.3430997 | 0.48496785 | 3.15E-28 | 3.51E-25 | HERVL18_11q14.2b |
| NB | 68.1825026 | 3.36527491 | 0.31318367 | 6.23E-27 | 3.47E-24 | HML2_1q22 |
| NB | 54.6747308 | 3.62913079 | 0.34056663 | 1.63E-26 | 6.07E-24 | HERVL_2p12a |
| NB | 26.8514584 | 3.52390742 | 0.35350931 | 2.10E-23 | 5.84E-21 | HERVFRD_2p12a |
| NB | 198.648314 | -2.1717859 | 0.22052682 | 6.98E-23 | 1.56E-20 | HERVH_2p14b |
| NB | 97.3158729 | -1.8246397 | 0.20821681 | 1.90E-18 | 3.53E-16 | HERVH_15q26.3b |
| NB | 102.48819 | -2.8722752 | 0.33270862 | 5.98E-18 | 9.52E-16 | HERVP71A_3p26.1 |
| NB | 102.36481 | -2.0725408 | 0.24705674 | 4.91E-17 | 6.84E-15 | ERVLE_2p14b |
| NB | 123.700933 | 2.16855566 | 0.2630104 | 1.65E-16 | 2.04E-14 | HERV9_6q25.1 |
| NB | 47.5824101 | 1.77030829 | 0.21806237 | 4.73E-16 | 5.27E-14 | HERVK11_1q23.3a |
| PB | 160.78374 | 3.08148225 | 0.17442372 | 7.58E-70 | 8.45E-67 | HML3_1q25.2 |
| PB | 18.6471599 | 4.95495287 | 0.33177381 | 1.96E-50 | 1.09E-47 | HERVP71A_8q24.13 |
| PB | 102.48819 | 3.98135965 | 0.28015381 | 7.79E-46 | 2.90E-43 | HERVP71A_3p26.1 |
| PB | 74.2730714 | 2.85271936 | 0.22421539 | 4.40E-37 | 1.23E-34 | HML2_1q23.3 |
| PB | 44.8270989 | 1.88966056 | 0.15791043 | 5.31E-33 | 1.18E-30 | HERVP71A_15q24.2 |
| PB | 18.9632769 | 4.81596924 | 0.43653806 | 2.67E-28 | 4.97E-26 | HML3_8q24.13 |
| PB | 7.1632018 | 5.51179855 | 0.5095672 | 2.87E-27 | 4.57E-25 | ERVLB4_15q13.3a |
| PB | 102.36481 | 2.25871145 | 0.22286633 | 3.87E-24 | 5.39E-22 | ERVLE_2p14b |
| PB | 13.8075309 | 4.28984401 | 0.43258816 | 3.52E-23 | 4.36E-21 | LTR25_3q24a |
| PB | 194.639084 | -2.1472848 | 0.21704713 | 4.46E-23 | 4.97E-21 | ERVL_16p12.2 |

**Supplementary Table 2: Top 10 differentially expressed HERVs in each B-cell subtype in the B-AG dataset**

| **DZ** | **LZ** | **PB** | **BMPC** | **NB** | **MB** |
|---|---|---|---|---|---|
| NOL9 | TMEM131L | CALCOCO1 | SPTBN2 | CD72 | SESN3 |
| BCL7A | CD22 | RP11-16E12.2 | SCARB2 | COL19A1 | PARP15 |
| DCK | SWAP70 | FBH1 | SPAG4 | MACF1 | L1FLnI_3q13.2a |
| TMEM131L | LCP1 | SLC17A9 | FOXO3 | TREML2 | FCRL4 |

| | | | | | |
|---|---|---|---|---|---|
| DPY19L2P2 | RFTN1 | WDR45 | ABCB9 | SMAD3 | L1FLnI_9q21.31k |
| ABR | MYO1E | CNKSR1 | AMPD1 | PARP15 | CD96 |
| POLH | BCL7A | AF127936.3 | BMP6 | TRIM22 | NLRC5 |
| BACH2 | EBF1 | TMEM39A | TLCD4 | CR2 | BANK1 |
| MFHAS1 | LCK | SPAG4 | SCNN1B | SESN3 | MCOLN2 |
| GCSAM | PTK2 | MCEE | ITGA8 | EML4 | TTC39C |
| DNASE1 | IRAG2 | IGHV4-61 | BTD | CD47 | CENPK |
| SGO1 | LMO2 | LINC02576 | FXN | BANK1 | DEK |
| NCAPD2 | SORL1 | RP11-114N1.1 | FBXW7 | L3MBTL3 | MS4A1 |
| CTB-193M12.5 | CD72 | PLPP5 | MEI1 | MGAT5 | PIK3R6 |
| HES6 | HIVEP3 | SSR4 | ISCU | FMNL3 | TMEM273 |
| FANCD2 | BCL6 | GMPPA | MOXD1 | CLEC17A | FCMR |
| TMPO-AS1 | MCOLN2 | TNFRSF17 | ANKRD28 | FCMR | CCDC141 |
| TRIM59 | IRF8 | HML3_1q25.2 | MTCYBP41 | TRAF3IP3 | ZBTB32 |
| HAUS8 | MS4A1 | SIL1 | TBCEL | DTX3L | NAPSB |
| DTX1 | WEE1 | SELENOM | RP11-720D4.2 | KCNG1 | SPIB |
| SLC6A6 | LYSMD2 | RUSF1 | FER1L4 | ABCB4 | ITGAM |
| SLC9A7 | PAX5 | IGKV3-11 | L1FLnI_5q21.3u | RP11-421L21.3 | EPHA4 |
| FAM241A | MARCKSL1 | MLKL | HDLBP | PARP9 | CDK5R1 |
| RNF19B | ZC3H12D | CTD-2227E11.1 | RNF207 | RP11-429O22.1 | WDR76 |
| IL4R | BICDL1 | FICD | KCNH2 | GPR174 | TRIM22 |
| SSBP2 | TMED8 | MBNL2 | FICD | L1FLnI_3q21.1c | CD1C |
| CRACD | TMOD2 | PPP1R3B | CCPG1 | PDE7B | CCR1 |
| RCCD1 | GCNT2 | BET1L | CRELD2 | LINC02397 | TESPA1 |
| CDK19 | PIK3AP1 | MTMR9LP | LMNA | VAV3 | SYNPO |
| RUBCNL | DCK | DQX1 | EPHA10 | S1PR1 | CDCA7 |
| SLX4 | ANKRD33B | NOL3 | MINAR1 | GAPT | FAM111B |
| IRF8 | SYT11 | CD79A | DENND2C | PPFIBP1 | SIGLEC6 |
| C21orf58 | SYPL1 | DENND6B | SEL1L | LPGAT1 | BHLHE40-AS1 |
| REXO5 | GPR18 | C1GALT1C1 | CTSF | LINC00926 | UST |
| RRM2B | HLA-DMB | IGKV3-15 | PPFIA3 | MYO7B | ZBED2 |
| SGPP1 | HTR3A | ATF6 | REPS2 | PLEKHA1 | CSF1 |
| ARHGEF39 | GCNT1 | SELENOS | RP11-44D5.1 | TRAF5 | GINS2 |
| CRYBG1 | CR2 | BTG2 | H1-10-AS1 | STEAP1B | NCR3 |
| JPT1 | VNN2 | STARD5 | MDK | UST | L1FLnI_9q21.31i |
| KNSTRN | PANK1 | MZB1 | L1FLnI_10p13e | ABCB1 | ITGAX |
| TACC3 | CLEC17A | QPCTL | OS9 | SH3BP2 | WDHD1 |

| | | | | | |
|---|---|---|---|---|---|
| RAD21 | PUS10 | FNDC3A | CCR2 | MMP17 | MYO1F |
| HMGN2 | ARHGAP17 | OSBPL3 | CYFIP1 | LAIR1 | DHFR |
| TCL1A | ALOX5AP | IFNG-AS1 | ACADVL | PCDH9 | AHNAK2 |
| APBB2 | LHFPL2 | TMC3-AS1 | ASS1 | C1orf162 | TRAC |
| SUGCT | SLC16A2 | ZBP1 | NOL3 | RP11-861A13.2 | BRIP1 |
| RBBP7 | WDR76 | ST6GAL1 | SIL1 | PARP14 | CNR2 |
| CIT | RGS8 | ERN1 | ZCCHC24 | HPSE | SCML4 |
| RASAL1 | STAT6 | IGKC | GRIK4 | CRTC3 | EPHB6 |
| RACGAP1 | MAML3 | FBXW7 | PDK1 | FAM117B | TNR |
| MTA3 | SERPINA9 | PLD3 | EIF2AK4 | TMC8 | CLNK |
| CKAP5 | IL4R | LINC02352 | INPP4A | DDX60L | BICDL1 |
| RFTN1 | P2RY12 | CRELD2 | GLDC | LBH | RP1-47M23.3 |
| BCL6 | BRI3BP | SERP1 | CDC14B | SP110 | H1-1 |
| WEE1 | PHF6 | ACADVL | RAPGEF2 | SHISAL2A | MCM4 |
| CNTROB | PRAMENP | IGKV2-24 | MB21D2 | SCN3A | L1FLnI_9q21.31h |
| RMI2 | MED12L | ARFGAP3 | KIF13B | CEPT1 | CCR7 |
| BRI3BP | TMEM159 | CFAP54 | PLPP5 | RASGRP2 | NPAP1P4 |
| KANK2 | SPRED2 | SLAMF7 | LTK | FGR | TNFRSF1B |
| TMPO | FAM81A | FBXO16 | SLC6A9 | AC002480.4 | CDC45 |
| NDC1 | LTA | APOL2 | C11orf80 | ITGA4 | RP11-712B9.2 |
| CFAP251 | LAT2 | EIF2AK4 | BMI1 | ARMH1 | ERVLB4_14q23.3 |
| TERF2 | CD80 | RWDD2A | LDLR | RP11-564A8.4 | BHLHE40 |
| CENPK | RP11-203B7.2 | OGT | RIPOR3 | SELL | CELF2-AS1 |
| KIF22 | SGPP1 | TXNDC11 | CALCOCO1 | BTLA | CENPI |
| MCUB | SPIB | PPCDC | IGHV4-61 | L1FLnI_6q13i | SELL |
| FAM83D | PHLPP1 | ADA2 | P3H3 | DOP1B | PIK3R5 |
| SLC35E3 | RAPGEF5 | AC026202.3 | C12orf73 | NIBAN3 | FGR |
| TIMELESS | SH3RF1 | IGHV4-28 | SLC17A9 | SPRY1 | FAM81A |
| KDM1B | CENPK | IGLV3-21 | L1FLnI_15q25.2a | L1FLnI_3q13.2a | FUT7 |
| HMGB2 | AFF2 | MEI1 | SAR1B | C12orf42 | GPR34 |
| ZNF831 | CTD-2325P2.3 | HERVP71A_8q24.13 | TPM4 | MACROD2 | L1FLI_6p22.3 |
| CENPI | RMI2 | DENND2C | PLOD3 | DENND11 | MCM6 |
| DEK | PAG1 | SEL1L | PHLPP2 | TGFBR2 | E2F8 |
| NCAPG2 | HLA-DOA | ACP2 | PLEKHN1 | RBMS1 | RMI2 |
| GIHCG | NAPSB | IGLV1-47 | RRAGD | MS4A1 | XRCC2 |
| PEX5 | GALNT14 | HM13 | KIF19 | HVCN1 | ARAP2 |
| AICDA | MYBL1 | GMPPB | HOMER3 | FCER2 | GAPT |

| | | | | | |
|---|---|---|---|---|---|
| WDR76 | TESPA1 | GALK2 | MYO5B | DNAH11 | KYNU |
| UBE2C | BPNT1 | CPEB4 | CFLAR | ESAM | MCM2 |
| PEG10 | SAMD15 | IGKV3-20 | FNDC3A | MAML2 | RRM2 |
| CCNB2 | CNR2 | NKX6-3 | ERN1 | IFNGR1 | MCM10 |
| STMN1 | SIGLEC10 | CHPF | SLC41A2 | ZBTB37 | RP11-403N16.3 |
| PTTG1 | ACTN2 | REXO2 | MYO5C | RP11-564A8.8 | SLC37A2 |
| MYO1E | FEZ1 | MANEA | IGHV3OR16-13 | L1FLnI_5q11.2ta | ORC1 |
| SMARCA4 | L1FLnI_4q31.22d | ALG2 | IGHV3OR16-8 | P2RY14 | NT5C3AP2 |
| TROAP | FGD6 | FER1L4 | PPCDC | GCNT1 | BRCA1 |
| ZNF106 | CAMK1 | SAR1B | SLC38A4 | GVINP1 | EXO1 |
| CENPO | LPP-AS2 | SEC61A1 | IGHV3OR16-17 | SLC38A11 | RAB31 |
| LBR | SGO1 | IGKV1-16 | ARMC2 | MTSS1 | DSCC1 |
| CDCA7 | ASAP3 | IGHV4-59 | FAM13A | CHML | CDCA5 |
| CKAP2 | L1FLnI_14q32.13a | MINAR1 | ABHD2 | CD22 | EML4-AS1 |
| SLC2A5 | BRIP1 | CTD-2240J17.5 | MYO1D | RP11-452F19.4 | LTB |
| PSRC1 | IL7 | IGHV1-18 | RPS27AP8 | SNX18 | CAPG |
| LINC01991 | ZNF608 | LINC00698 | SELENOM | CR1 | AC008697.1 |
| PTK2 | LINC02099 | TMEM214 | FBH1 | CELF2-AS1 | L1CAM |
| EZR | DEF8 | CCPG1 | ITPRIP | CNR2 | H2BC13 |
| KIF20A | FCRL3 | FKBP11 | PSAP | ZNF528-AS1 | ASF1B |
| CCNB1 | LPP | FBXL8 | PIP5KL1 | GBP4 | CTD-2509G16.5 |
| SPDL1 | DMD | CHPF2 | C16orf54 | MARCHF1 | RASGRP2 |
| PAX5 | HOPX | L1FLnI_3q23t | RAB3D | IL24 | ESCO2 |
| HMGN2P46 | CDCA7 | RP1-134E15.3 | IGHV3OR16-9 | HML2_1q22 | ADAMTS6 |
| SLC30A4 | VAV3 | ARMCX3 | GOLGA2 | RP11-281P23.3 | H2BC9 |
| STIL | REL | PDIK1L | RAB36 | FAM177B | TK1 |
| NUF2 | WDHD1 | ACOXL | LINC02711 | SP100 | LINC01991 |
| TPX2 | DEK | GLRX | VPS37B | CTNND1 | GPR82 |
| CKS1B | MAST2 | HERVP71A_3p26.1 | CSGALNACT1 | PAXIP1-AS2 | TNFSF12 |
| MYBL1 | U62631.5 | DERL3 | FAM114A1 | CD1A | LFNG |
| DBF4B | B3GALNT1 | ARF4 | ATP8B2 | RP3-323N1.2 | CD80 |
| CTPS2 | FCRLA | FNDC3B | FNDC3B | LIX1-AS1 | POC1A |
| DNMT3B | HLA-DRA | EDEM2 | LYPD6B | CASP4LP | AICDA |
| KIF18B | L1FLnI_15q22.2a | MINDY1 | BMP8B | ZNF528 | CCNA2 |
| POU4F1 | CAMK2B | PGM3 | TTLL7 | ZBTB16 | CCNB1 |

| | | | | | |
|---|---|---|---|---|---|
| ARL6IP1 | KCNMB4 | RP11-294C11.2 | PERP | SATB1-AS1 | CR1 |
| ESPL1 | EPS15-AS1 | IGKV1-12 | NPC2 | DDX60 | CCR5 |
| FOXM1 | CDK5R1 | RPN2 | IGF1 | ARHGAP15 | OSTN-AS1 |
| C1orf112 | SNX29P1 | IGHV3-21 | TXNDC11 | JAM3 | RP11-403N16.4 |
| SYBU | HELLS | NXPE3 | CTD-2653D5.1 | CD1C | CXCR3 |
| FBXO43 | SLC1A1 | RAB3D | CLPTM1L | BEND5 | TTC24 |
| ERCC6L | RP13-786C16.1 | MIR5571 | PECAM1 | ABAT | DTL |
| WDR62 | RP11-231C14.7 | IGHV3-30 | RAPGEF3 | DEXI | TNFRSF13B |
| NDC80 | AC023590.1 | XBP1 | RDX | MX2 | CLSPN |
| DEPDC1B | AC073043.1 | IGHV4-39 | HEXB | HHEX | TGM2 |
| NEIL1 | BCL2A1 | METTL7A | GPR176 | ZCCHC18 | RP3-323N1.2 |
| TRAF5 | L1FLnI_4q13.3j | AFF1 | CASP10 | AC104530.1 | AHNAK |
| ALPK1 | LINC01991 | LARP1B | PRDX4 | RHOBTB1 | CIITA |
| CDKN3 | RP11-415F23.5 | L1FLnI_5q33.3l | FAM174A | HS3ST1 | PREX1 |
| AURKB | LRRC32 | PLEKHN1 | UAP1 | CCDC141 | SCIMP |
| CENPN | TCL1A | IGHJ5 | HSP90B1 | PRICKLE1 | PCLAF |
| DNMT1 | TMEM229B | C1R | RP11-665E10.2 | L1FLnI_1q32.2e | DDX60L |
| CENPH | CFAP20DC | TXNDC15 | ARID3B | PEAK1 | CDC20 |
| KIFC1 | PXDN | PRDX4 | TMEM59 | ARAP2 | CALHM2 |
| BCAS4 | RPRD1B | CLIP4 | RBM47 | L1FLnI_4q32.2i | DMC1 |
| KIF2C | LDHAL6B | ZCCHC24 | CD63 | TPK1 | E2F1 |
| GTSE1 | NLRP4 | JCHAIN | SELENOS | L1FLnI_11q12.1v | AURKB |
| RAD51AP1 | ANKLE1 | IGKV4-1 | ELL2 | MOB3B | H2BC10 |
| CENPL | LINC02137 | SRPRB | FKBP11 | EML4-AS1 | NPAP1P6 |
| CDC25C | ADARB1 | ERVLE_2p14b | WDR45 | RP11-35G9.3 | RIN3 |
| AFF2 | GPR137B | TMC3 | SLAMF7 | PTPRK | PLAC4 |
| CCNA2 | DLGAP1 | LINC02227 | WIPI1 | IFNK | ZWINT |
| CDK1 | PCDHGC4 | RBM47 | TCN2 | P2RY10 | CD247 |
| CDCA2 | RP11-131H24.4 | LINC01485 | FGFRL1 | BCL11A | KIFC1 |
| NEIL3 | UGT8 | PREB | UBE2QL1 | GPR65 | SERPINB6 |
| DHFR | KIF5C | IL10RA | JSRP1 | HLA-DMB | H1-5 |
| HJURP | SCIMP | IGHV3OR16-9 | SCAMP5 | BTBD6P1 | H4C13 |
| FAM81A | CIITA | PIM2 | B9D1 | TTC24 | PBK |
| NSD2 | E2F8 | PDIA4 | ZNF275 | HLA-DPB1 | CCL22 |
| DCP2 | EBI3 | IGKV3D-11 | SEPTIN10 | HLA-DOA | STMN1 |
| TTK | LINC01857 | RP11-490O6.2 | BCL2 | ST3GAL1 | CHAF1B |

| | | | | | |
|---|---|---|---|---|---|
| IQGAP3 | RGS13 | B9D1 | TMEM63B | C8orf37 | ACP5 |
| HERV3_14q32.33 | HERVI_15q25.1 | HML3_1q25.2 | HML2_20q11.22 | HML2_1q22 | ERVLB4_14q23.3 |
| HERVFH21_1p36.31 | ERVLB4_6p23 | HERVP71A_8q24.13 | MER4_9p13.3a | HERVL_2p12a | HERVL_2p12a |
| MER61_3q13.11 | ERVLB4_8p21.3b | HERVP71A_3p26.1 | MER4_9p13.3b | HERVFRD_2p12a | MER61_3q13.11 |
| HERVFH21_7q11.21 | MER101_2p25.2 | HML2_1q23.3 | HERVP71A_15q24.2 | HERV9_6q25.1 | HARLEQUIN_5q33.3 |
| HARLEQUIN_19p12b | HERVH_2p14b | HERVP71A_15q24.2 | ERV316A3_15q25.2a | HERVK11_1q23.3a | ERVLE_9q21.31d |
| HERVL_1q23.3a | HERV3_14q32.33 | HML3_8q24.13 | ERV316A3_5q15a | PRIMA41_Yq11.223a | MER61_19p12c |
| MER4_2q14.1b | ERV316A3_2q35i | ERVLB4_15q13.3a | ERVLE_4q31.3a | HML5_Yq11.223e | HERVFRD_2p12a |
| HERVL_12p13.1b | MER41_4q31.1 | ERVLE_2p14b | HERVW_2q24.3 | HUERSP2_6p22.3 | HARLEQUIN_19p12b |
| ERV316A3_Xp21.2 | HERVL_12p13.1b | LTR25_3q24a | HERVL18_7q31.32 | MER4B_20q13.12 | HML2_1q22 |
| HML5_12q23.1 | HML5_3q26.2 | HERVL40_11q13.4b | MER101_19q13.2c | PRIMA41_Yq11.223b | PABLB_7q11.21 |
| HERVL66_19p12f | HERVE_11q13.4c | HARLEQUIN_11q13.4 | HERVH_9p13.3b | ERV316A3_6q24.1a | HERVL_5q12.3 |
| HML5_Xq11.2a | HUERSP2_22q11.22 | MER4_22q12.3 | HERVP71A_8q24.13 | HERVIP10FH_2p14 | HERVH_9q21.31a |
| ERVLB4_8p21.3b | HERVL18_11q22.3 | HERVL40_19q13.33 | HERVL18_11q14.2b | ERVLB4_6p23 | MER4_19p12a |
| MER61_19p12c | HERVL_3q13.11c | HUERSP2_19q13.2 | ERV316A3_17q24.3a | HERVK11_1q23.3b | ERV316A3_12q24.13 |
| HERVIP10F_11q24.2 | MER4_19p12a | HERVFH21_Xq11.2a | ERVLE_5q21.3b | ERVLE_5q31.1a | MER4_2p11.2a |
| HERVIP10F_2q21.2 | MER61_19p12c | ERV316A3_3q24a | ERV316A3_8q13.3a | HERVH_19p13.2a | HERVL18_1q32.2 |
| MER101_3q26.31 | HERVEA_6q22.31 | ERVLB4_16q13b | PABLB_2q31.1 | MER4_7q21.12 | HERVH_12q13.2b |
| MER4_19p12a | HERVL_17p11.2b | MER61_3q24a | HERVL_6p25.2 | ERVL_Xq21.1a | ERVLE_8q21.13g |
| HERV3_1q23.3 | HML2_4p16.3a | ERVLE_16p11.2b | HERV9_10p13 | HERV3_16p13.3 | MER34B_1q23.3b |
| ERV316A3_1p34.3c | MER61_3q13.11 | HERVFH21_Xq11.2b | HML6_14q24.2 | ERV316A3_12q24.13 | HERVL_12p13.1b |
| HERVIP10FH_17q21.32 | MER101_2p16.3 | ERVLB4_3q12.3 | ERVLE_18q11.2c | MER4_2p11.2a | HERV3_19p13.3 |
| MER4_8p11.1b | HUERSP2_7q35 | HERV9_10p13 | ERV316A3_2p25.1a | ERV316A3_10q23.33 | MER41_1q44a |
| ERVLB4_3q25.2c | HARLEQUIN_10q23.1 | PABLB_2q31.1 | ERVLE_8q24.22g | ERVLE_8q21.13g | ERVLE_15q15.1 |
| ERVL_6q15 | HERVL18_1q32.2 | MER61_12q13.12 | HML1_6q23.2 | ERVLE_4q21.23a | HERV3_4p16.1 |
| HUERSP3_8p11.1b | HARLEQUIN_19p12b | HERVIP10FH_19q13.43 | HERVP71A_2q32.2 | ERV316A3_21q21.2g | ERVLE_4q21.23a |

**Supplementary Table 3: Top genes and HERVs from the B-AG dataset used to create B-cell-specific sets for HAGSEAS analysis**
